# Hemocyte Differentiation to the Megacyte Lineage Enhances Mosquito Immunity Against *Plasmodium*

**DOI:** 10.1101/2021.12.24.474101

**Authors:** Ana Beatriz Barletta Ferreira, Banhisikha Saha, Nathanie Trisnadi, Octavio Talyuli, Gianmarco Raddi, Carolina Barillas-Mury

**Author notes:** Correspondence should be addressed to (CB-M). Nathanie Trisnadi, Atropos Therapeutics Inc., San Carlos, California, USA. Gianmarco Raddi, School of Clinical Medicine, University of Cambridge, Cambridge CB2 0SP, UK, CRUK Cambridge Institute, Cambridge CB2 0RE, UK.

## Abstract

Silencing Cactus, a suppressor of Toll signaling, in *Anopheles gambiae*, eliminates *Plasmodium* ookinetes by enhancing local release of hemocyte-derived microvesicles that promote activation of the mosquito complement-like system. We report that Cactus silencing dramatically increases the proportion of megacytes, a new effector hemocyte subpopulation of large granulocytes, from 5 to 79% of circulating granulocytes. Transcriptomic and morphological analysis, as well as cell counts, *in situ* hybridization and expression of cell-specific markers, indicate that Cactus silencing triggers granulocyte differentiation into megacytes, a process mediated by the Rel1 transcription factor of the *Toll* pathway. Megacytes are very plastic cells that can extend long filopodia, tend to form clusters *in vivo*, and are massively recruited to the basal midgut surface in response to bacterial feeding and *Plasmodium* infection. We show that hemocyte differentiation to the megacyte lineage greatly enhances mosquito immunity against *Plasmodium*.

## Introduction

Ookinete traversal of the *Anopheles gambiae* midgut disrupts the barriers that normally prevent bacteria of the gut microbiota from coming in direct contact with epithelial cells (Kumar et al., 2010), and this attracts hemocytes to the basal surface of the midgut (Barletta et al., 2019). *Plasmodium* ookinetes also cause irreversible damage to the cells they invade and trigger a strong caspase-mediated nitration response (Han et al., 2000, Oliveira Gde et al., 2012, Trisnadi and Barillas-Mury, 2020). When hemocytes come in contact with a nitrated midgut surface, they undergo apoptosis and release hemocyte-derived microvesicles (HdMv) (Castillo et al., 2017). Local HdMv release promotes activation of thioester containing-protein 1 (TEP1) (Castillo et al., 2017), a major final effector of the mosquito complement-like system that binds to the parasite’s surface and forms a complex that lyses the ookinete (Blandin et al., 2004).

Mosquito hemocytes are classified into three cell types, prohemocytes, oenocytoids and granulocytes, based on their morphology. However, single cell RNA sequencing (sc-RNAseq) analysis of *An. gambiae* hemocytes identified several novel subpopulations of granulocytes based on their transcriptional profiles, and defined molecular markers specific for hemocyte subpopulations (Raddi et al., 2020). Furthermore, Lineage analysis revealed that regular granulocytes derive from prohemocytes and can further differentiate into distinct cell types, including dividing granulocytes, and two final effector cells, megacytes and antimicrobial (AM) granulocytes (Raddi et al., 2020).

Silencing *Cactus*, a negative regulator of *Toll* signaling in *A. gambiae* mosquitoes, elicits a very strong TEP1-mediated immune response that eliminates *Plasmodium berghei* ookinetes (Frolet et al., 2006). This phenotype can be rescued by co-silencing Cactus with either TEP1 or the *Rel1* transcription factor, indicating that parasite elimination is mediated by activation of *Toll* signaling, with TEP1 as a final effector (Frolet et al., 2006). Later studies showed that hemocytes mediate this enhanced immune response, as transfer of *Cactus*-silenced hemocytes into naïve mosquitoes recapitulates the phenotype of systemic *Cactus* silencing (Ramirez et al., 2014). Furthermore, cactus silencing also increases HdMv release in response to ookinete midgut invasion (Castillo et al., 2017), indicating that hemocytes are more reactive to *Plasmodium* infection. However, the nature of the functional changes in *Cactus*-silenced hemocytes that enhance immunity against *Plasmodium* are not known. Here, we explore the effect of Cactus silencing on circulating hemocyte populations and their response to infection of mosquitoes with bacteria and *Plasmodium*.

## Results

### Effect of Cactus-silencing on mRNA markers of granulocyte populations

The effect of silencing Cactus, a suppressor of Toll signaling, on hemocyte differentiation was explored. Hemocytes that adhere to glass (mostly granulocytes) or that remain in suspension (mostly prohemocytes and oenocytoids) were collected 4 days post-injection from *dsLacZ* control and *dsCactus*-injected females. Bulk sequencing of cDNA libraries generated between 16.2 and 25.3 million fragments that mapped to the *Anopheles gambiae* AgamP4.9 transcriptome. Only transcripts with 10 or more reads were included in the analysis, resulting in a total of 9,421 unique transcripts (https://www.ebi.ac.uk/biostudies/arrayexpress/studies/E-MTAB-11252). Glass-bound and unbound hemocyte samples were analyzed together, because the differences in expression between ds*LacZ* vs. ds*Cactus*-silenced hemocytes explained 81% of the variance between the four experimental groups (Fig. S1A and S1B). Differential expression (DE) analysis of *Cactus*-silenced hemocytes using the DESeq2 software identified 1071 differentially expressed genes (Q-value < 0.001), of which 407 were upregulated (log2 fold change >2), while 664 were downregulated (log2 fold change < −2) (Fig 1A).

**Fig. 1:**
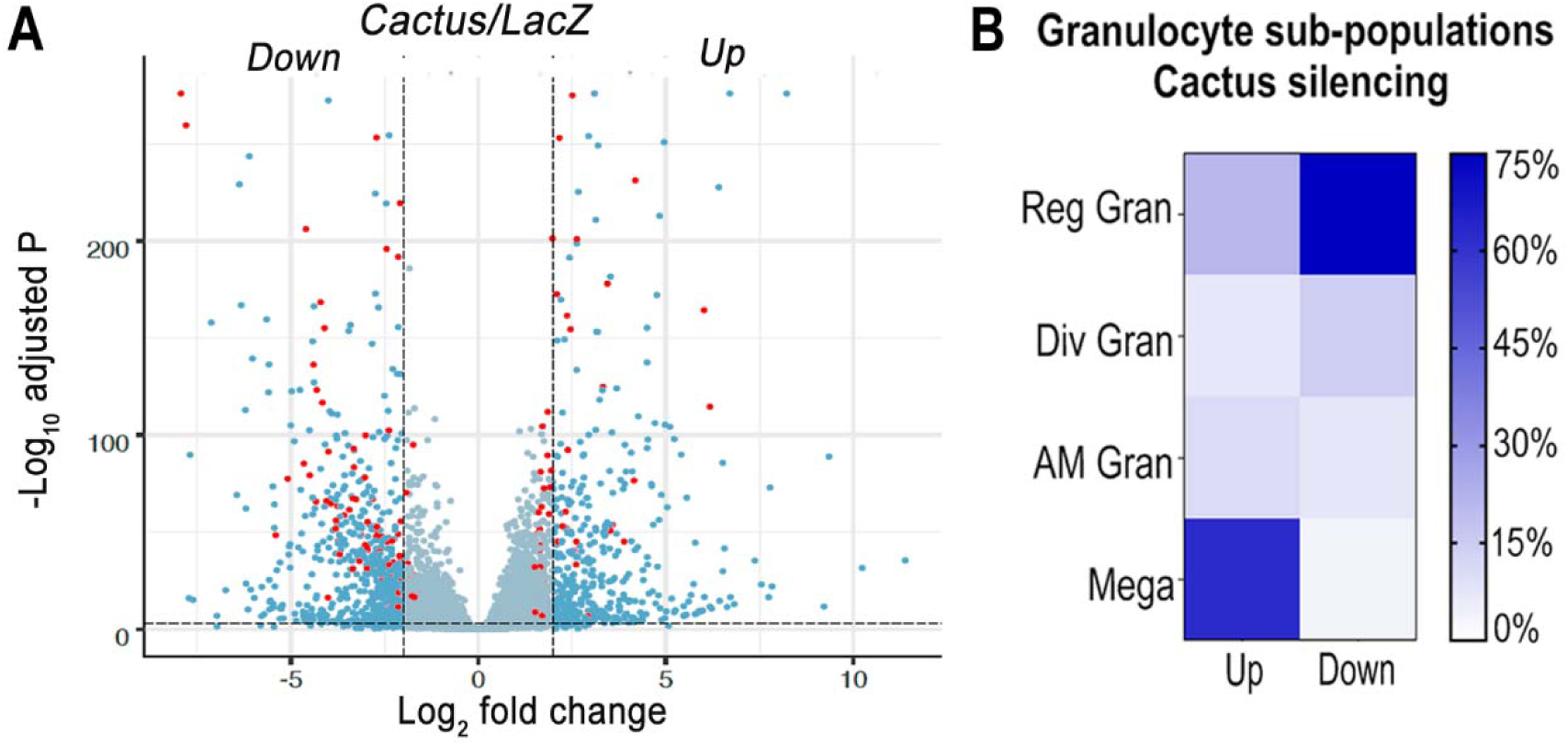
Effect of *Toll* pathway activation on mRNA markers of granulocyte populations. (A) Differential expression of Cactus dsRNA knockdown. From a total of 9421 filtered genes. Volcano plot of DE genes in Cactus silenced hemocytes compared to LacZ control filtered for log2 fold change > 2 and Q-value <0.001. Dark blue dots on the right represent upregulated DE genes and on the left the downregulated ones. Red dots show genes that are hemocytes specific markers. Complete list of up and down regulated genes is listed in Tables S1 and S2. (B) Percentage of granulocyte sub-population markers up and downregulated in Cactus silenced hemocytes. Complete list of up and down regulated genes for each hemocyte subpopulation is in Tables S1 and S2.

The effect of Cactus silencing on expression of the transcripts that define the different hemocyte clusters established by (sc-RNAseq) (Raddi et al., 2020) was analyzed (Tables S1 and S2), to establish whether there was a significant effect on the relative abundance of specific hemocyte subpopulations. Overall, 23 oenocytoid markers, 2 from prohemocytes and 57 from granulocytes were differentially expressed between dsLacZ and dsCactus hemocytes (Tables S1 and S2). Most differentially expressed oenocytoid markers 22/23 (95%) were down-regulated, while one of the prohemocyte markers was up-regulated and the other one was down-regulated (Fig. 1B). The number of down-regulated granulocyte markers 28/57 (49%) was very similar to that of up-regulated ones 29/57 (51%). However, detailed analysis of granulocyte subpopulations revealed that most up-regulated markers 18/29 (62%) correspond to megacytes, while most down-regulated markers correspond to regular granulocytes 21/28 (75%). This suggests that d*sCactus* silencing increases the proportion of circulating megacytes, at the expense of a reduction in regular granulocytes. As expected, expression of several genes involved in *Toll* signaling or final effector this pathway, such as Toll-like receptors, CLIP proteases, Serpins, C-type Lectins and Defensin are upregulated in *dsCactus* hemocytes (Table S3).

### Cactus silencing promotes granulocyte differentiation into megacytes

Cactus silencing did not significantly increase the proportion of total circulating granulocytes, based on hemocyte counts by light microscopy (Fig. S2), suggesting that the observed enhanced immune response could be due to functional changes in hemocytes. The morphology of hemocytes perfused from *Cactus*-silenced females was analyzed using fluorescent probes to stain the actin cytoskeleton and the nucleus. *Cactus* silencing dramatically increased the proportion of large granulocytes (diameter > 40 µm), presumably megacytes, from 5.3% to 79.2% (p<0.0001, X^2^ test) (Fig. 2A and B), in agreement with the observed increase in up-regulated megacyte-specific markers in the transcriptomic analysis of Cactus-silenced hemocytes (Fig. 1B). Interestingly, megacytes from *Cactus*-silenced mosquitoes (Fig. 2A) are even larger (average diameter of 47 µm after spreading in a glass surface) than megacytes from dsLacZ controls (average diameter of 30 µm) (Figure 2C and D). *In situ* RNA hybridization of dsCactus granulocytes with a fluorescent probe for the megacyte-specific marker TM7318, confirmed that the proportion of TM7318-positive granulocytes was much higher (80%) in Cactus-silenced females than in dsLacZ controls (4%) (p<0.0001, X^2^ test) (Fig. 2E and F), providing direct evidence that overactivation of *Toll* signaling triggers a dramatic increase in the proportion of circulating megacytes. Expression analysis of the TM7318 marker in perfused hemocyte samples confirmed that mRNA levels were 42-fold higher in dsCactus hemocytes than the dsLacZ control group (p<0.001, T-test) (Fig. 2G), while a modest increase (2.8-fold) in FBN50 mRNA, a marker of antimicrobial (AM) effector granulocytes, was observed (p<0.0001, T-test). Conversely, expression of FBN11228, a marker of regular granulocytes, decreased by 30-fold in circulating hemocytes of Cactus-silenced mosquitoes (Fig. 2G). The changes in the relative abundance of mRNAs from cell-specific markers in *dsCactus*-hemocytes agrees with the observed changes in hemocyte morphology and the *in situ* hybridization and transcriptomic data (Fig. 2G).

**Fig. 2:**
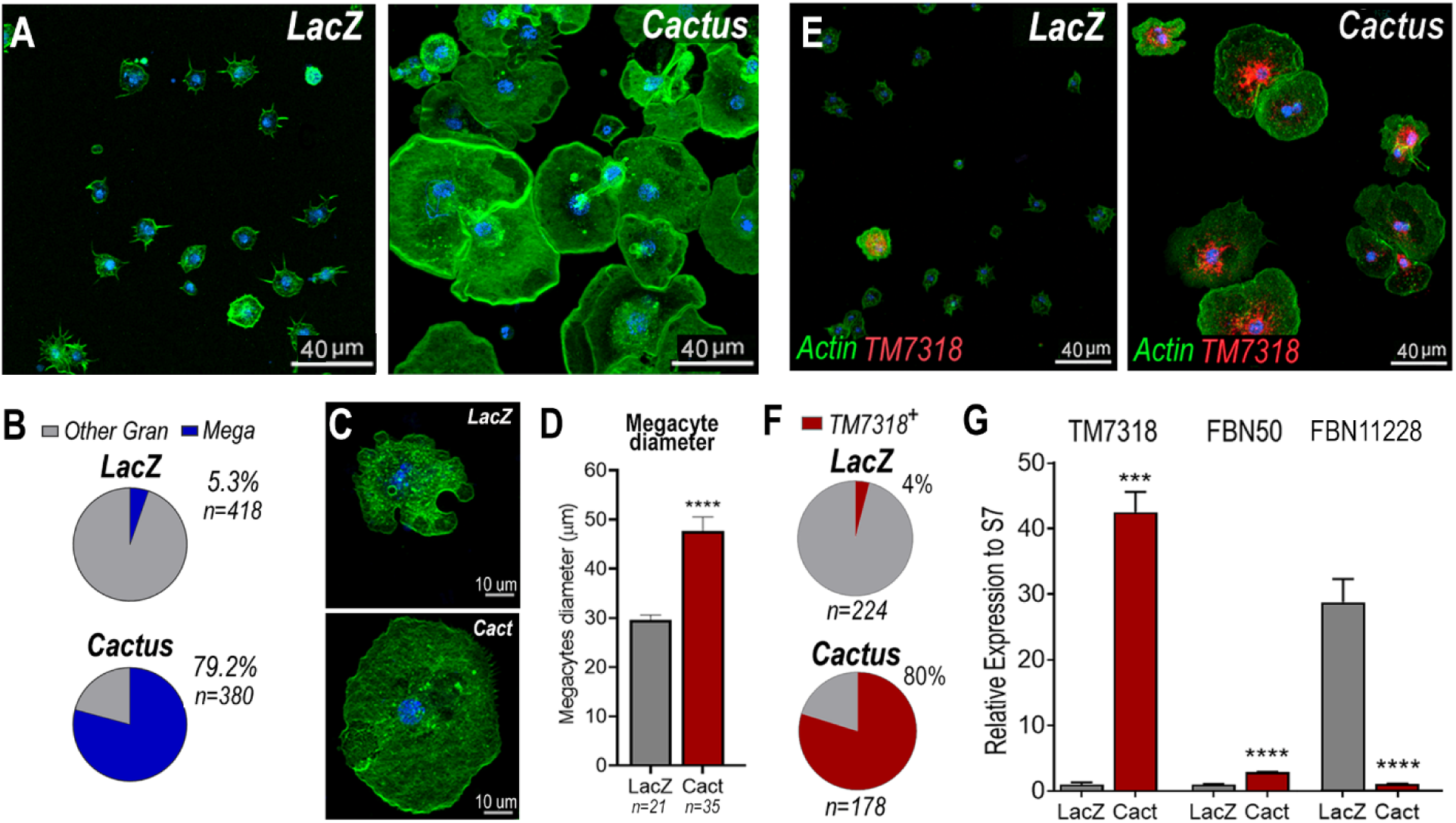
Cactus silencing promotes granulocyte differentiation into megacytes. (A) *An. gambiae* hemocytes in LacZ control and Cactus attached to a glass surface. Actin is shown in green and nuclei in blue. Scale Bar: 40µm. (B) Percentage of megacytes among all granulocytes in dsLacZ and dsCactus mosquitoes. Percentages were compared using X^2^ test. ****P≤0.0001. (C) Megacyte in control LacZ mosquitoes (upper) and in Cactus-silenced mosquitoes (lower). Actin is showing in green, and nuclei is in blue. Scale Bar: 10um. (D) Diameter of megacytes from LacZ control and Cactus-silenced mosquitoes. Error bars represent mean± SEM. Unpaired t-test. ****P≤0.0001. (E) RNA *in situ* hybridization for megacyte specific marker TM7318. Actin is shown in green (phalloidin), TM7318 mRNA in red and the nuclei in blue (Hoechst). Scale bar: 40µm. (F) Percentage of TM7318 positive cells in LacZ and Cactus silenced granulocytes. Percentages were compared using X^2^ test. ****P≤0.0001. (G) Relative mRNA expression of hemocyte specific markers in LacZ control and Cactus hemocytes for transcriptome validation. Megacyte marker (TM7318), antimicrobial granulocytes (FBN50) and regular granulocytes (FBN11228). Gene expression was normalized using RpS7 expression. Error bars represent mean ± SEM. Unpaired t-test, ****P≤0.0001.

The relative increase in megacytes in *Cactus*-silenced *An. gambiae* females could be due to enhanced megacyte proliferation or to increased differentiation of regular granulocytes into megacytes. A total hemocyte count revealed that the number of circulating hemocytes was not significantly different between *dsLacZ* (mean 17,151/mosquito, n=14) and *dsCactus* mosquitoes (mean 23,477/mosquito, n=14, Mann Whitney T-test) (Fig.S2A) 2 days post-injection. The mean number of total hemocytes was lower 4 days post-silencing, but there was also no significant difference between *dsLacZ* (mean 8,725/mosquito, n=14) and *dsCactus* mosquitoes (mean 10,836 /mosquito, n=14, Mann Whitney T-test) (Fig.S2A). Furthermore, there was no difference in the proportion granulocytes at 2 days, *dsLacZ* (3.62%) and *dsCactus* females (3.48%, Mann Whitney T-test), and 4 days, *dsLacZ* (3.91%) and *dsCactus* females (5.47%, Mann Whitney T-test), post-injection. In contrast, a significant increase in the proportion of megacyte was already apparent 2 days post-injection in *Cactus*-silenced mosquitoes (2.08% of all hemocytes, p<0.0001, Mann Whitney T-test), relative to *dsLacZ* controls (0.07%) (Fig.S2B); with a corresponding decrease in the proportion of other granulocytes from 3.5% in *dsLacZ* controls to 1.4% in *dsCactus* (Fig.S2B). At four days post injection, the differences were more pronounced, with the proportion of megacytes reaching 3.7% in *Cactus*-silenced mosquitoes (p<0.0001, Mann Whitney T-test) relative to *dsLacZ* controls (0.07%) (Fig.S2C), with a corresponding decrease in other granulocytes from 3.8% in *dsLacZ* to 1.8% in *dsCactus* mosquitoes (Fig.S2C). Taken together, these data indicate that, although the total number of hemocytes and the percentage of total granulocytes remained unchanged in response to *dsCactus* silencing, the proportion of megacytes increased at the expense of other granulocytes.

The effect of Cactus silencing on granulocyte proliferation was evaluated by quantitating the proportion of hemocytes that incorporated Bromodeoxyuridine /5-bromo-2’-deoxyuridine (BrdU), a thymidine analog. The proportion of BrdU+ hemocytes that adhered to glass (mostly granulocytes) in dsCactus mosquitoes (51%, n=694 cells) is not significantly different from dsLacZ controls (52%, n=410 cells) (Fig. S3A and B). BrdU fluorescence intensity (RFU) is also not significantly different between dsLacZ and dsCactus hemocytes (Fig. S3C). However, the ratio of BrdU fluorescence intensity to nuclear volume is significantly lower in dsCactus hemocytes (Fig. S3D). This indicates that the increase in nuclear volume in megacytes does not involve DNA replication. However, BrdU labeling can be lost over time, making it hard to establish when DNA replication occurred. The proportion of hemocytes undergoing mitosis after Cactus silencing was directly evaluated using phospho-Histone H3 (PHH3) staining, which only labels mitotically active cells. Two days post-injection, the proportion of PHH3+ hemocytes that adhered to glass (mostly granulocytes) in *dsCactus* mosquitoes (0.7%, n=896) was small, and not significantly different from *dsLacZ* controls (0.6%, n=718 cells) (Fig. S2D). At four days the proportions were also similar, with very few hemocytes positive for PHH3 staining both in *dsLacZ* (0.5%, n=799 cells) and *dsCactus* (0.3%, n=1039) (Fig. S2D). Moreover, the few hemocytes that were positive for PHH3 in *dsCactus* mosquitoes did not have the characteristic size or morphology of megacytes (Fig.S2D). These observations, together with the increase in the proportion of megacytes in *dsCactus* females, at the expense of other regular granulocytes (Fig. 1B and 2G), indicate that *Cactus silencing* promotes differentiation of granulocytes to the megacyte lineage.

### Cell dynamics of mosquito granulocytes

Megacytes are about twice as large as regular granulocytes. Regular granulocytes (Fig. 3A) reach an average diameter of 14.2 μm (Fig. 3B) when they spread over a glass surface, while the average diameter of megacytes is 28.6 μm (p<0.0001, Unpaired T-test) (Fig. 3A and 3B). Granulocyte cellular dynamics was evaluated by live imaging of perfused hemocytes *in vitro* as they adhered and spread on a glass surface. Hemocytes were labeled *in vivo*, through systemic injection of adult females with a red lipophilic dye (Vybrant CM-DiI) that accumulates on intracellular vesicles. After perfusion, a green, fluorescent probe (Cell Mask) was added to label the plasma membrane. Both regular granulocytes and megacytes attached to the glass surface and spread fully within one hour (Videos S1-S4). Megacytes already have a larger cell diameter when they first attach to glass (Fig. 3C, upper panel and Video S3), and exhibit a peripheral “halo”, corresponding to an area of extended thin cytoplasm, almost devoid of vesicles (Fig. 3C, upper panel and Video S3). Lateral views revealed that, initially, megacytes have a large nucleus and a voluminous cytoplasm in the central region of the cell that flattens dramatically as the cell “spreads” over the glass surface (Fig. 3C, lower panel and Video S4). In contrast, the central region of regular granulocytes remains mostly unchanged (Fig. 3D, lower panel and Video S2) and the periphery of the cell exhibits a modest increase in diameter as the cell spreads along the surface (Fig. 3D, upper panel and Video S1).

**Fig. 3:**
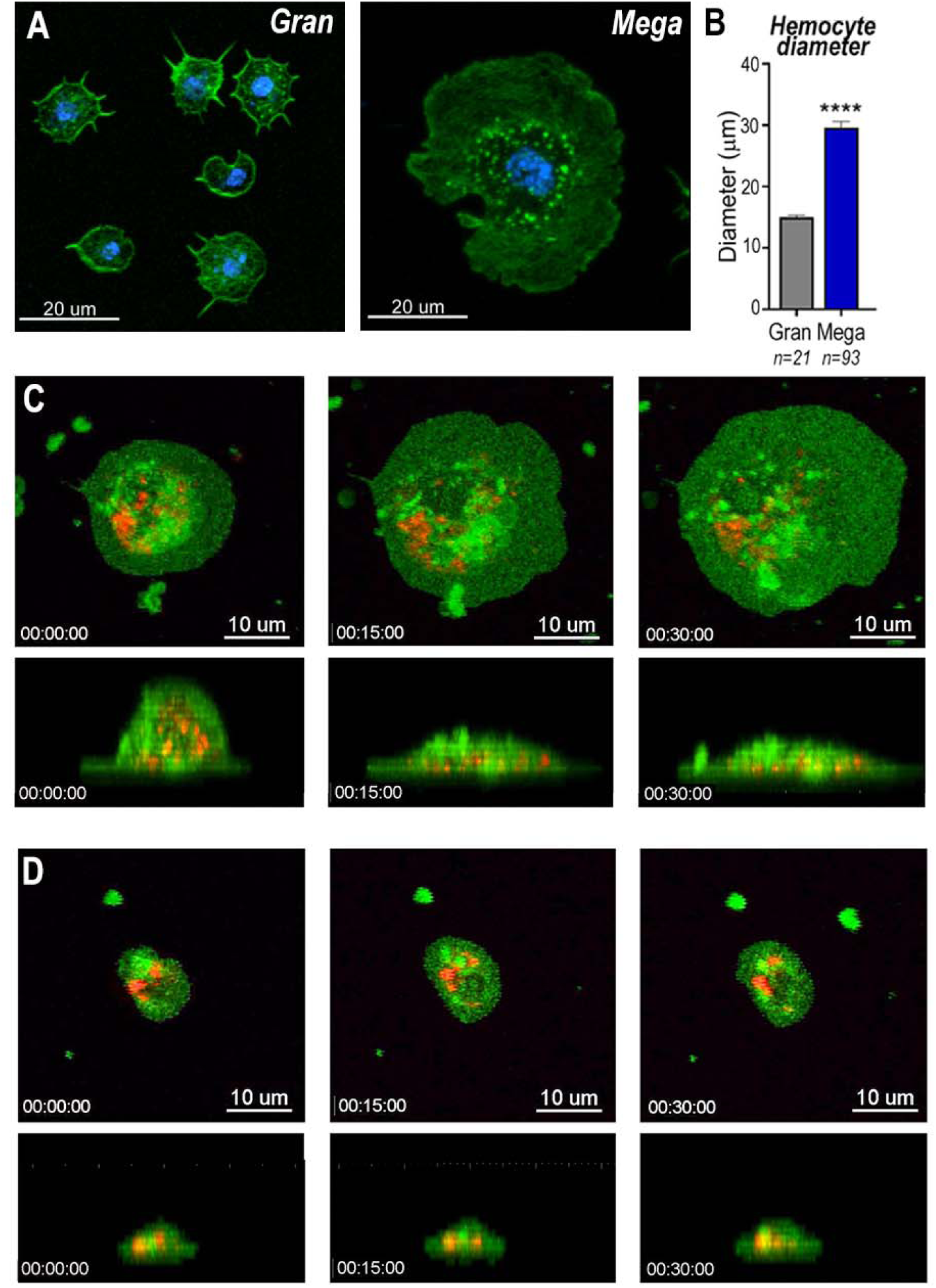
Snapshots of megacyte and granulocyte cell dynamics. (A) Regular granulocytes and megacytes from *An. gambiae* females spread on a glass surface. Actin, green (phalloidin) and nuclei, blue (Hoechst). Scale bar: 20um. (B) Granulocyte diameter of sugar-fed mosquitoes after spreading on a glass surface. Error bars represent mean± SEM. Unpaired t-test. ****P≤0.0001. (C) Live imaging time-lapse of a megacyte spreading in a glass surface for 30 minutes. Plasma membrane stained in green and microvesicles in red. Top (XY) and lateral view (XZ) of a megacyte. Scale Bars: 10µm and 5µm, respectively. (D) Live imaging time-lapse of a granulocyte spreading on a glass surface for 30 minutes. Top (XY) and lateral view (XZ) of a regular granulocyte. Scale Bars: 10µm and 5µm, respectively. (See Videos S1-S4).

### Characterization of megacyte *in vivo* dynamics and ultrastructure

The effect of *Cactus* silencing on granulocyte dynamics was evaluated *in vivo*, through live imaging of hemocytes circulating in adult female mosquitoes. Female mosquitoes were imaged for 2h, one day after blood feeding on a healthy mouse. Hemocytes were visualized by systemic injection of Vybrant CM-DiI, a fluorescent lipophilic dye that is preferentially taken up by granulocytes. Circulating hemocytes in *dsLacZ* females (presumably normal granulocytes) have a smaller diameter than those of *dsCactus* females (Videos S5 and S6) (Fig. 3A and B), and they seldom come in contact with each other as they patrol the basal surface of the midgut (Video S5). Hemocytes from *dsCactus* females (presumably megacytes) are larger and have a spindle shape (Fig. A and B, Videos S3 and S4). They appear to have higher plasticity, as they can readily stretch their cytoplasm and often come into contact with each other (Videos S6). The plasticity of dsCactus megacytes was confirmed by *in vitro* live imaging of perfused hemocytes labeled by systemic injection of Vybrant CM-DiI and green Cell Mask. Some megacytes from dsCactus mosquitoes projected long thin filopodia towards other megacytes (Video S7). This process was not observed in regular granulocytes or in megacytes from the dsLacZ controls. Taken together, our live imaging data indicates that, in addition to their larger diameter (Fig. 2C and D), *dsCactus* megacytes are also more active, have increased plasticity as they patrol the midgut (Video S6), and greater tendency to interact with each other and form clusters (Videos S6 and S7).

The detailed ultrastructure of megacytes was explored using Transmission Electron Microscopy (TEM). Hemocytes from *Cactus*-silenced females were collected by perfusion, fixed in suspension, and allowed to settle. As expected, the maximum diameter of hemocytes fixed while in suspension was smaller than when they were allowed to spread on a glass surface. However, regular granulocytes were still significantly smaller (6-10 µm) than megacytes (15-20 µm), with nuclei that are also proportionally smaller (Fig. 4A and B). Extensive electrodense areas are observed in the nuclei of megacytes, probably corresponding to the nucleolus. Large numbers of cytoplasmic vacuoles that contain abundant amorphous material are observed, as well as an extensive mitochondrial network (Fig. 4A and B). Mitochondrial organization of perfused hemocytes was further investigated using Mitotracker staining. Mitochondria of regular granulocytes have a punctate pattern with strong staining on individual organelles (Fig. 4D). In contrast, megacytes exhibit a more diffuse and extensive mitochondrial network (Fig. 4E). It is noteworthy that large membrane-bound mitochondria-like extracellular structures and small vesicles are often observed “budding off” from the surface of *dsCactus* megacytes (Fig. 4B-C), but not from regular granulocytes (Fig. 4A).

**Fig. 4:**
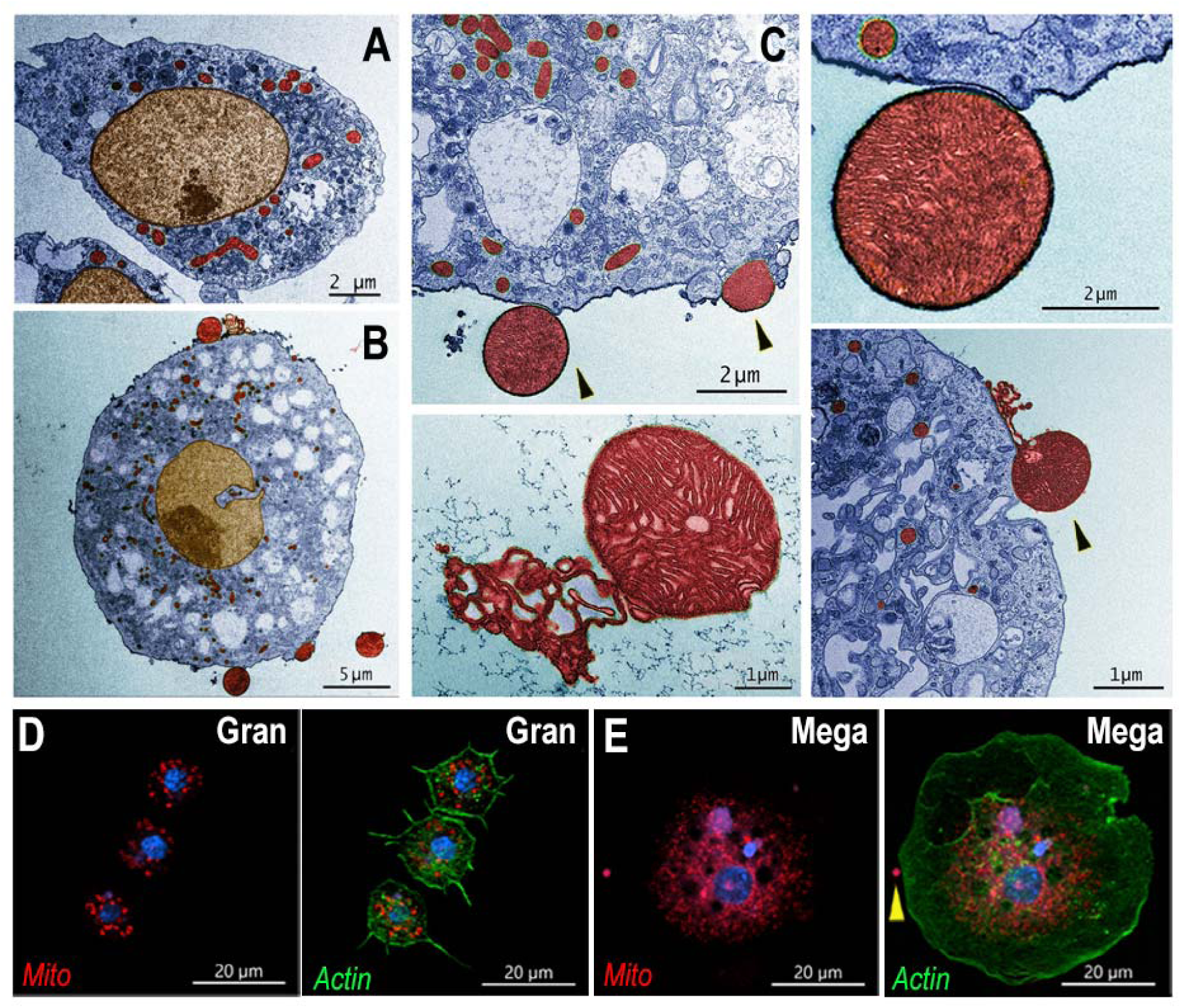
Ultrastructure of megacytes in Cactus-silenced mosquitoes. (A) Transmission Electron Microscopy (TEM) of regular granulocytes from Cactus-silenced mosquitoes. Scale Bar: 2µm. (B) TEM of megacytes from Cactus-silenced mosquitoes. Scale Bar: 5um. (C) Extracellular giant mitochondria-like structures (black arrows). Close-up of a mitochondria-like structure (lower center). Scale Bars: 2µm and 1µm. TEM images were digitally colorized, cytoplasm is shown in blue, mitochondria in red and nuclei in golden yellow. (D) Mitotracker staining in regular granulocytes. Scale Bar: 20um. (E) Mitochondrial staining of Cactus-silenced megacytes. Actin is stained in green (phalloidin), mitochondria in red (mitotracker) and nuclei in blue (Hoechst). Yellow arrow indicates an extracellular mitochondrion like structure outside of a megacyte. Scale bar: 20µm.

### Megacytes associate with the basal surface of the midgut in response to bacterial feeding

We have shown that direct contact of bacteria with epithelial cells, before the peritrophic matrix is formed, triggers PGE2 release and attracts hemocytes to the basal surface of the midgut (Barletta et al., 2019). Hemocyte recruitment to the midgut in *dsCactus* females was explored by providing a BSA protein meal containing bacteria. As expected, bacterial feeding attracted hemocytes to the midgut surface in both *dsCactus* and *dsLacZ* control females (Fig. 5A and B). However, there are important differences in hemocyte recruitment between them. In *dsLacZ* females, hemocytes attach to the midgut basal lamina individually or in doublets (Fig.5A), while hemocytes from *dsCactus* females form large clusters on the basal midgut surface, with multiple hemocytes in very close association (Fig.5B and C). *dsCactus* hemocytes on the midgut surface have the characteristic morphology of megacytes, with a larger cytoplasm and nuclei than those from *dsLacZ* hemocytes (Fig. 5B and C). Accumulation of actin was often observed in the boundaries where hemocytes from *dsCactus* females come in direct contact with each other as they form extensive clusters (Fig. 5D).

**Fig. 5:**
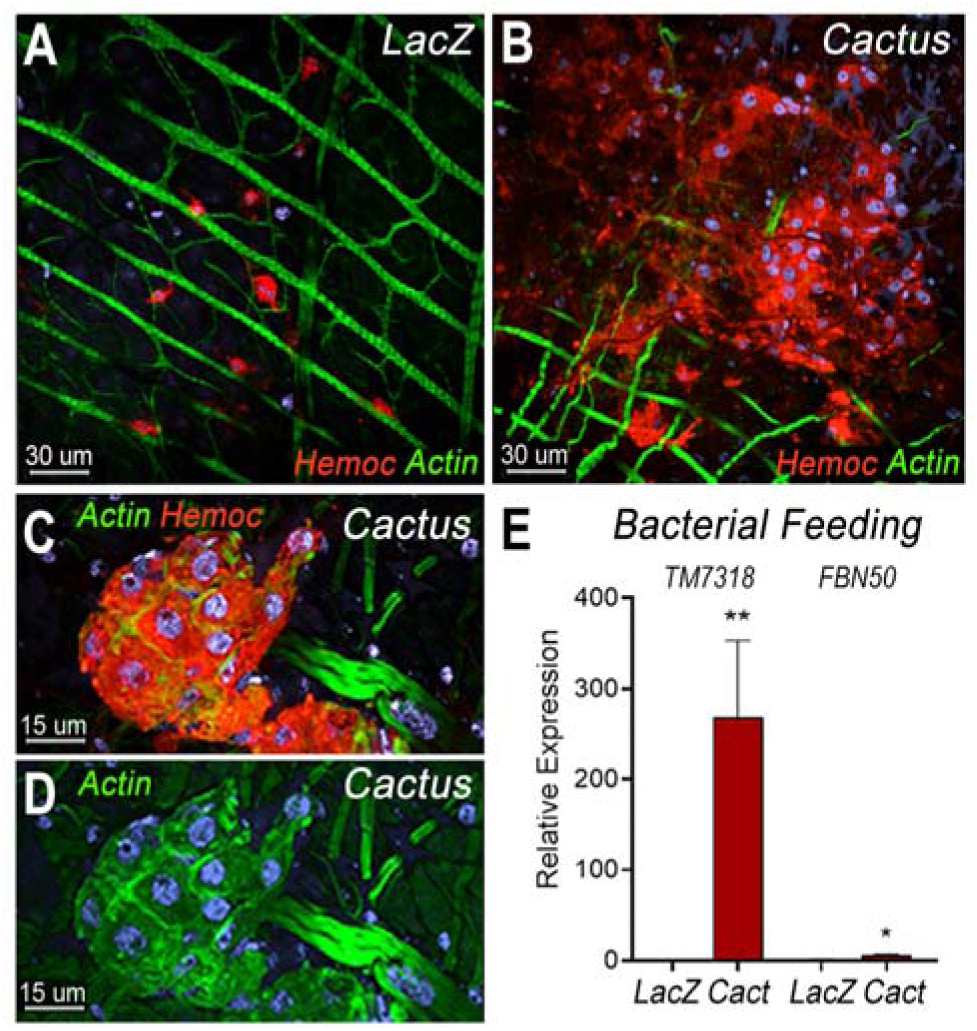
Bacterial feeding increases megacyte association to the midgut basal surface. (A) Effect of bacterial feeding in LacZ-injected controls on hemocytes associated to the midgut basal surface. (B) Effect of Cactus silencing on the hemocytes associated to the basal surface of the midgut 4 hours post bacterial feeding. (A) and (B) Scale Bar: 30um. (C) and (D) Hemocyte cluster attached to the midgut surface in Cactus-silenced mosquitoes 4 hours post bacterial feeding. Scale bar: 15 µm. (A-D) Midgut actin is shown in green (phalloidin), hemocytes (stained with Vybrant CM-DiI) in red and nuclei in blue (Hoechst). (E) Relative mRNA levels of effector hemocyte markers in the midgut 4 hours after bacterial feeding in LacZ and Cactus-silenced mosquitoes. Scale bar: 15µm. Error bars in (E) represent mean ± SEM. Unpaired t-test, *P≤0.05, **P≤0.01.

The recruitment of granulocyte subpopulations to the midgut of *dsCactus* females in response bacterial feeding was confirmed by quantitation of midgut-associated mRNAs transcripts of markers expressed in specific hemocyte subpopulations. TM7318 mRNA levels increased dramatically in *dsCactus* midguts after bacterial feeding (250-fold increase) relative to dsLacZ control (p=0.0022, Mann-Whitney test) (Fig. 5E), indicative of extensive megacyte recruitment. A significant, but more modest increase in FBN50 (5-fold) (p=0.0152, Mann-Whitney test) a marker of antimicrobial granulocytes, was also observed (Fig. 5E).

### *Toll* signaling is required for megacyte differentiation and *Plasmodium* ookinete elimination in *Cactus*-silenced females

To establish whether differentiation of granulocytes to the megacyte lineage in *dsCactus* females was mediated by the *Toll* pathway, the effect of co-silencing the *Rel1* transcription factor was evaluated. As expected, *Cactus* silencing dramatically increased the proportion of megacytes, from 3.5% to 76% (p<0.0001, X^2^ test) (Fig. 6A and B). Co-silencing *Cactus* and *Rel1* reverted this effect, resulting in a proportion of megacytes (6.6%) not significantly different from *LacZ* controls (Fig. 6B). This indicates that *Toll* signaling, through *Rel1*, mediates megacyte differentiation in *dsCactus* females.

**Fig. 6:**
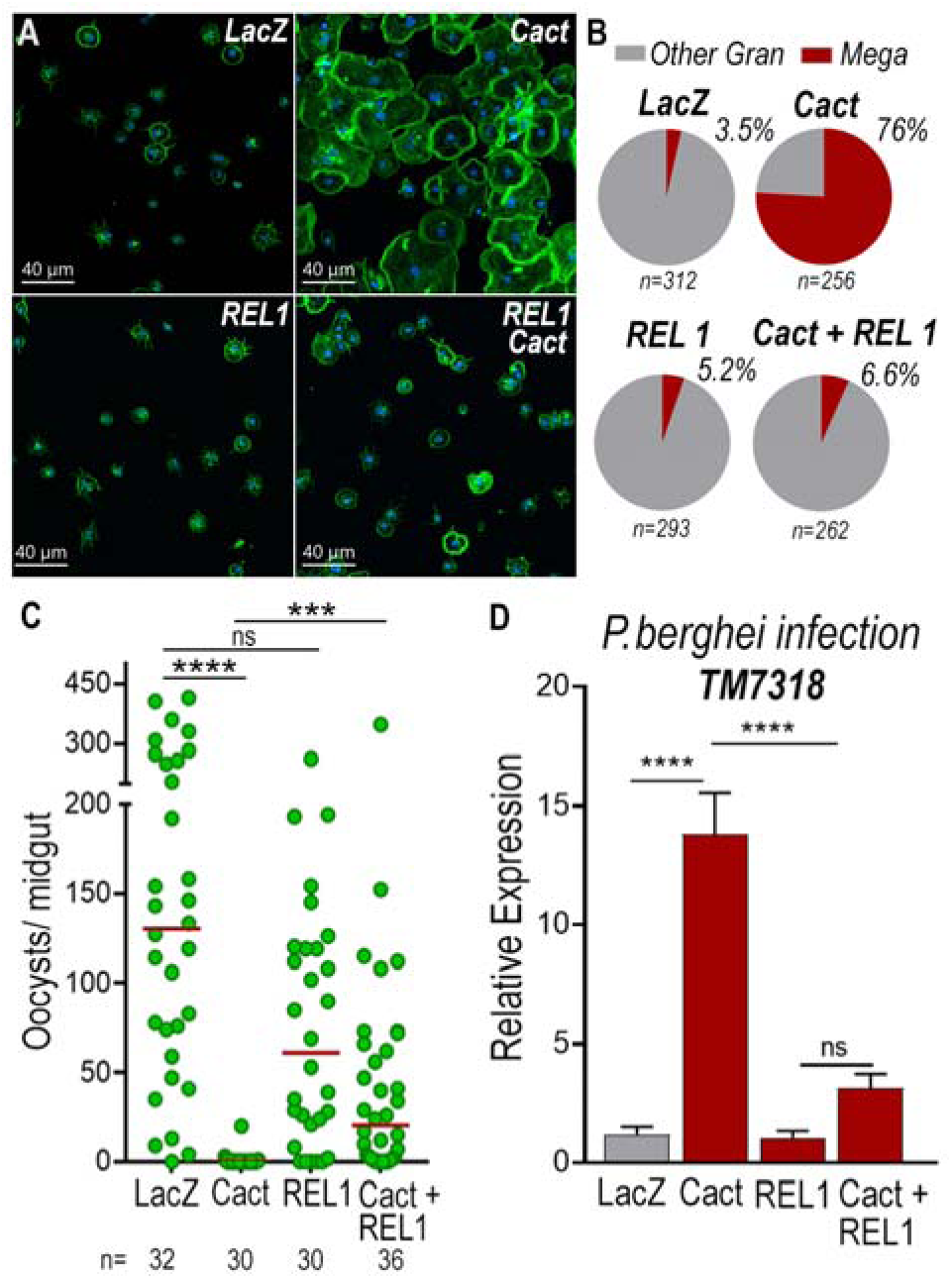
*Toll* signaling is required for megacyte differentiation and *Plasmodium* ookinete elimination in *dsCactus* females. (A) *An.gambiae* hemocytes in LacZ control, Cactus, *Rel1* and Cactus + *Rel1* attached to a glass surface. Actin is shown in green and nuclei in blue. Scale Bar: 40µm. (B) Percentage of megacytes among all granulocytes in dsLacZ, dsCactus, ds *Rel1* and dsCactus + *Rel1* mosquitoes. Percentages were compared using X^2^ test. ****P≤0.0001. (C) Mosquito susceptibility to *P. berghei* infection after dsRNA injection for LacZ, Cactus, *Rel1* and Cactus + *Rel1*. Each dot in C represent the number of oocysts or hemocytes, respectively, for individual midguts. The median is indicated by the red line. Mann-Whitney U test, ****p ≤ 0.0001; ***p ≤ 0.001, NS, p > 0.05. (D) Relative mRNA levels of TM7318, megacyte marker, in the midgut 26 h post *P. berghei* infection (post-invasion) in LacZ, Cactus, *Rel1* and Cactus + *Rel1* silenced mosquitoes. Error bars in (D) represent mean ± SEM. Unpaired t-test, ****P≤0.0001, NS, p > 0.05.

We next investigated the effect of *Toll* signaling on the immune response to *Plasmodium* of *dsCactus*-silenced females and megacyte HdMv release. *Cactus* silencing drastically reduced oocyst numbers (median= 0, p< 0.0001, ANOVA, Dunn’s multiple comparison test) relative to LacZ controls (median= 127) (Fig. 6C), and co-silencing *Cactus* and *Rel1* significantly increase oocyst numbers (median= 24, p=0.0002, ANOVA, Dunn’s multiple comparison test) (Fig. 6C), in agreement with previous reports (Frolet *et al.,* 2006). A strong increase in TM7318 mRNA associated with the midgut 24 h post-infection was detected in *dsCactus* infected females (13-fold increase), relative to dsLacZ (p<0.0001, Mann-Whitney test) (Fig. 6D and S4), indicative of midgut recruitment of megacytes. Furthermore, when *Rel1* was co-silenced with *Cactus*, the levels of midgut-associated TM7318 mRNA decreased and were not significantly different from those of *dsLacZ* controls. In contrast, mRNA levels of FBN11228, a marker of regular granulocytes, did not change significantly in *dsCactus* females or after co-silencing *Rel1* and *Cactus*, relative to *dsLacZ* (Fig. S5).

## Discussion

We recently described specific subsets of mosquito granulocytes based on single-cell transcriptomic analysis (Raddi et al., 2020). Here we present a functional characterization of megacytes, a newly described subpopulation of final effector granulocytes and provide direct evidence of their recruitment to the basal surface of the mosquito midgut and their participating in the mosquito immune response to ookinete midgut invasion. The almost complete elimination of *P. berghei* parasites by the mosquito complement system when Cactus is silenced was documented more than fifteen years ago (Frolet et al., 2006). However, the mechanism by which Cactus silencing enhanced hemocyte responses to *Plasmodium* infection remained a mystery.

Our transcriptomic analysis indicated that Cactus silencing increased the proportion of circulating megacytes, at the expense of regular granulocytes (Fig. 2). This was confirmed by morphological analysis, *in situ* hybridization, cell counts and mRNA quantitation of hemocyte-specific markers, TM7318 (megacytes) and FBN11228 (regular granulocytes). We also provide direct evidence that, besides being larger, megacytes also have higher plasticity, as they can greatly extend their cytoplasm and flatten their nucleus as they spread on a glass surface (Fig. 1C).

The lack of DNA replication and the concomitant reduction in the proportion of regular granulocytes, indicates that circulating megacytes increase in response to Cactus silencing by promoting final differentiation of granulocytes to the megacyte lineage. Besides the dramatic increase in circulating megacytes, Cactus silencing also results in megacytes that are even larger and more plastic than megacytes from dsLacZ controls. Fine ultrastructural analysis revealed that the cytoplasm of megacytes exhibits extensive large vacuolar structures filled with amorphous material, as well as small vesicles and mitochondria-like structures that are secreted from the cell membrane. In vertebrates, mitochondrial extrusion has been recently documented as a trigger of inflammation. Activated platelets release their mitochondria, both within microparticles or as free organelles; and secreted phospholipase A2 IIA can hydrolyze the membrane, releasing inflammatory mediators, such as lysophospholipids, fatty acids, and mitochondrial DNA, that promote leukocyte activation. Furthermore, extracellular mitochondria also interact directly with neutrophils *in vivo*, and trigger their adhesion to the endothelial wall (Boudreau et al., 2014). Activated monocytes release mitochondria, and their proinflammatory effect on endothelial cells is determined by the activation status of the monocytes that released them. It has been proposed that free mitochondria could be important mediators of cardiovascular disease by inducing activation of type I IFN and TNF signaling (Puhm et al., 2019).

Large numbers of megacytes were recruited to the midgut of Cactus-silenced females in response to bacterial feeding, forming extensive clusters of cells in close contact with each other, indicating that Cactus silencing also results in functional differences in megacytes. Expression of midgut associated markers of specific hemocyte subpopulations indicates that *Plasmodium* midgut invasion triggers strong recruitment of megacytes to the basal surface of the midgut in *dsCactus* females, in agreement with the documented increase in HdMv associated with epithelial cells invaded by ookinetes (Castillo et al., 2017). We also show that co-silencing the transcription factor *Rel1* and *Cactus* disrupts the differentiation of granulocyte to the megacyte lineage observed when only *Cactus* is silenced, indicating that megacyte differentiation requires a functional *Toll* pathway. Co-silencing *Rel1* and *Cactus* also reduced midgut recruitment of megacytes 24 h post-infection, a critical time when ookinetes are invading the mosquito midgut, and significantly increases *Plasmodium* survival, relative to *dsCactus* females. We propose that Toll signaling promotes hemocyte differentiation into the megacyte lineage, and that the dramatic increase in the proportion of circulating megacytes and their midgut recruitment mediates the documented increase in HdMv (Castillo et al., 2017) that promotes activation of the mosquito complement system that ultimately eliminates *P. berghei* ookinetes. The release of free mitochondria-like structures by megacytes from Cactus-silenced females raises the question of whether this is an ancient systemic danger signal that promotes immune activation.

## Data Availability

The raw data and detailed information on individual experiments and number of replicates are available at Supplementary tables file.

## Acknowledgments

This work was supported by the Intramural Research Program of the Division of Intramural Research Z01AI000947, NIAID, National Institutes of Health. We thank Kevin Lee, Yonas Gebremicale and André Laughinghouse for insectary support, and Asher Kantor for editorial assistance.

## Author Contributions

Experiments were designed by A.B.F.B., N.T., B.S., G.R. and C.B.M., carried out by A.B.F.B. B.S., N.T., and analyzed by A.B.F.B., N.T., B.S., G.R. and C.B.M. A.B.F.B. and C.B.M. wrote the paper.

## Material and Methods

### Mosquitoes and mouse feeding

*Anopheles gambiae* mosquitoes (G3 strain – CDC) were reared at 28°C, 80% humidity under a 12h light/ dark cycle and kept with 10% Karo syrup solution during adult stages. For mosquito infections with *Plasmodium berghei,* we used the transgenic GFP *P.berghei* parasites (ANKA 2.34 strain) kept by serial passages into 3-4 weeks old female BALB/c mice (Charles River, Wilmington, MA) starting from frozen stocks. Mouse infectivity was evaluated before feeding by parasitemia levels from Giemsa-stained thin blood films and *in vitro* microgamete exflagellation counting. Briefly, one microliter of tail blood was mixed with 9ul of gametocyte activating medium (RPMI 1640 with 25mM HEPES + 2mM glutamine, Sodium Bicarbonate 2g/L, 100uM xanthurenic acid, 50ug/ml hypoxanthine). After 10 minutes of incubation exflagellations were quantified using a 40X objective by phase contrast. Four to five-day old mosquitoes were fed when mice reached 3-5% parasitemia and 2-3 exflagellation per field. To feed blood-fed control mosquitoes, three-to four-week-old uninfected mice were used. Following feeding, both control and infected mosquitoes were maintained at 19°C, 80% humidity and 12h light/dark cycle until the day of dissection.

### Ethics statement

Public Health Service Animal Welfare Assurance #A4149-01 guidelines were followed according to the National Institutes of Health Animal (NIH) Office of Animal Care and Use (OACU). These studies were done according to the NIH animal study protocol (ASP) approved by the NIH Animal Care and User Committee (ACUC), with approval ID ASP-LMVR5.

### Perfused hemocytes live imaging

Three-day-old adult females were injected with Vybrant DiI (1:10 water diluted, ThermoFisher Scientific, Waltham, MA, USA) on one side of the thorax. The next day, mosquitoes were injected with 69 nL of either dsCactus or dsLacZ at 3 µg/µL on the other side of the thorax. After 4 days, hemocytes were ready for perfusion or mosquitoes were used for in vivo live imaging as described below. Mosquitoes were cold-anesthetized and, using forceps, a small cut was made in the abdomen. Transfer buffer (95% Schneider media + 5% citrate buffer) was injected at the thorax and 10-15 µL of hemolymph was harvested at the cut-site. This was repeated for 5-7 mosquitoes and collected in a microcentrifuge tube stored on ice. To stain the plasma membrane of hemocytes we used CellMask green plasma membrane stain stock solution (C37608, Invitrogen, Waltham, MA, USA) and for the nuclei we used the Hoechst 33342 Solution (20mM) (ThermoFisher Scientific, Waltham, MA, USA). Two microliters of fluorescent label solution (58 µL H2O + 1 µL Cell Mask stock + 1µL Hoechst stock) was added for every 20 µL of perfusion and 100 µL of this mixture was mounted on an ibidi µ-Slide 18 Well Glass Bottom slide. Cells were allowed to settle for 30 minutes then imaged. Images were taken on a Leica SP5 confocal microscope using a 63x 1.4 NA oil objective with 405 nm wavelength laser (at 3% transmission) for Hoechst, 488 nm (5%) for Cell Mask, and 561 nm (3%) for DiI. Pinhole was set to 1 AU and frame average was 12. Z-intervals of 1-2 µm encompassing the full cell height was taken every 5 minutes for 2 hours.

### Bacterial artificial feeding

We used a bacterial mixture obtained from the midguts of the Anopheles gambiae G3 from our colony (Barletta et al., 2019). A pre-inoculum was set up in LB media from the frozen stocks containing the bacterial mixture and allow to grow overnight at 28°C, 250rpm in a shaker incubator. At the day of the experiment, the pre-inoculum was diluted in fresh LB media and allowed to grow for 2 hours in the same condition described above. Briefly, after 2 hours of growth, bacteria were washed with sterile PBS to remove toxins and the concentration of the culture was estimated based on the Optical Density (OD) of the culture. At 600nm, 1OD was considered the equivalent of 10^9^ bacteria/mL. Three-to-four day mosquitos were fed a sterile 10% sucrose solution containing antibiotics (Penicillin, 100U/mL and Streptomycin, 100ug/mL) for 2 days prior the bacterial feeding. Control group was fed with a sterile 10% Bovine Serum Albumin (BSA) solution in HBSS without calcium and magnesium and the bacteria group was fed with the same solution containing 4 x 10^9^ bacteria per feeder. Mosquitoes were dissected 6 hours post feeding for visualization of hemocytes attached to the midgut basal surface.

### Hemocyte collection, morphology staining and quantification

Hemocytes were collected by perfusion using anticoagulant buffer (60% Schneider medium, 30% citrate buffer, pH 4.5 and 10% FBS), pH was adjusted to 7-7.2 after mixing all the components. After perfusion, hemocytes were placed in a µ-slide angiogenesis chamber (ibidi GmbH, Gräfelfing, Germany) and were allowed to settle for 15 minutes. Cells were fixed for an hour at room temperature by adding 16% paraformaldehyde (PFA) solution in anticoagulant buffer to a final concentration of 4%. Following fixation cells were washed with PBS 0.1% Triton and incubated for 30 minutes at room temperature with 1U of phalloidin (Alexa Fluor 488, Molecular Probes, ThermoFisher Scientific, Waltham, MA, USA) and 20 µM Hoechst 33342 (405, Molecular Probes, ThermoFisher Scientific, Waltham, MA, USA), both diluted in PBS 0.1% Triton. Cells were then placed in mounting media for storage by adding 2 drops of Prolong Gold Antifade Mountant (Molecular Probes, ThermoFisher Scientific, Waltham, MA, USA). For determination of proportion of megacytes upon Cactus silencing, the hemocytes were imaged, the diameter of every cell was measured and classified as granulocytes (cell diameter >12.5-25 µm) or megacytes (cell diameter >25 µm) as mentioned before. The total number of granulocytes and megacytes obtained from hemolymph pooled from 16-20 mosquitoes was noted and the percentage of megacytes amongst granulocytes was determined for each sample. Data from three independent biological replicates were used to plot the graphs.

### Measurement and categorization of the hemocytes by size

The mosquito hemolymph was collected and the hemocytes were allowed to attach on a coated well of 15µm chamber slide. For each well 8-10 mosquitoes were bled and for every sample, bleeding was done in two wells with a total of 16-20 mosquitoes. Post attachment, the hemocytes were fixed with 4% p-formaldehyde and stained with Phalloidin and DAPI to visualize the morphology. Images were taken for at least 10 random fields for each well and the images were used to measure the cell diameter using Imaris software. Using the “Pairs” option of “Measurement points” tool in the software, the largest diameter of every cell was determined. For categorizing the hemocytes into different subtypes, the following size reference was followed for every image analysis. Cells with diameter ranging from 4-7.5µm were classified as prohemocytes, >7.5 µm-12.5 µm as oenocytoids, >12.5-25 µm as granulocytes and >25 µm as megacytes.

### dsRNA synthesis

Three-to-four day old female *An.gambiae* females were cold-anesthetized and injected with 69nl of a 3ug/ul dsCactus or dsLacZ control. Double-stranded RNA for *Cactus* (AGAP007938) was synthesized by *in vitro* transcription using the MEGAscript RNAi kit (Ambion, ThermoFisher Scientific, Waltham, MA, USA). DNA templates were obtained by PCR using *An.gambiae* cDNA extracted from whole body sugar-fed females. A 280-bp fragment was amplified with primers containing T7 promoters (F-**TAATACGACTCACTATAGGG**TAACACTGCGCTTCATTTGG and R-**TAATACGACTCACTATAGGG**GCCCTTTTCAATGCTGATGT), using an annealing temperature of 58°C. Double-stranded RNA for LacZ was synthetized by amplifying a 218-bp fragment from LacZ gene clones into pCRII-TOPO vector using M13 primers to generate a dsRNA control as previously described (Molina-Cruz et al., 2012). A 386-bp fragment from *Rel1* gene was amplified using primers containing T7 promoters (F-**TAATACGACTCACTATAGGGA**TCAACAGCACGACGATGAG and R-**TAATACGACTCACTATAGGG**TCGAAAAAGCGCACCTTAAT) using an annealing temperature of 58°C. For double silencing experiments, 138 nl of dsRNA mixture at 3ug/ul was injected into female *Anopheles gambiae*.

### RNA extraction and bulk RNAseq library preparation

Hemocytes were collected as previously described above. In short, *An.gambiae* females were perfused using anticoagulant buffer and immediately transferred to a glass tube for attachment. After one hour, hemocytes that did not attach to the glass tube were collected and transferred to a 1.5 ml microcentrifuge containing 800ul of TRIZOL LS reagent (Invitrogen, Waltham, MA, USA), that correspond to the unbound fraction enriched mainly by prohemocytes and oenocytoids. Hemocytes that attached to the glass surface were washed twice with PBS and resuspended in 1mL of TRIZOL LS reagent (Invitrogen, Waltham, MA, USA), this corresponds to the bound fraction, mainly enriched by granulocytes. Hemocytes were then lysed in TRIZOL reagent for 15-30 minutes at room temperature to allow for full dissociation, then stored at 4°C overnight and then at −20C until RNA extraction. The homogenate of hemocyte samples were transferred to Phase Lock Gel Heavy 2 mL tubes (QuantaBio, Beverly, MA, USA) that had been pre-spun for 1500 RCF for 1 minute, and allowed to incubate for 5 minutes at room temperature. 100 uL of chloroform (200 uL per 1 mL TRIZOL or TRIZOL plus media) was added, the tubes capped, and then vigorously shaken for 15 seconds. Samples were then centrifuged for 12,000 RCF, 10 minutes, 4°C. If the clear, aqueous phase was still mixed with TRIZOL matrix then 100 uL more of chloroform was added, and the samples again mixed vigorously and spun as before. The aqueous phase was then transferred to a fresh 1.5 mL Eppendorf tube and the RNA precipitated by adding 0.25 mL of isopropyl alcohol (500 mL per 1 mL TRIZOL reagent used). 20 uL of glycogen (5 mg / mL) were also added to aid in precipitation and pelleting. Samples were mixed by repeated inversion 10 times, incubated for 10 minutes at room temperature, and then spun at 12,000 RCF, 10 minutes, 4°C. All the supernatant was removed, and the RNA pellets washed twice with 75% ethanol (minimum 1 mL of ethanol per 1 mL of TRIZOL used). Each time the samples were mixed by vortexing and centrifuged 7,500 RCF, 5 minutes, 4C. At the end, the supernatant was removed and samples air-dried until almost dry, but not completely (still translucent). RNA was resuspended with 30 uL of RNAse free water, pipetting a few times to homogenize and then incubating at 55°C for 10 minutes to completely resuspend. Samples were then stored at −20C until library preparation by Bespoke Low-Throughput Team at the Wellcome Sanger institute. Total RNA quantity was assessed on a Bioanalyser and ranged from 300 ng to 39 ng. mRNA was then isolated with the NEBNext Poly(A) mRNA magnetic isolation module. RNA-seq libraries were prepared from mRNA using the NEBNext Ultra II Directional RNA Library Prep Kit for Illumina (New England Biolabs) as by manufacturer instructions, except that a proprietary Sanger UDI (Unique Dual Indexes) adapters / primer system was used. Furthermore, Kapa Hifi polymerase rather than NEB Q5 was employed. For bulk RNAseq sequencing samples libraries were run on the Illumina HiSeq 4000 instrument with standard protocols using a 150-cycle kit set to a 75bp paired-end configuration. Libraries supplied at 2.8 nM and loaded with a loading concentration of 280 pM.

### Bulk RNA-seq bioinformatic analysis

Sequencing reads in CRAM format were fed into a personal BASH pipeline to convert cram files to fastq using biobam’s bamtofastq program (Version 0.0.191) (Raddi et al., 2020). Forward and reverse fastq reads in paired mode were aligned to the A. gambiae AgamP4.3 reference genome using hisat2 (Version 2.0.4) and featureCounts (Version 1.5.1) with recommended settings. Count matrices were combined before downstream data processing and analysis within R version 3.5.3 (RStudio version 1.0.153). Downstream normalization, differential expression analysis and visualization were done with DESeq2 R package (Version 1.18.1) (Love et al., 2014). Base factor was defined as the LacZ, unbound condition. Data was normalized by making a scaling factor for each sample. First the log(e) of all the expression values were taken, then all rows (genes) were averaged (geometric average). Genes with zero counts in one or more samples were filtered out and the average log value from log (counts) for all genes was subtracted. Finally, the median of the ratios calculated as above for each sample was computed and raised to the e to make the scaling factor. Original read counts were divided by the scaling factor for each sample to get normalized counts. Then, the dispersion for each gene was estimated, and a negative binomial generalized linear model fitted. P values for the differential expression analysis were adjusted for multiple testing using the Bonferroni correction. Genes were considered as differentially expressed in Cactus knockdown compared to LacZ control if they had an adjusted P value < 0.001 (Wald T-test) and a log2 fold change > 2. Gene lists with vectorbase IDs were converted to gene annotations with g:Profiler (Raudvere et al., 2019). g:Profiler utilizes Ensembl as its primary data source and is anchored to its quarterly release cycle. g:GOSt was used to perform functional enrichment analysis on input gene lists to map the data onto enriched biological processes or pathways. In addition to Ensembl, also KEGG, Reactome, WikiPathways, miRTarBase, and TRANSFAC databases were used. Functional enrichment is evaluated with a cumulative hypergeometric test with g:SCS (Set Counts and Sizes) multiple testing correction (adjusted P value reported only < 0.05). Gene lists were ordered on log-fold changes. Complete dataset is available publicly in https://www.ebi.ac.uk/biostudies/arrayexpress/studies/E-MTAB-11252.

### Transmission Electron Microscopy (TEM)

Hemocytes were collected by perfusion using anticoagulant buffer, described above and they were allowed to settle on Thermanox™ coverslips (Ted Pella, Redding, CA) for 15 minutes at room temperature then fixed 2.5% glutaraldehyde in 0.1 M sodium cacodylate buffer overnight at 4°C, and then post-fixed 1hr with 1.0% osmium tetroxide/0.8% potassium ferricyanide in 0.1 M sodium cacodylate buffer, washed with buffer then stained with 1% tannic acid in dH2O for one hour. After additional buffer washes, the samples were further osmicated with 2% osmium tetroxide in 0.1M sodium cacodylate for one hour. The samples were then washed with dH2O and additionally stained overnight with 1% uranyl acetate at 4°C, dehydrated with a graded ethanol series, and embedded in Spurr’s resin. Thin sections were cut with a Leica UC7 ultramicrotome (Buffalo Grove,IL) prior to viewing at 120 kV on a FEI BT Tecnai transmission electron microscope (Thermo fisher/FEI, Hillsboro, OR). Digital images were acquired with a Gatan Rio camera (Gatan, Pleasanton, CA).

### Mitotracker staining

Hemocytes were perfused with anticoagulant buffer, described above. Cells were incubated at room temperature for 15 minutes for spreading. Then washed three times with 95% Schneider media, 5% citrate buffer to remove most of the serum from the cells. Hemocytes were placed with 200nM Deep Red Mitotracker 644/665 which is retained after fixation (Molecular Probes, ThermoFisher Scientific, Waltham, MA, USA) diluted in 95% Schneider media, 5% citrate buffer. Cells were incubated for 45 minutes at room temperature in the dark, then washed with PBS and fixed with 4% Paraformaldehyde in PBS for 15 minutes at room temperature. Hemocytes were then counterstained with phalloidin and Hoechst as described above.

### TM7318 in situ hybridization (ISH)

The ISH protocol includes a permeabilization step with a protease treatment, which compromises the cell morphology. To evaluate the morphology of hemocytes and RNA expression by ISH, we used a two-step protocol to image morphology first and then proceed to image the probes, described in (Raddi et al., 2020). Hemocytes collected by perfusion four days after dsCactus injection, fixed and stained with Alexa 488 phalloidin (actin) as described above. Ten random fields of each well were imaged using a tile scan “mark and find” tool, where coordinates of the field are recorded and can be restored to image the same cells later. Then, hemocytes were subjected to ISH using RNAscope multiplex fluorescent reagent kit v2 assay (cat# 323110, ACDBio, Abingdon, United Kingdom) following the manufacturer’s instructions. TSA based fluorophores Opal 4-color automation IHC kit (cat # NEL801001KT, PerkinElmer, Waltham, MA, USA) was used for the development of fluorescence (Opal 620 – C3). A specific RNA probe for TM7318 (cat# 543201-C3; Aga-Transmembrane-C3) designed by ACDBio was used to stain specifically megacytes. At the end of the ISH protocol, hemocytes were placed in prolong gold and re-imaged using the “mark and find” tool to recall the positions of the morphology pictures. Images were merged using Imaris 9.3.1 (Bitplane, Concord, MA, USA). Each well was imaged taking 12 fields per well. Post imaging, the cell diameter of every cell was measured by Phalloidin stain as described previously and the total number of granulocytes were determined for each sample. Amongst the granulocytes and larger cells (cells with diameter >12.5), the number of cells positive for the TM7318 probe were counted and their percentage was determined for both the control and Cactus silencing.

### Confocal microscopy and Tile scan imaging

Confocal images were captured using a Leica TCS SP8 (DM8000) confocal microscope (Leica Microsystems, Wetzlar, Germany) with either a 40x or a 63x oil immersion objective equipped with a photomultiplier tube/ hybrid detector. Hemocytes were visualized with a white light laser, using 498-nm excitation for Alexa 488 (phalloidin); 588-nm excitation for Opal620 (TM7318 probe) and Vybrant DiI (hemocytes); 644-nm excitation for Deep Red Mitotracker (Mitochondria) and a 405-nm diode laser for nuclei staining (Hoechst 33342). Images were taken using sequential mode and variable z-steps. For combined morphology and in RNA in situ hybridization, we used tile scan “mark and find” tool included in LASX software to capture the same areas of the slide before and after the hybridization. Image processing and merge was performed using Imaris 9.3.1 (Bitplane, Concord, MA, USA) and Adobe Photoshop CC (Adobe Systems, San Jose, CA, USA).

### RNA extraction, cDNA synthesis and qPCR analysis

*An. gambiae* hemocytes were collected as described above four days after dsRNA injection (dsLacZ and dsCactus). Hemolymph pools of 20 mosquitoes (5ul/ each mosquito) were placed directly into 800ul of TRIzol LS reagent (ThermoFisher Scientific, Waltham, MA, USA). For midgut RNA extraction, pools of 20 midguts were homogenized directly in 1mL TRIzol reagent. RNA extraction was carried out as described above in the section *RNA extraction and bulk RNAseq library preparation.* Total extracted RNA was resuspended in nuclease free water and one microgram was used for cDNA synthesis using the Quantitect reverse transcription kit (Qiagen, Germantown, MD, USA) following the manufacturer’s instructions. Quantitative PCR (qPCR) was used to measure FBN11228 (AGAP011228), TM7318 (AGAP007318) and FBN50 (AGAP005848) gene expression in hemocytes cDNA. We used the DyNamo SYBR green qPCR kit (ThermoFisher Scientific, Waltham, MA, USA) with target specific primers and the assay ran on a CFX96 Real-Time PCR Detection System (Bio-Rad, Hercules, CA, USA). A 139-bp fragment was amplified for FBN11228 (F-CCAGCATCGGTACAACGGAA and R-AAGCTCGTGTTTTCGTGCTG). A 150-bp fragment was amplified for TM7318 (F-AAAACATCCAGAAACACGCC and R-GGATTCCGGTTAAGTCCACC). A 92-bp fragment was amplified for FBN50 (F-ATCACAAGGTTCCGGCTATG and R-CGTTGGTGTAGGTGAGCAGA). Relative expression was normalized against An. gambiae ribosomal protein S7 (RpS7) as internal standard and analyzed using the ΔΔ Ct method (ref – Livak and Schmittgen, 2001; Pfaffl, 2001). RpS7 (AGAP010592) primers sequences were: F-AGAACCAGCAGACCACCATC and R – GCTGCAAACTTCGGCTATTC. Statistical analysis of the fold change was performed using Unpaired t-test (GraphPad, San Diego, CA, USA). Each independent experiment was performed with three biological replicates (three pools of 20 mosquitoes) for each condition.

### In vivo live imaging

Mosquitoes were prepared the same way for imaging of perfused hemocytes and injected with Vybrant DiI cell labelling (ThermoFisher Scientific, Waltham, MA, USA) for both dsCactus and dsLacZ. After 4 days, mosquitoes were starved in the morning and then fed on a BALB/c mouse in the afternoon. Imaging took place the next day at 18-20 hours post-bloodmeal. Mosquitoes were imaged as previously described (Trisnadi and Barillas-Mury, 2020). Briefly, 5-10 mosquitoes with legs and head removed were placed between a coverslip and glass slide with craft putty as a spacer. Images were taken on a Leica SP5 confocal microscope using a 40x 1.25 NA oil objective with 561 nm (3%) for Vybrant DiI. A z-stack with 1 µm intervals was taken to include hemocytes circulating in the hemolymph to the midgut lumen. The z-stack was taken every 1 minute for 1-2 hours.

### Visualizing hemocytes attached to the midgut basal lamina

To preserve hemocyte-midgut bound, midguts were quick fixed using a higher concentration of fixative injected straight into the hemolymph of the mosquito (207nl of 16% paraformaldehyde). To stain hemocytes, the day before the dsRNA treatment (dsLacZ and dsCactus), three-to-4-day-old mosquitoes were injected with 69nl of a 100uM solution Vybrant CM-DiI cell labelling solution (ThermoFisher Scientific, Waltham, MA, USA), final concentration in the hemolymph (approximately 3.5uM). Engorged mosquitoes fed with 10% BSA solution containing bacteria were anesthetized and injected with 207nl of 16% paraformaldehyde, rested 40 seconds before midgut dissection in 4% paraformaldehyde solution. After dissected, midguts were placed in ice-cold PBS and opened longitudinally, and the bolus was removed. Clean opened tissues were then fixed overnight at 4°C in 4% paraformaldehyde. The following day, midguts were washed twice with PBS, blocked for 40 minutes with PBS containing 1% BSA and washed twice with the same solution. For actin and nuclei staining, midguts were incubated for 30 minutes at room temperature with 1U of phalloidin (Alexa Fluor 488, Molecular Probes, Waltham, MA, USA) and 20uM Hoechst 33342 (405, Molecular Probes, Waltham, MA, USA), both diluted in PBS. Tissues were mounted in microscope slides using Prolong Gold Antifade mounting media (Molecular Probes, Waltham, MA, USA). Hemocytes were visualized by confocal microscopy and the number of hemocytes per midgut in each biological condition was also analyzed.

### PHH3^+^ and BrdU staining

For BrdU staining, *An. gambiae* females injected with dsRNA were treated for 3 days with a sugar solution containing 1mg/ml Bromodeoxyuridine (Sigma Aldrich, St. Louis, MO, USA). At day four post dsRNA injection, hemocytes were collected with anticoagulant buffer (70% Schneider media, 30% citrate buffer and 10% FBS) pH 7.4. Hemocytes were allowed to settle on a ibidi µ-Slide 18 Well Glass Bottom slide (Gräfelfing, Germany) for 15 minutes at room temperature and then fixed for 30 minutes with 4% PFA followed by a permeabilization step with PBS 0.5% Triton for 20 minutes. Hemocytes were washed twice with PBS with 1% BSA before treatment with 2N HCL for 40 minutes to denature the DNA. Cells were neutralized with 0.1M Sodium Borate (pH 8.5) for 3 minutes, washed four times with PBS and then blocked with PBS 2% BSA for 1 hour at room temperature. Cells were then incubated with murine anti-BrdU antibody monoclonal (1:100; Invitrogen, MoBU-1, stock 0.1mg/ml) in blocking buffer overnight in the cold room. Next day, hemocytes were washed twice with PBS 0.1% Tween 20 and incubated with Alexa 594 Goat-anti-mouse (1:2000) in blocking buffer for 2 hours at room temperature. Hemocytes were washed three times with blocking buffer and counterstained with 20 µM Hoechst 33342 (405, Molecular Probes, ThermoFisher Scientific, Waltham, MA, USA) and then mounted by adding 2 drops of Prolong Gold Antifade Mountant (Molecular Probes, ThermoFisher Scientific, Waltham, MA, USA).

For PHH3 staining, hemocytes were collected and fixed as described above. Following fixation, hemocytes were washed three times with PBS 0.1% triton and then blocked with PBS 2% BSA and 10% goat serum for 1-2 hours at room temperature. Hemocytes were then incubated with Anti-phospho-Histone H3 (Ser10) Antibody, Mitosis Marker (1:500, Millipore Sigma, # 06-570) in blocking buffer overnight in a cold room. Next day, hemocytes were washed with blocking buffer three times and then placed in a solution containing Alexa 594 goat anti-rabbit (1:2000) in blocking buffer for 2 hours at room temperature. Hemocytes were washed three times with PBS 0.1% triton and then counter stained with 1U of phalloidin (Alexa Fluor 488, Molecular Probes, ThermoFisher Scientific, Waltham, MA, USA) and 20 µM Hoechst 33342 (405, Molecular Probes, ThermoFisher Scientific, Waltham, MA, USA), both diluted in PBS 0.1% Triton at room temperature. Cells were then placed in mounting media for storage by adding 2 drops of Prolong Gold Antifade Mountant (Molecular Probes, ThermoFisher Scientific, Waltham, MA, USA).

### Oocyst counting in the midgut

*Plasmodium berghei* infections were evaluated by counting oocyst numbers after feeding on an infected mouse. Infected mosquitoes were kept at 20°C for 10 days after feeding when they were dissected, and their midgut fixed in 4% PFA for 15 minutes at room temperature. After washing with PBS three times, midguts were mounted in a slide and counted under a fluorescence microscope, where live oocysts were identified by their GFP expression.

