## Supplemental figures for "Hemocyte Differentiation to the Megacyte Lineage Enhances Mosquito Immunity Against *Plasmodium*"

**antiplasmodial immunity**

Ana Beatriz Barletta Ferreira^1^, Banhisikha Saha^1^, Nathanie Trisnadi^1#^, Octavio Talyuli^1,2^, Gianmarco Raddi^1#^, and Carolina Barillas-Mury^1*^.

^1^Laboratory of Malaria and Vector Research, National Institute of Allergy and Infectious Diseases, National Institutes of Health, Rockville, MD 20852.

^2^Instituto de Bioquímica Médica Leopoldo de Meis, Universidade Federal do Rio de Janeiro, Rio de Janeiro, Brazil.

^#^ Present address: Nathanie Trisnadi, Atropos Therapeutics Inc., San Carlos, California, USA. Gianmarco Raddi, School of Clinical Medicine, University of Cambridge, Cambridge CB2 0SP, UK, CRUK Cambridge Institute, Cambridge CB2 0RE, UK.

**Supplementary figures**

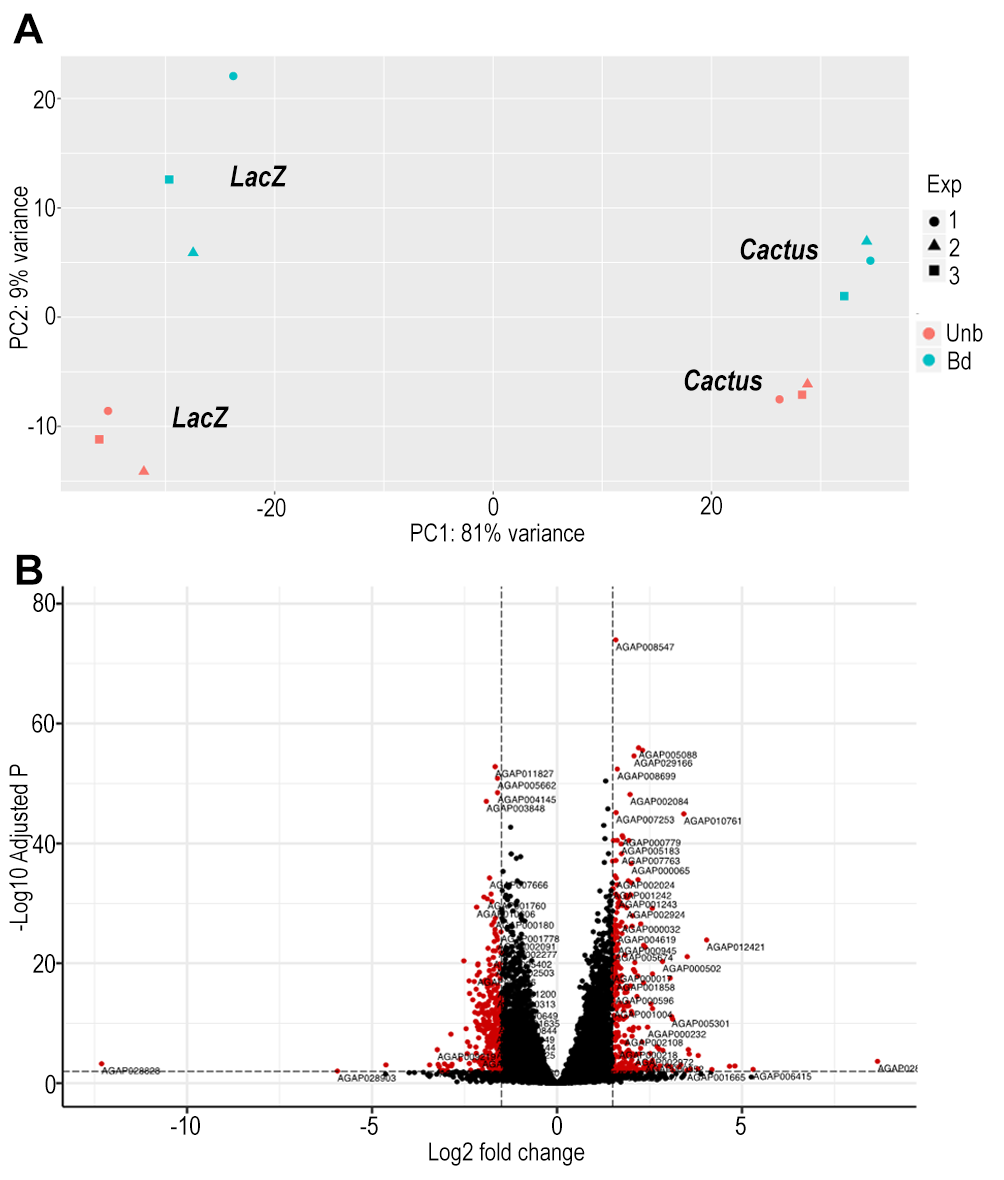

**Supplementary Figure 1. Quality control of dsCactus knockdown bulk RNA seq and Differential expression between bound and unbound fractions.** (A) PCA plot showing first and second principal component of dsRNA Cactus and control LacZ knock-down, for bound and unbound hemocyte fractions. Cactus and LacZ cluster separately, and so do bound and unbound hemocyte fractions. Bd = bound. UnB = unbound. One, two and three represent different biological replicates for each condition. (B) From a total of 9421 filtered genes (B) Volcano plot of DE genes between bound and unbound hemocytes filtered for log2 fold change >1.5 and Q-value <0.01.

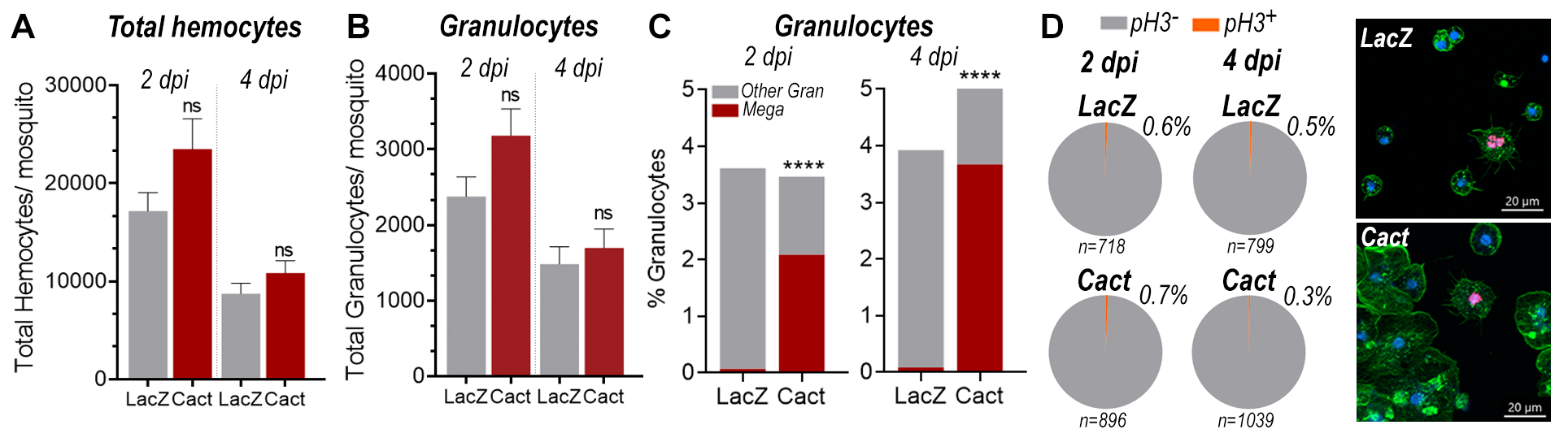

**Supplementary Figure 2. Megacyte increase in response to Cactus silencing is not a result of hemocyte proliferation.** (A) Total hemocyte counts per mosquito, at day 2 and 4 post dsRNA injection. (B) Granulocyte and megacyte percentages per mosquito against total hemocyte count, at day 2 and 4 post dsRNA injection. (C) Percentage of ph3^+^ cells among all granulocytes at day 2 and 4 post dsRNA injection. Actin is showing in green, nuclei in blue and ph3 in red. Scale Bar: 20um. Percentages were compared using X^2^-test, p=0.7746, nonsignificant for day 2 and p=0.4647, nonsignificant for day 4. Total hemocyte counts and percentages were calculated from 2 two independent experiments.

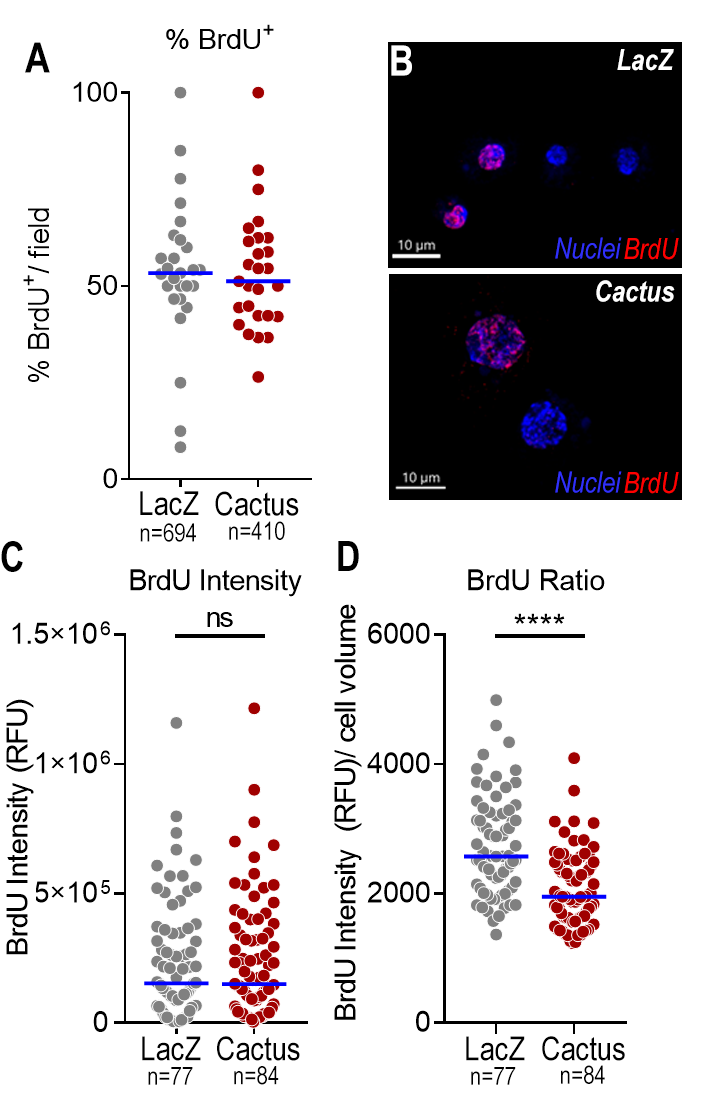

**Supplementary Figure 3. Toll activation controls megacyte differentiation and not proliferation.** (A) Percentage of BrdU^+^ hemocytes in dsLacZ and dsCactus mosquitoes 4 days after silencing. (B) Representative pictures of positive BrdU nuclei in LacZ and Cactus hemocytes. Nuclei is in Blue and BrdU is showing in Red. Scale Bar: 10um. (C) BrdU fluorescence intensity (Relative fluorescent units) in hemocytes from dsLacZ and dsCactus female mosquitoes. (D) Ratio between BrdU fluorescence intensity (Relative fluorescent units) and cell volume calculated based on the fluorescence of the nuclei (Hoechst staining). Quantification of cells was performed from 2 independent experiments. In each experiment 10 fields were collected, counted, and analyzed for BrdU staining. Blue bar in A, C and D represent medians. Mann-Whitney t-test, ****p<0.0001, ns – nonsignificant.

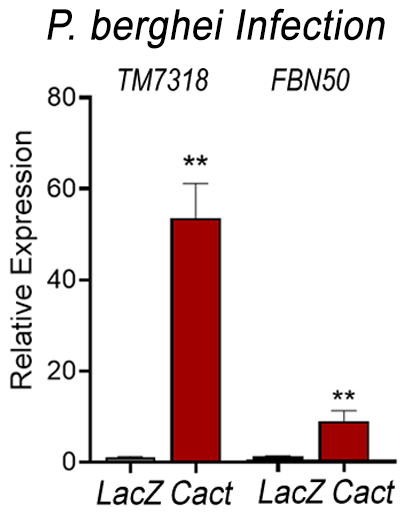

**Supplementary Figure 4. *Plasmodium berghei* infection increases megacyte association to the midgut basal surface.** Relative mRNA levels of effector hemocyte markers in the midgut 26 h post P.*berghei* infection (post-invasion) in LacZ and Cactus-silenced mosquitoes. TM7318, as a megacyte and FBN50, as an antimicrobial granulocyte marker. Error bars represent mean ± SEM. Unpaired t-test, *P≤0.05, **P≤0.01.

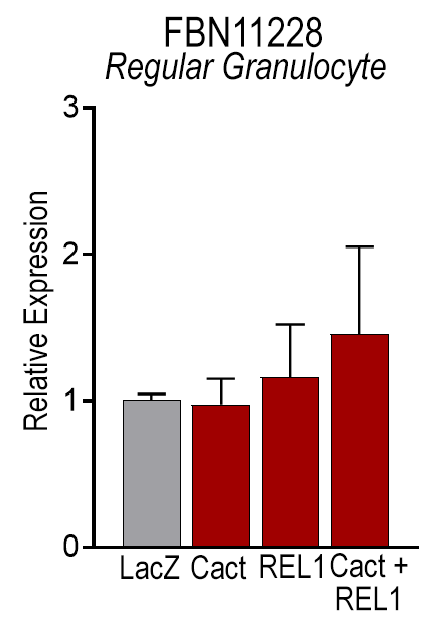

**Supplementary Figure 5. Regular granulocytes are not associated with the midgut basal lamina in response to *P.berghei* invasion in Cactus silenced mosquitoes.** Relative mRNA levels of FBN11228, a marker for regular granulocytes, in the midgut 26 h post *P*.*berghei* infection (post-invasion) in LacZ Cactus, REL-1, and Cactus + REL1- silenced mosquitoes. Error bars represent mean ± SEM. Unpaired t-test, **P≤0.01, ****P≤0.0001, ns - nonsignificant.

**Video S1. Top view (XY) of a regular granulocyte from *An.gambiae* mosquito female.** Showing in red is the microvesicle staining and in green the plasma membrane. Scale Bar: 10um. Hemocyte was imaged for 1 hour in intervals of 5 minutes.

**Video S2. Side view (XZ) of a regular granulocyte from An. gambiae mosquito female.** Showing in red is the microvesicle staining and in green the plasma membrane. Scale Bar: 5um. Hemocyte was imaged for 1 hour in intervals of 5 minutes.

**Video S3. Top view (XY) of a megacyte from An. gambiae mosquito female.** Showing in red is the microvesicle staining and in green the plasma membrane. Scale Bar: 10um. Hemocyte was imaged for 1 hour in intervals of 5 minutes.

**Video S4. Side view (XZ) of a megacyte from An. gambiae mosquito females** Showing in red is the microvesicle staining and in green the plasma membrane. Scale Bar: 5um. Hemocyte was imaged for 1 hour in intervals of 5 minutes.

**Video S5. *In vivo* hemocyte patrolling activity in *dsLacZ* mosquitoes.** Hemocytes stained in red were imaged through the cuticle of the mosquito for 1 hour and 20 minutes. Scale Bar: 30um.

**Video S6. *In vivo* hemocyte patrolling activity in *dsCactus* mosquitoes.** Hemocytes stained in red were imaged through the cuticle of the mosquito for 1 hour and 20 minutes. Scale Bar: 30um.

**Video S7.** ***In vitro* dynamics of *dsCactus* megacytes.** Perfused hemocytes from dsCactus mosquitoes. Plasma membrane is showing in green, microvesicles in red and nuclei in blue. Scale Bar: 20um.

**Table S1. List of upregulated genes in Cactus-silenced hemocytes**

**Table S2. List of downregulated genes in Cactus-silenced hemocytes**

**Table S3. List of Toll pathway components and final effectors upregulated in Cactus-silenced hemocytes**
