## Supplemental table S1 for "Hemocyte Differentiation to the Megacyte Lineage Enhances Mosquito Immunity Against *Plasmodium*"

| Upregulated genes by Cactus Knockdown | | | | | | | | | |
|---|---|---|---|---|---|---|---|---|---|
| Accesion Number | Name | Description | baseMean | log2FoldChange | lfcSE | stat | pvalue | padj | Hemocyte type |
| AGAP007343 | LYSC2 | C-type lysozyme | 150003.6078 | 11.38985051 | 0.890888229 | 12.78482547 | 1.9931E-37 | 3.07315E-36 | |
| AGAP011920 | AGAP011920 | eupolytin | 147.2376396 | 10.24020054 | 0.850232306 | 12.04400311 | 2.08583E-33 | 2.77551E-32 | |
| AGAP002842 | AGAP002842 | CLIPD1 protein | 908.0228542 | 9.35826901 | 0.461470828 | 20.27922121 | 1.96213E-91 | 1.31101E-89 | |
| AGAP006674 | AGAP006674 | chymotrypsin-like protease (Precursor) | 162.6513041 | 9.223197473 | 1.269757003 | 7.263750031 | 3.76504E-13 | 1.82743E-12 | |
| AGAP000188 | AGAP000188 | Uridine phosphorylase [Source:UniProtKB/TrEMBL;Acc:Q7PFX7] | 35514.58459 | 8.223334693 | 0.206853775 | 39.75433709 | 0 | 0 | |
| AGAP010771 | AGAP010771 | nan | 212.6244713 | 7.835796021 | 0.779798413 | 10.04848932 | 9.32865E-24 | 8.12248E-23 | |
| AGAP028044 | CPLCP23 | cuticular protein (putative) CPLCP23 | 568.6912146 | 7.782049203 | 0.423632051 | 18.36983107 | 2.29124E-75 | 1.07392E-73 | |
| AGAP028137 | CPLCP17 | cuticular protein (putative) CPLCP17 | 23.15155164 | 7.748117482 | 0.894497512 | 8.661977679 | 4.63649E-18 | 3.05885E-17 | |
| AGAP009728 | AGAP009728 | ficolin | 96.0328881 | 7.552158704 | 0.734283475 | 10.2850724 | 8.22802E-25 | 7.47504E-24 | |
| AGAP012936 | nan | nan | 771.7781334 | 7.376956435 | 0.577741905 | 12.76860197 | 2.45525E-37 | 3.76725E-36 | |
| AGAP010256 | AGAP010256 | nan | 24.41954606 | 6.826258554 | 0.892933963 | 7.644751838 | 2.09349E-14 | 1.10678E-13 | |
| AGAP004497 | AGAP004497 | nan | 140.0929815 | 6.723899197 | 0.776063174 | 8.664113205 | 4.55041E-18 | 3.00416E-17 | |
| AGAP009489 | AGAP009489 | Cytidine deaminase [Source:UniProtKB/TrEMBL;Acc:Q7QG43] | 26454.65472 | 6.710199366 | 0.127644774 | 52.56932305 | 0 | 0 | |
| AGAP002007 | AGAP002007 | reticulon/nogo receptor | 10.82279296 | 6.638250029 | 0.923462513 | 7.188434759 | 6.55383E-13 | 3.1326E-12 | |
| AGAP029133 | AGAP029133 | nan | 150.5264926 | 6.565219163 | 0.474005189 | 13.8505217 | 1.26291E-43 | 2.4232E-42 | |
| AGAP011980 | AGAP011980 | nan | 120.7652387 | 6.522865768 | 0.556577551 | 11.71959911 | 1.0115E-31 | 1.25882E-30 | |
| AGAP008341 | AGAP008341 | serine/threonine-protein kinase Chk2 | 412.0975372 | 6.519154025 | 0.327414123 | 19.9110349 | 3.265E-88 | 2.10682E-86 | |
| AGAP028008 | CPLCP19 | cuticular protein (putative) CPLCP19 | 26.32277697 | 6.433828174 | 0.790844224 | 8.13539251 | 4.10606E-16 | 2.42528E-15 | |
| AGAP004115 | AGAP004115 | cystinosin | 4407.482278 | 6.413747248 | 0.197665013 | 32.44755943 | 5.8634E-231 | 3.0689E-228 | |
| AGAP007150 | AGAP007150 | nan | 11.35455667 | 6.222639299 | 0.900634018 | 6.909176395 | 4.87476E-12 | 2.19108E-11 | |
| AGAP001002 | AGAP001002 | Toll protein | 116127.845 | 6.177628682 | 0.268236101 | 23.03056397 | 2.3037E-117 | 2.7825E-115 | Megacytes |
| AGAP013210 | AGAP013210 | Down syndrome cell adhesion molecule-like protein 1 | 104.7167101 | 6.177452535 | 0.853967007 | 7.233830448 | 4.69558E-13 | 2.2674E-12 | |
| AGAP000473 | AGAP000473 | Aa_trans domain-containing protein [Source:UniProtKB/TrEMBL;Acc | 32.72500638 | 6.030795595 | 0.694109812 | 8.688532407 | 3.67151E-18 | 2.43244E-17 | |
| AGAP011226 | AGAP011226 | nan | 2959.788347 | 6.023113599 | 0.218447016 | 27.57242332 | 2.3834E-167 | 5.2219E-165 | Megacytes |
| AGAP028013 | CPLCP25 | cuticular protein (putative) CPLCP25 | 9.869237005 | 5.98271328 | 0.911017367 | 6.567068309 | 5.13155E-11 | 2.15342E-10 | |
| AGAP028178 | CPLCP13 | cuticular protein (putative) CPLCP13 | 9.845746458 | 5.903622512 | 0.934991859 | 6.314089748 | 2.71756E-10 | 1.07617E-09 | |
| AGAP012544 | CASPS14 | short caspase 14 | 9.247569773 | 5.88732349 | 0.910161253 | 6.468440042 | 9.90198E-11 | 4.0489E-10 | |
| AGAP006982 | AGAP006982 | nan | 7.337402519 | 5.814868909 | 0.975476518 | 5.961054728 | 2.50615E-09 | 9.02885E-09 | |
| AGAP013142 | AGAP013142 | nan | 213.6895269 | 5.786289236 | 0.403777627 | 14.33038596 | 1.41333E-46 | 2.93929E-45 | |
| AGAP028156 | CPLCP14 | cuticular protein (putative) CPLCP14 | 6.344508639 | 5.780752493 | 0.982548003 | 5.883430098 | 4.0185E-09 | 1.42164E-08 | |
| AGAP006368 | Obp69 | odorant-binding protein 69 | 42.20555312 | 5.74398315 | 0.556557706 | 10.32055273 | 5.68947E-25 | 5.26527E-24 | |
| AGAP001508 | AGAP001508 | nan | 5677.718823 | 5.67371698 | 0.63276711 | 8.966516896 | 3.06039E-19 | 2.14843E-18 | |
| AGAP010833 | CLIPB14 | CLIP-domain serine protease | 350.1728028 | 5.609302809 | 0.696698509 | 8.051262835 | 8.19441E-16 | 4.75953E-15 | |
| AGAP005101 | AGAP005101 | nan | 828.5282682 | 5.561330369 | 0.314177259 | 17.70125053 | 4.10099E-70 | 1.6798E-68 | |
| AGAP009918 | AGAP009918 | SGS5 [Source:UniProtKB/TrEMBL;Acc:Q5XLG6] | 7.330305498 | 5.546120623 | 0.960926948 | 5.771636057 | 7.85055E-09 | 2.69436E-08 | |
| AGAP009917 | AGAP009917 | SGS4 [Source:UniProtKB/TrEMBL;Acc:Q5XLG7] | 7.641458514 | 5.501527096 | 0.991755504 | 5.54726147 | 2.90179E-08 | 9.41058E-08 | |
| AGAP005072 | AGAP005072 | nan | 28.87051263 | 5.47858478 | 0.747639945 | 7.327838506 | 2.33894E-13 | 1.14528E-12 | |
| AGAP011232 | AGAP011232 | nan | 5.109307773 | 5.472165276 | 0.983504031 | 5.56394799 | 2.63739E-08 | 8.59751E-08 | |
| AGAP009916 | nan | nan | 2842.248885 | 5.410882466 | 0.26518758 | 20.4039815 | 1.5414E-92 | 1.07567E-90 | |
| AGAP010811 | AGAP010811 | Fibrinogen C-terminal domain-containing protein [Source:UniProtKB | 91.01407029 | 5.256865908 | 0.796529885 | 6.599709572 | 4.11964E-11 | 1.73885E-10 | |
| AGAP028060 | AGAP028060 | glucuronosyltransferase | 887.3372409 | 5.228176685 | 0.245624618 | 21.28523083 | 1.5558E-100 | 1.30867E-98 | |
| AGAP028032 | AGAP028032 | nan | 11.14647247 | 5.22662839 | 0.835531452 | 6.255453793 | 3.96362E-10 | 1.552E-09 | |
| AGAP013201 | AGAP013201 | nan | 32.2763464 | 5.159542477 | 0.608926828 | 8.473173193 | 2.38797E-17 | 1.51089E-16 | |
| AGAP011578 | nan | nan | 408.1797303 | 5.113309696 | 0.232704283 | 21.97342322 | 5.172E-107 | 5.1835E-105 | |
| AGAP010398 | AGAP010398 | Flavin-containing monooxygenase FMO GS-OX-like 1 | 2219.565421 | 5.04210753 | 0.295949076 | 17.03707813 | 4.35976E-65 | 1.51005E-63 | |
| AGAP000745 | AGAP000745 | nan | 12.33594826 | 5.017490393 | 0.784368753 | 6.396851448 | 1.58614E-10 | 6.39135E-10 | |
| AGAP013059 | AGAP013059 | nan | 312.6951851 | 4.983082365 | 0.225695342 | 22.07880018 | 5.0532E-108 | 5.2315E-106 | |
| AGAP008235 | AGAP008235 | nan | 4.826954054 | 4.979070018 | 0.993874755 | 5.009755999 | 5.44991E-07 | 1.56297E-06 | |
| AGAP008851 | AGAP008851 | E3 ubiquitin-protein ligase mind-bomb [Source:UniProtKB/TrEMBL;A | 12042.35188 | 4.958656949 | 0.145571218 | 34.06344342 | 2.5668E-254 | 1.8601E-251 | |
| AGAP013480 | AGAP013480 | nan | 3.590374652 | 4.891248243 | 1.070640044 | 4.568527277 | 4.91163E-06 | 1.27192E-05 | |
| AGAP011576 | nan | nan | 172.8423044 | 4.884975144 | 0.273106829 | 17.88668252 | 1.49763E-71 | 6.327E-70 | |
| AGAP028432 | AGAP028432 | nan | 29.99478111 | 4.852627814 | 0.697149222 | 6.960673072 | 3.38651E-12 | 1.54351E-11 | |
| AGAP028138 | AGAP028138 | nan | 5559.34555 | 4.838569559 | 0.154181625 | 31.38227105 | 3.532E-216 | 1.4467E-213 | |
| AGAP002291 | AGAP002291 | ebony | 1221.910971 | 4.821937032 | 0.298908838 | 16.1317981 | 1.52515E-58 | 4.64997E-57 | |
| AGAP001767 | AGAP001767 | Peptidase (mitochondrial processing) beta | 29.30456531 | 4.808103681 | 0.539067204 | 8.919302909 | 4.69232E-19 | 3.25526E-18 | |
| AGAP004382 | GSTD3 | glutathione S-transferase delta class 3 | 301.6395523 | 4.782964611 | 0.392783217 | 12.17711043 | 4.11608E-34 | 5.64448E-33 | |
| AGAP001763 | Fatp | fatty acid transporter protein 1 | 1430.266221 | 4.763243002 | 0.168815063 | 28.21574645 | 3.7483E-175 | 9.5439E-173 | |
| AGAP029077 | AGAP029077 | nan | 20.54793606 | 4.731097678 | 0.610346367 | 7.75149642 | 9.08159E-15 | 4.96274E-14 | |
| AGAP007216 | AGAP007216 | DE-cadherin | 328.681905 | 4.712111296 | 0.212535955 | 22.17089008 | 6.56E-109 | 6.8669E-107 | |
| AGAP004316 | AGAP004316 | nan | 4851.502155 | 4.668543972 | 0.296069202 | 15.76842148 | 5.13164E-56 | 1.44746E-54 | |
| AGAP012581 | AGAP012581 | nan | 30.44937246 | 4.647732107 | 0.491782187 | 9.450793934 | 3.3627E-21 | 2.61602E-20 | |
| AGAP007666 | AGAP007666 | calcyphosin-like protein | 10259.11802 | 4.629661893 | 0.250532256 | 18.47930466 | 3.03062E-76 | 1.47935E-74 | |
| AGAP012529 | GALE8 | galectin 8 | 209.4838637 | 4.614028629 | 0.248138641 | 18.59455914 | 3.55625E-77 | 1.80126E-75 | |
| AGAP001753 | AGAP001753 | 15-hydroxyprostaglandin dehydrogenase (NAD) | 34.74159678 | 4.54898947 | 0.497561631 | 9.142564826 | 6.09884E-20 | 4.46443E-19 | |
| AGAP005205 | PGRPLA | peptidoglycan recognition protein (long) | 4109.111356 | 4.519663807 | 0.212616761 | 21.257326 | 2.8204E-100 | 2.35137E-98 | |
| AGAP011225 | AGAP011225 | nan | 57.3460484 | 4.513355285 | 0.364459668 | 12.38368927 | 3.20275E-35 | 4.57862E-34 | |
| AGAP010385 | AGAP010385 | nan | 86.07016609 | 4.50518505 | 0.344460126 | 13.07897406 | 4.34262E-39 | 7.05376E-38 | |
| AGAP010390 | COE12O | carboxylesterase | 1082.615231 | 4.50345105 | 0.216708186 | 20.78117642 | 6.40693E-96 | 4.79045E-94 | |
| AGAP007684 | AGAP007684 | Tubulointerstitial nephritis antigen | 2986.014541 | 4.500908187 | 0.167961969 | 26.79718631 | 3.4847E-158 | 6.6998E-156 | |
| AGAP013179 | AGAP013179 | nan | 1393.535639 | 4.500873941 | 0.178477963 | 25.21809346 | 2.5367E-140 | 3.9831E-138 | |
| AGAP001509 | AGAP001509 | nan | 15.66755972 | 4.473477738 | 0.763041733 | 5.862690789 | 4.55426E-09 | 1.60515E-08 | |
| AGAP006345 | AGAP006345 | nan | 50.6959971 | 4.40142724 | 0.408312501 | 10.77955544 | 4.29959E-27 | 4.40287E-26 | |
| AGAP001679 | nan | nan | 75.51096568 | 4.393964211 | 0.371476076 | 11.82839082 | 2.78428E-32 | 3.56396E-31 | |
| AGAP013382 | AGAP013382 | nan | 4.610872597 | 4.379260006 | 0.988667687 | 4.42945599 | 9.44711E-06 | 2.36391E-05 | |
| AGAP000667 | AGAP000667 | G_PROTEIN_RECEP_F1_2 domain-containing protein [Source:UniPro | 10.966911 | 4.375489321 | 0.821273441 | 5.327688808 | 9.94704E-08 | 3.06145E-07 | |
| AGAP000133 | AGAP000133 | nan | 26.74337643 | 4.372513068 | 0.550486435 | 7.942998761 | 1.97351E-15 | 1.11868E-14 | |
| AGAP029075 | AGAP029075 | nan | 6.608415866 | 4.331558838 | 0.97505347 | 4.442380822 | 8.89689E-06 | 2.23394E-05 | |
| AGAP001061 | AGAP001061 | nan | 12.8658239 | 4.323205349 | 0.69351603 | 6.233749703 | 4.55401E-10 | 1.7736E-09 | |
| AGAP001705 | AGAP001705 | nan | 598.5040026 | 4.270835173 | 0.189637716 | 22.52102205 | 2.5834E-112 | 2.7657E-110 | |
| AGAP002232 | GPR5HT2A | putative serotonin 5HT-2a receptor | 4.627635348 | 4.25586234 | 0.980578749 | 4.340153551 | 1.42383E-05 | 3.47781E-05 | |
| AGAP010818 | AGAP010818 | nan | 263.0759335 | 4.253454758 | 0.304341039 | 13.97594874 | 2.18581E-44 | 4.25466E-43 | |
| AGAP006367 | AGAP006367 | nan | 7339.224631 | 4.186676762 | 0.12803169 | 32.70031628 | 1.5457E-234 | 9.1011E-232 | Prohemocytes and Megacytes |
| AGAP028652 | AGAP028652 | nan | 83.33450069 | 4.180241485 | 0.305182657 | 13.69750671 | 1.05071E-42 | 1.92957E-41 | |
| AGAP013194 | AGAP013194 | nan | 26.73183862 | 4.179948626 | 0.514950267 | 8.117188975 | 4.77106E-16 | 2.79702E-15 | |
| AGAP028742 | AGAP028742 | nan | 5.526065886 | 4.171738628 | 0.936050441 | 4.456745541 | 8.32133E-06 | 2.09781E-05 | |
| AGAP010816 | TEP3 | thioester-containing protein 3 | 62660.7236 | 4.153502512 | 0.220728572 | 18.81724002 | 5.45532E-79 | 2.90365E-77 | AM granulocytes |
| AGAP010709 | CTL2 | C-type lectin (CTL) | 6.962910679 | 4.149323731 | 0.815225457 | 5.089786753 | 3.58466E-07 | 1.05108E-06 | |
| AGAP012787 | AGAP012787 | nan | 4.25996281 | 4.141323136 | 1.098656544 | 3.769442923 | 0.000163612 | 0.000351819 | |
| AGAP001004 | TOLL1A | TOLL-like receptor 1A | 757.4906654 | 4.105939711 | 0.211676876 | 19.39720481 | 8.1483E-84 | 4.95259E-82 | |
| AGAP004181 | AGAP004181 | Fibroblast growth factor [Source:UniProtKB/TrEMBL;Acc:A0NDD8] | 108.3931144 | 4.082026573 | 0.282219388 | 14.46401892 | 2.045E-47 | 4.37863E-46 | |
| AGAP012748 | AGAP012748 | nan | 226.2312456 | 4.076905591 | 0.198811373 | 20.50640021 | 1.88758E-93 | 1.34719E-91 | |
| AGAP001048 | AGAP001048 | nan | 8.569616286 | 4.073769288 | 0.777939014 | 5.236617796 | 1.63546E-07 | 4.9431E-07 | |
| AGAP009590 | AGAP009590 | amiloride-sensitive sodium channel, other | 25.08986149 | 4.027708717 | 0.489815124 | 8.222916195 | 1.98614E-16 | 1.19029E-15 | |
| AGAP010617 | AGAP010617 | nan | 5.187413678 | 4.014771016 | 1.081080162 | 3.713666346 | 0.000204278 | 0.000434229 | |
| AGAP010869 | AGAP010869 | fibrinogen | 211.8191342 | 4.01427125 | 0.504674447 | 7.954179717 | 1.80322E-15 | 1.0271E-14 | |
| AGAP002591 | AGAP002591 | nan | 68.18572522 | 4.009709645 | 0.350556533 | 11.43812558 | 2.69648E-30 | 3.21564E-29 | |
| AGAP028424 | AGAP028424 | nan | 12.44339884 | 3.950739816 | 0.655786976 | 6.024425553 | 1.69711E-09 | 6.2309E-09 | |
| AGAP012172 | nan | nan | 61.20090705 | 3.942407848 | 0.357271542 | 11.03476597 | 2.59727E-28 | 2.80606E-27 | |
| AGAP004170 | AGAP004170 | nan | 861.686701 | 3.936937407 | 0.203077887 | 19.3863422 | 1.00645E-83 | 6.07804E-82 | |
| AGAP028574 | AGAP028574 | nan | 25.15266406 | 3.914226008 | 0.46645047 | 8.391514769 | 4.79918E-17 | 2.9765E-16 | |
| AGAP012011 | AGAP012011 | nan | 608.3912712 | 3.909853698 | 0.186469643 | 20.96777594 | 1.29162E-97 | 9.893E-96 | |
| AGAP004837 | nan | nan | 911.9044456 | 3.905203598 | 0.209913931 | 18.60383244 | 2.99138E-77 | 1.52334E-75 | |
| AGAP010387 | AGAP010387 | alanine-glyoxylate aminotransferase | 208.9900095 | 3.898506459 | 0.71001505 | 5.490737779 | 4.00258E-08 | 1.28566E-07 | |
| AGAP000835 | AGAP000835 | nan | 14.19801452 | 3.891478334 | 0.789899311 | 4.92654985 | 8.36943E-07 | 2.35298E-06 | |
| AGAP013005 | AGAP013005 | nan | 48169.71659 | 3.889840306 | 0.269699084 | 14.42289031 | 3.71455E-47 | 7.88171E-46 | |
| AGAP009155 | AGAP009155 | nan | 28.75046445 | 3.875752747 | 0.429856158 | 9.016394617 | 1.94381E-19 | 1.38105E-18 | |
| AGAP012171 | nan | nan | 322.3687888 | 3.864774626 | 0.211464148 | 18.27626418 | 1.27892E-74 | 5.87741E-73 | |
| AGAP011654 | AGAP011654 | membrane dipeptidase | 317.542632 | 3.864620461 | 0.36441992 | 10.60485514 | 2.82902E-26 | 2.77338E-25 | |
| AGAP006520 | AGAP006520 | Fatty acid elongase 2 | 285.4705459 | 3.795738454 | 0.20123076 | 18.86261554 | 2.3148E-79 | 1.25332E-77 | |
| AGAP010855 | AGAP010855 | solute carrier family 45, member 1/2/4 | 26.94987032 | 3.767997156 | 0.456747421 | 8.249629853 | 1.58886E-16 | 9.55852E-16 | |
| AGAP010542 | dpr6 | defective proboscis extension response 6 | 13.87282103 | 3.765043673 | 0.579086417 | 6.501695713 | 7.94196E-11 | 3.26588E-10 | |
| AGAP006986 | AGAP006986 | nan | 63.95069943 | 3.760568818 | 0.528657152 | 7.113435999 | 1.13189E-12 | 5.30788E-12 | |
| AGAP010851 | GPRLKK | putative leukokinin receptor | 19.02684305 | 3.753645357 | 0.506382831 | 7.412663164 | 1.23788E-13 | 6.19006E-13 | |
| AGAP009050 | AGAP009050 | cyclic nucleotide gated channel, invertebrate | 4.971353142 | 3.751865174 | 0.968715484 | 3.873031076 | 0.00010749 | 0.000236936 | |
| AGAP004574 | AGAP004574 | nan | 181.2982478 | 3.734529071 | 0.27437419 | 13.61108009 | 3.4412E-42 | 6.22257E-41 | |
| AGAP010240 | ESP | late trypsin | 7.684845367 | 3.70010401 | 0.72645855 | 5.093344983 | 3.51801E-07 | 1.0325E-06 | |
| AGAP012588 | AGAP012588 | nan | 4.070845174 | 3.699617854 | 0.907309853 | 4.077568256 | 4.55092E-05 | 0.000104929 | |
| AGAP006449 | AGAP006449 | nan | 6.577349064 | 3.697748192 | 0.773326834 | 4.781611122 | 1.73896E-06 | 4.72397E-06 | |
| AGAP005949 | AGAP005949 | 2-acylglycerol O-acyltransferase 2 | 1869.718015 | 3.689835743 | 0.154001684 | 23.9597103 | 7.3195E-127 | 9.9938E-125 | |
| AGAP001376 | SRPN17 | serine protease inhibitor (serpin) 17 | 2134.989079 | 3.685525524 | 0.244477096 | 15.07513619 | 2.36029E-51 | 5.67252E-50 | |
| AGAP009467 | ABCG16 | ATP-binding cassette transporter (ABC transporter) family G membe | 162.6789378 | 3.658598516 | 0.277057569 | 13.20519244 | 8.18902E-40 | 1.36065E-38 | |
| AGAP007356 | AGAP007356 | CRAL-TRIO domain-containing protein [Source:UniProtKB/TrEMBL;A | 7.428289046 | 3.650557961 | 0.796475364 | 4.583390935 | 4.57496E-06 | 1.1899E-05 | |
| AGAP011355 | AGAP011355 | nan | 1146.688915 | 3.632880017 | 0.329790955 | 11.01570544 | 3.21012E-28 | 3.44841E-27 | |
| AGAP010294 | AGAP010294 | nan | 8.978569735 | 3.631947615 | 0.691394661 | 5.253074433 | 1.49581E-07 | 4.53557E-07 | |
| AGAP003627 | AGAP003627 | nan | 312.5971806 | 3.591767533 | 0.332455278 | 10.80376151 | 3.30385E-27 | 3.39799E-26 | |
| AGAP007858 | AGAP007858 | lysyl-tRNA synthetase, class II | 435.1658098 | 3.585353666 | 0.198879698 | 18.02775091 | 1.17995E-72 | 5.17035E-71 | |
| AGAP007262 | AGAP007262 | chymotrypsin-like protease | 3.017637292 | 3.583170245 | 1.00830931 | 3.553641932 | 0.000379936 | 0.00077392 | |
| AGAP010759 | AGAP010759 | nan | 9550.050842 | 3.581034892 | 0.227030083 | 15.77339376 | 4.74312E-56 | 1.34189E-54 | Megacytes |
| AGAP028541 | AGAP028541 | nan | 41.48900921 | 3.576155379 | 0.389605454 | 9.178915079 | 4.35456E-20 | 3.20503E-19 | |
| AGAP005285 | AGAP005285 | nan | 388.9666566 | 3.566213826 | 0.164646555 | 21.65981441 | 4.9121E-104 | 4.5819E-102 | |
| AGAP010760 | AGAP010760 | nan | 12109.61415 | 3.52948473 | 0.223693646 | 15.77820738 | 4.39494E-56 | 1.24713E-54 | |
| AGAP005742 | AGAP005742 | semaphorin | 54.73561392 | 3.529220116 | 0.331256398 | 10.65404362 | 1.66952E-26 | 1.65216E-25 | |
| AGAP003139 | SRPN9 | serine protease inhibitor (serpin) 9 | 3098.83565 | 3.528515234 | 0.1217241 | 28.98781114 | 9.3726E-185 | 2.6757E-182 | |
| AGAP013487 | AGAP013487 | nan | 95.5750241 | 3.52552181 | 0.378350519 | 9.318136572 | 1.18401E-20 | 8.96672E-20 | |
| AGAP006985 | AGAP006985 | nan | 98.6841087 | 3.525303596 | 0.447248609 | 7.882201363 | 3.21663E-15 | 1.80273E-14 | |
| AGAP007318 | TM7318 | Transmembrane protein - megacyte marker | 7478.159662 | 3.519945262 | 0.22981957 | 15.31612502 | 5.96704E-53 | 1.52346E-51 | Megacytes |

| | | | | | | | | | |
|---|---|---|---|---|---|---|---|---|---|
| AGAP006343 | PGRPS2 | peptidoglycan recognition protein (short) | 19.36251327 | 3.509206816 | 0.504347815 | 6.957910224 | 3.45357E-12 | 1.57179E-11 | |
| AGAP005756 | COEAE1D | carboxylesterase | 287.5124919 | 3.505892189 | 0.253240293 | 13.84413257 | 1.38037E-43 | 2.63248E-42 | |
| AGAP003892 | AGAP003892 | solute carrier family 29 (equilibrative nucleoside transporter), memb | 7735.653319 | 3.505463334 | 0.180513216 | 19.41942763 | 5.28759E-84 | 3.2347E-82 | |
| AGAP028094 | AGAP028094 | nan | 102.5874086 | 3.482743526 | 0.328566226 | 10.59982203 | 2.98548E-26 | 2.92372E-25 | |
| AGAP012321 | OBP26 | odorant-binding protein 26 | 8.069470565 | 3.459364058 | 0.707432684 | 4.890025776 | 1.00823E-06 | 2.80855E-06 | |
| AGAP000976 | AGAP000976 | nan | 20.96702416 | 3.456514724 | 0.483776496 | 7.144858731 | 9.00885E-13 | 4.26923E-12 | |
| AGAP006603 | AGAP006603 | nan | 253.6852551 | 3.454719664 | 0.226954278 | 15.22209536 | 2.52318E-52 | 6.35586E-51 | |
| AGAP001652 | AGAP001652 | lipase | 33397.88846 | 3.446083482 | 0.120094935 | 28.69466134 | 4.4477E-181 | 1.2324E-178 | Megacytes |
| AGAP005440 | AGAP005440 | nan | 11.53728419 | 3.441455479 | 0.637010503 | 5.402509792 | 6.57149E-08 | 2.06298E-07 | |
| AGAP010762 | nan | nan | 723.8389244 | 3.438150439 | 0.21566802 | 15.94186494 | 3.24522E-57 | 9.55415E-56 | |
| AGAP010675 | LRIM18 | leucine-rich immune protein (Coil-less) | 554.0630047 | 3.394752947 | 0.219684695 | 15.45284232 | 7.21887E-54 | 1.88914E-52 | |
| AGAP008100 | AGAP008100 | Spire [Source:UniProtKB/TrEMBL;Acc:A0A1S4GX95] | 465.3383139 | 3.388809853 | 0.215974124 | 15.69081421 | 1.74799E-55 | 4.83674E-54 | |
| AGAP011063 | AGAP011063 | nan | 11.66403047 | 3.376061995 | 0.565508515 | 5.969957847 | 2.37315E-09 | 8.58251E-09 | |
| AGAP013117 | AGAP013117 | nan | 4832.055875 | 3.37436503 | 0.226525766 | 14.89616429 | 3.49051E-50 | 8.05982E-49 | |
| AGAP002643 | AGAP002643 | nan | 8.926780993 | 3.366014173 | 0.953295628 | 3.53092375 | 0.000414111 | 0.000838998 | |
| AGAP008403 | AGAP008403 | nan | 307.8027194 | 3.346911517 | 0.247522516 | 13.5216447 | 1.16541E-41 | 2.05605E-40 | |
| AGAP001005 | AGAP001005 | nan | 10.89803658 | 3.344267139 | 0.744617998 | 4.491252087 | 7.08057E-06 | 1.79898E-05 | |
| AGAP003318 | AGAP003318 | nan | 146.408358 | 3.33645302 | 0.220070341 | 15.16084812 | 6.42345E-52 | 1.58003E-50 | |
| AGAP003319 | AGAP003319 | nan | 10632.10269 | 3.327043033 | 0.13845288 | 24.03014675 | 1.3465E-127 | 1.8655E-125 | Megacytes |
| AGAP028531 | AGAP028531 | nan | 83.38038377 | 3.324423113 | 0.237709119 | 13.98525695 | 1.91783E-44 | 3.74076E-43 | |
| AGAP009424 | AGAP009424 | nan | 709.9146119 | 3.314407578 | 0.138854208 | 23.86969491 | 6.3243E-126 | 8.2752E-124 | |
| AGAP007290 | AGAP007290 | nan | 35.66227852 | 3.311507228 | 0.513620037 | 6.447387149 | 1.13795E-10 | 4.63494E-10 | |
| AGAP003626 | AGAP003626 | nan | 86.15910223 | 3.304448947 | 0.291439478 | 11.33837105 | 8.47065E-30 | 9.80368E-29 | |
| AGAP013166 | Cht5-1 | chitinase | 464.379508 | 3.295251224 | 0.327529936 | 10.06091615 | 8.22295E-24 | 7.19967E-23 | |
| AGAP001648 | CLIPB17 | CLIP-domain serine protease | 9264.846222 | 3.29031506 | 0.30899714 | 10.64836735 | 1.7745E-26 | 1.7542E-25 | |
| AGAP004243 | AGAP004243 | nan | 16.50408841 | 3.289421142 | 0.589621426 | 5.57886976 | 2.42086E-08 | 7.92734E-08 | |
| AGAP004920 | CASPS6 | short caspase 6 | 720.3876092 | 3.26116271 | 0.175039789 | 18.63097946 | 1.80196E-77 | 9.27666E-76 | |
| AGAP010027 | AGAP010027 | nan | 15.31748505 | 3.253123128 | 0.495850555 | 6.560692722 | 5.35584E-11 | 2.24056E-10 | |
| AGAP006641 | nan | nan | 46.0072813 | 3.252011142 | 0.293969843 | 11.06239712 | 1.90926E-28 | 2.07464E-27 | |
| AGAP004309 | AGAP004309 | solute carrier family 22 | 1105.673549 | 3.237728565 | 0.138468626 | 23.38239827 | 6.4555E-121 | 8.0022E-119 | |
| AGAP011667 | AGAP011667 | nan | 17.1722601 | 3.202296789 | 0.615962563 | 5.198849706 | 2.00526E-07 | 6.01641E-07 | |
| AGAP001704 | AGAP001704 | isopentenyl-diphosphate delta-isomerase | 1108.654866 | 3.194506374 | 0.094113214 | 33.94322909 | 1.5353E-252 | 1.0331E-249 | |
| AGAP011560 | AGAP011560 | nan | 7031.969228 | 3.186375928 | 0.119727501 | 26.61356748 | 4.7288E-156 | 8.25E-154 | |
| AGAP005755 | AGAP005755 | DNA-binding protein D-ETS-3 | 17.09583769 | 3.167032606 | 0.487523736 | 6.496160843 | 8.23956E-11 | 3.38382E-10 | |
| AGAP003496 | AGAP003496 | nan | 11011.69344 | 3.154169972 | 0.118490752 | 26.61954549 | 4.0323E-156 | 7.1676E-154 | |
| AGAP006689 | AGAP006689 | BTB (POZ) domain containing 9 | 269.5604966 | 3.138907034 | 0.204997953 | 15.31189453 | 6.36816E-53 | 1.62147E-51 | |
| AGAP009490 | AGAP009490 | Necdin-like 2 | 456.0357986 | 3.135336635 | 0.100399436 | 31.22862801 | 4.3556E-214 | 1.7097E-211 | |
| AGAP004335 | AGAP004335 | filamin | 17572.09317 | 3.132381678 | 0.143729188 | 21.79363651 | 2.6661E-105 | 2.6164E-103 | |
| AGAP006642 | nan | nan | 158.541787 | 3.122376973 | 0.201035871 | 15.53144199 | 2.12546E-54 | 5.65649E-53 | |
| AGAP028005 | AGAP028005 | nan | 24.09103098 | 3.111365845 | 0.431623752 | 7.208513975 | 5.65658E-13 | 2.71614E-12 | |
| AGAP003839 | AGAP003839 | Fem-1 homolog c | 2390.28197 | 3.10470337 | 0.075717415 | 41.0038216 | 0 | 0 | |
| AGAP008671 | AGAP008671 | nan | 138.1995901 | 3.09377269 | 0.215079967 | 14.38429032 | 6.49408E-47 | 1.3626E-45 | |
| AGAP000235 | AGAP000235 | Thymosin | 39370.31323 | 3.090485049 | 0.078616245 | 39.31102347 | 0 | 0 | Megacytes |
| AGAP002677 | AGAP002677 | coiled-coil domain-containing protein lobo homolog | 5.832111103 | 3.090367103 | 0.726377629 | 4.254491025 | 2.09525E-05 | 5.01126E-05 | |
| AGAP029047 | CTL5 | C-type lectin | 8.473021694 | 3.083190214 | 0.680354664 | 4.531739657 | 5.84999E-06 | 1.50335E-05 | |
| AGAP013197 | AGAP013197 | nan | 16.19845417 | 3.076212214 | 0.468490176 | 6.566225652 | 5.16066E-11 | 2.16371E-10 | |
| AGAP007289 | OBP46 | odorant-binding protein 46 | 59.83945933 | 3.065679524 | 0.296011218 | 10.35663291 | 3.90466E-25 | 3.64939E-24 | |
| AGAP007287 | OBP47 | odorant-binding protein 47 | 16.24857223 | 3.065551738 | 0.551290808 | 5.560679941 | 2.68726E-08 | 8.74193E-08 | |
| AGAP009184 | AGAP009184 | nan | 731.5342998 | 3.063042997 | 0.272885531 | 11.2246442 | 3.08626E-29 | 3.47795E-28 | |
| AGAP011389 | AGAP011389 | nan | 31.10288838 | 3.052289506 | 0.396559049 | 7.696935705 | 1.39368E-14 | 7.46015E-14 | |
| AGAP005713 | AGAP005713 | nan | 15.10544587 | 3.051343247 | 0.452592633 | 6.741919826 | 1.56307E-11 | 6.76733E-11 | |
| AGAP003249 | CLIPB3 | CLIP-domain serine protease | 699.1854621 | 3.050823047 | 0.238996434 | 12.76514047 | 2.56687E-37 | 3.93211E-36 | |
| AGAP007207 | AGAP007207 | nan | 76.68301283 | 3.049419766 | 0.257618134 | 11.83697637 | 2.51352E-32 | 3.22614E-31 | |
| AGAP028615 | AGAP028615 | nan | 129.3585815 | 3.04496956 | 0.293550926 | 10.37288351 | 3.29431E-25 | 3.082E-24 | |
| AGAP004707 | para | voltage-gated sodium channel | 55.99610076 | 3.042386927 | 0.302148911 | 10.06916398 | 7.56181E-24 | 6.63313E-23 | |
| AGAP013034 | IAP8 | inhibitor of apoptosis 8 | 19.73609989 | 3.020611559 | 0.390511598 | 7.735011126 | 1.03394E-14 | 5.62399E-14 | |
| AGAP003186 | AGAP003186 | upstream stimulatory factor | 79.93399687 | 3.004181712 | 0.264086101 | 11.37576608 | 5.52159E-30 | 6.48615E-29 | |
| AGAP003495 | AGAP003495 | nan | 7.878124946 | 2.999308208 | 0.664596658 | 4.512975156 | 6.39245E-06 | 1.63158E-05 | |
| AGAP006215 | GPRMTH1 | methuselah receptor 1 | 282.8197377 | 2.980383514 | 0.199087966 | 14.97018413 | 1.15002E-50 | 2.71538E-49 | |
| AGAP010571 | AGAP010571 | nan | 102.0622474 | 2.972367263 | 0.209646622 | 14.17798785 | 1.25391E-45 | 2.53499E-44 | |
| AGAP003656 | AGAP003656 | Terribly reduced optic lobes, isoform B [Source:UniProtKB/TrEMBL;A | 33382.07926 | 2.969547437 | 0.233294818 | 12.72873295 | 4.09429E-37 | 6.22135E-36 | |
| AGAP029166 | AGAP029166 | nan | 14302.24015 | 2.953223582 | 0.236675158 | 12.4779618 | 9.84752E-36 | 1.43169E-34 | |
| AGAP009576 | AGAP009576 | collagen alpha 1 | 5.519976201 | 2.949676701 | 0.807847551 | 3.651278879 | 0.000260938 | 0.000545321 | |
| AGAP001169 | Gr49 | gustatory receptor 49 | 34.47423754 | 2.94746034 | 0.330978616 | 8.905289344 | 5.3247E-19 | 3.68311E-18 | |
| AGAP005729 | AGAP005729 | cyclin-dependent kinase 14 | 722.6162693 | 2.94622387 | 0.136867615 | 21.52608467 | 8.872E-103 | 7.9603E-101 | |
| AGAP006690 | AGAP006690 | BTB (POZ) domain containing 9 | 6099.788427 | 2.943786111 | 0.171740485 | 17.14089784 | 7.35068E-66 | 2.64316E-64 | |
| AGAP008366 | TEP2 | thioester-containing protein 2 | 2622.051057 | 2.943343731 | 0.0858671 | 34.27789849 | 1.6754E-257 | 1.5784E-254 | |
| AGAP010272 | IR25a | ionotropic receptor IR25a | 24.4642251 | 2.942145602 | 0.369958492 | 7.952637038 | 1.82583E-15 | 1.03934E-14 | |
| AGAP000786 | AGAP000786 | ZP domain-containing protein [Source:UniProtKB/TrEMBL;Acc:Q7QE | 22.87413715 | 2.929738135 | 0.448887659 | 6.52666224 | 6.72515E-11 | 2.78617E-10 | |
| AGAP002802 | AGAP002802 | neural cell adhesion molecule | 308.3665291 | 2.924663628 | 0.224796015 | 13.01030017 | 1.06921E-38 | 1.7102E-37 | |
| AGAP010002 | AGAP010002 | nan | 7.904902055 | 2.924369934 | 0.632889224 | 4.620666336 | 3.8251E-06 | 1.00351E-05 | |
| AGAP011294 | DEF1 | defensin anti-microbial peptide | 9881.820375 | 2.922360646 | 0.52534384 | 5.562758 | 2.65544E-08 | 8.64739E-08 | AM granulocytes |
| AGAP012991 | AGAP012991 | Fascin [Source:UniProtKB/TrEMBL;Acc:F5HLZ2] | 1219.272325 | 2.912953405 | 0.179343014 | 16.2423578 | 2.5301E-59 | 7.73898E-58 | |
| AGAP003120 | AGAP003120 | nan | 11.14672152 | 2.910909881 | 0.538088499 | 5.409723281 | 6.31222E-08 | 1.98423E-07 | |
| AGAP013430 | AGAP013430 | nan | 97.31632396 | 2.905915283 | 0.296411448 | 9.803654009 | 1.08585E-22 | 9.02894E-22 | |
| AGAP027980 | ABCC10 | ATP-binding cassette transporter (ABC transporter) family C membe | 393.4229794 | 2.903222783 | 0.16422288 | 17.67855233 | 6.13514E-70 | 2.49134E-68 | |
| AGAP009212 | SRPN6 | serine protease inhibitor (serpin) 6 | 4770.187149 | 2.896154957 | 0.163756954 | 17.68569143 | 5.40539E-70 | 2.20451E-68 | |
| AGAP011391 | AGAP011391 | nan | 67.78722763 | 2.890477594 | 0.296254065 | 9.75675252 | 1.72594E-22 | 1.41761E-21 | |
| AGAP011390 | AGAP011390 | nan | 14.99695138 | 2.875637539 | 0.573188903 | 5.0169107 | 5.2509E-07 | 1.50911E-06 | |
| AGAP005194 | AGAP005194 | nan | 10.23566919 | 2.867844868 | 0.570150093 | 5.029982282 | 4.90525E-07 | 1.41365E-06 | |
| AGAP009049 | Cht16 | Chitinase [Source:UniProtKB/TrEMBL;Acc:A0A1S4GZY6] | 18.12612964 | 2.866043358 | 0.581910521 | 4.925230346 | 8.4261E-07 | 2.3668E-06 | |
| AGAP010241 | AGAP010241 | gamma-glutamyltranspeptidase | 43.78029184 | 2.850909362 | 0.310313239 | 9.187198613 | 4.03205E-20 | 2.97229E-19 | |
| AGAP012536 | AGAP012536 | nan | 13.33931205 | 2.837957243 | 0.543014631 | 5.226299774 | 1.72936E-07 | 5.22022E-07 | |
| AGAP000628 | AGAP000628 | nan | 95.15290212 | 2.834890127 | 0.327718106 | 8.650392138 | 5.13228E-18 | 3.37884E-17 | |
| AGAP012945 | AGAP012945 | CASPS4 protein | 109.8700581 | 2.834189324 | 0.222741975 | 12.72409172 | 4.34498E-37 | 6.58104E-36 | |
| AGAP013081 | AGAP013081 | nan | 96.77306728 | 2.833323638 | 0.241521745 | 11.7311327 | 8.82695E-32 | 1.1029E-30 | |
| AGAP005547 | AGAP005547 | alpha-crystallin B chain | 52.28380792 | 2.828605593 | 0.307739465 | 9.191559465 | 3.87187E-20 | 2.86318E-19 | |
| AGAP008368 | TEP14 | thioester-containing protein 14 | 9679.75662 | 2.826859645 | 0.179487146 | 15.74964956 | 6.9063E-56 | 1.93069E-54 | |
| AGAP010861 | nan | nan | 2139.682887 | 2.826063415 | 0.268617822 | 10.52075918 | 6.93128E-26 | 6.65643E-25 | |
| AGAP007917 | ABCC12 | ATP-binding cassette transporter (ABC transporter) family C membe | 7223.949465 | 2.813232634 | 0.147885435 | 19.0230541 | 1.09892E-80 | 6.27449E-79 | |
| AGAP013218 | AGAP013218 | sodium-independent sulfate anion transporter | 1458.174905 | 2.812393112 | 0.137755861 | 20.41577819 | 1.21088E-92 | 8.51319E-91 | |
| AGAP009549 | AGAP009549 | nan | 306.909315 | 2.810618837 | 0.221389088 | 12.69538105 | 6.27239E-37 | 9.43965E-36 | |
| AGAP013221 | AGAP013221 | nan | 4538.405584 | 2.794497608 | 0.279013629 | 10.01563119 | 1.30129E-23 | 1.12266E-22 | |
| AGAP010479 | AGAP010479 | nan | 459.3921529 | 2.794359145 | 0.137435605 | 20.33213409 | 6.68316E-92 | 4.49729E-90 | |
| AGAP004483 | AGAP004483 | Lethal giant larvae | 5028.816532 | 2.775249543 | 0.149955674 | 18.50713265 | 1.80876E-76 | 8.92164E-75 | |
| AGAP010417 | AGAP010417 | tetratricopeptide repeat protein 39B | 2743.259669 | 2.766584334 | 0.159767341 | 17.31633212 | 3.54229E-67 | 1.29852E-65 | |
| AGAP000868 | AGAP000868 | beta-1,4-mannosyltransferase egghead protein | 4835.104603 | 2.751447317 | 0.202379799 | 13.59546422 | 4.26049E-42 | 7.65994E-41 | |
| AGAP010328 | AGAP010328 | nan | 36.40248963 | 2.748746522 | 0.366021245 | 7.509800489 | 5.92175E-14 | 3.01724E-13 | |
| AGAP006917 | Gr55 | gustatory receptor 55 | 63.2237979 | 2.74278623 | 0.343766662 | 7.978627747 | 1.47969E-15 | 8.45884E-15 | |
| AGAP006442 | AGAP006442 | nan | 175.4100797 | 2.742217952 | 0.326780091 | 8.391631029 | 4.79444E-17 | 2.97552E-16 | |
| AGAP009460 | AGAP009460 | c-Jun N-terminal kinase | 258.877279 | 2.732183301 | 0.177903847 | 15.35764034 | 3.14826E-53 | 8.10377E-52 | |
| AGAP006342 | PGRPS3 | peptidoglycan recognition protein (short) | 17.74977137 | 2.729041906 | 0.702853207 | 3.882804945 | 0.000103258 | 0.00022841 | |
| AGAP007685 | AGAP007685 | nan | 1642.6336 | 2.719749988 | 0.128717413 | 21.12961975 | 4.2496E-99 | 3.33625E-97 | |
| AGAP007203 | AGAP007203 | DE-cadherin | 681.3832307 | 2.719236461 | 0.148617641 | 18.29686195 | 8.76545E-75 | 4.048E-73 | |
| AGAP013360 | AGAP013360 | nan | 57.65098178 | 2.705703525 | 0.43471435 | 6.224095252 | 4.84343E-10 | 1.87855E-09 | |
| AGAP005162 | AGAP005162 | nan | 2522.373747 | 2.702333581 | 0.191957742 | 14.07775251 | 5.20386E-45 | 1.02779E-43 | |
| AGAP009984 | AGAP009984 | cell cycle control protein 50A | 722.5773174 | 2.691292746 | 0.1492946 | 18.02672537 | 1.20203E-72 | 5.24275E-71 | |
| AGAP001676 | AGAP001676 | actin, cytoplasmic | 14.05968558 | 2.67556056 | 0.684284209 | 3.910013596 | 9.22909E-05 | 0.000205452 | |
| AGAP005100 | Vps60 | vacuolar protein sorting 60 | 4178.014177 | 2.667988174 | 0.082652298 | 32.27966105 | 1.3498E-228 | 6.6929E-226 | |
| AGAP007080 | AGAP007080 | hexosaminidase | 1438.181274 | 2.651861768 | 0.202725883 | 13.08102214 | 4.22717E-39 | 6.8781E-38 | |
| AGAP004281 | AGAP004281 | nan | 17.30103525 | 2.65113093 | 0.48407829 | 5.476657365 | 4.33435E-08 | 1.38608E-07 | |
| AGAP005408 | AGAP005408 | nan | 14.93859068 | 2.643982323 | 0.425276781 | 6.217086002 | 5.06472E-10 | 1.95954E-09 | |
| AGAP013066 | AGAP013066 | nan | 12.67584015 | 2.629542872 | 0.493623819 | 5.327017808 | 9.98384E-08 | 3.06977E-07 | |
| AGAP005246 | SRPN10 | serine protease inhibitor (serpin) 10 | 11070.86802 | 2.628637134 | 0.086237398 | 30.481406 | 4.5966E-204 | 1.6039E-201 | Megacytes |
| AGAP000784 | AGAP000784 | abhydrolase domain containing 4 | 3941.958531 | 2.627781415 | 0.086688661 | 30.31286183 | 7.7597E-202 | 2.6109E-199 | |
| AGAP000307 | AGAP000307 | Integrin alpha-PS1 | 2265.810929 | 2.624431837 | 0.105589431 | 24.85506173 | 2.2797E-136 | 3.3558E-134 | |
| AGAP007036 | APL1A | Anopheles Plasmodium-responsive Leucine-Rich Repeat 1A | 1456.477362 | 2.623653516 | 0.264571135 | 9.916627985 | 3.52463E-23 | 2.9888E-22 | |
| AGAP006427 | ABCC7 | ATP-binding cassette transporter (ABC transporter) family C membe | 5329.59177 | 2.623129975 | 0.199548049 | 13.14535516 | 1.80954E-39 | 2.96481E-38 | |
| AGAP008940 | AGAP008940 | solute carrier family 6 (neurotransmitter transporter, GABA) membe | 506.7084941 | 2.622706075 | 0.200646081 | 13.07130473 | 4.80345E-39 | 7.77548E-38 | |
| AGAP012916 | AGAP012916 | nan | 9061.550348 | 2.615506007 | 0.181322741 | 14.42458894 | 3.62423E-47 | 7.70741E-46 | Granulocytes and Megacytes |
| AGAP013027 | AGAP013027 | Protein toll | 9828.824707 | 2.611676022 | 0.210977531 | 12.3789297 | 3.39845E-35 | 4.84369E-34 | Megacytes |
| AGAP005527 | AGAP005527 | PBPe domain-containing protein [Source:UniProtKB/TrEMBL;Acc:A0A | 28.86170522 | 2.601288169 | 0.399993443 | 6.503327037 | 7.85627E-11 | 3.23488E-10 | |
| AGAP007156 | AGAP007156 | nan | 7.99232791 | 2.59173468 | 0.624807418 | 4.148053635 | 3.35314E-05 | 7.84648E-05 | |
| AGAP001623 | AGAP001623 | nan | 1089.6557 | 2.586344477 | 0.15900407 | 16.26590109 | 1.72313E-59 | 5.28783E-58 | |
| AGAP001647 | AGAP001647 | nan | 13.90756209 | 2.584874496 | 0.700825196 | 3.688329859 | 0.000225731 | 0.000475965 | |
| AGAP011479 | AGAP011479 | hyaluronoglucosaminidase | 59.91269357 | 2.583249954 | 0.256239988 | 10.08136934 | 6.67888E-24 | 5.87504E-23 | |
| AGAP013496 | Cht5-3 | chitinase | 44.57409181 | 2.580734555 | 0.339742141 | 7.596156741 | 3.05055E-14 | 1.58956E-13 | |
| AGAP011653 | AGAP011653 | membrane dipeptidase | 16.29744916 | 2.574841815 | 0.594793929 | 4.328964516 | 1.49812E-05 | 3.6498E-05 | |
| AGAP003247 | CLIPB19 | CLIP-domain serine protease | 1759.954841 | 2.570395939 | 0.274299017 | 9.370780717 | 7.1998E-21 | 5.51458E-20 | |
| AGAP028842 | SSU_rRNA_eukar | Eukaryotic small subunit ribosomal RNA [Source:RFAM;Acc:RF01960 | 90.2435724 | 2.55653131 | 0.711067906 | 3.595340597 | 0.000323967 | 0.000666834 | |
| AGAP013073 | AGAP013073 | nan | 24.81522149 | 2.55620036 | 0.336846785 | 7.588614392 | 3.23344E-14 | 1.68114E-13 | |
| AGAP000163 | GSTMS2 | glutathione transferase microsomal 2 | 163.6791333 | 2.547582272 | 0.223372999 | 11.40505916 | 3.94509E-30 | 4.66332E-29 | |
| AGAP003352 | AGAP003352 | Stomatin (EPB72)-like 3 | 5258.622442 | 2.546156399 | 0.186343329 | 13.66379149 | 1.67054E-42 | 3.05003E-41 | Megacytes |
| AGAP007291 | IAP4 | inhibitor of apoptosis 4 | 154.2438589 | 2.541840917 | 0.260407564 | 9.761010314 | 1.65498E-22 | 1.3629E-21 | |
| AGAP002326 | AGAP002326 | nan | 10.01928458 | 2.541710628 | 0.605077704 | 4.200635074 | 2.66167E-05 | 6.29724E-05 | |
| AGAP004784 | AGAP004784 | fatty acyl-CoA reductase 2 | 1301.202247 | 2.537466672 | 0.135168247 | 18.77265355 | 1.26411E-78 | 6.69054E-77 | |
| AGAP012696 | AGAP012696 | Sulfotransferase (Sult) | 300.8656569 | 2.523311391 | 0.235357097 | 10.72120377 | 8.09429E-27 | 8.16449E-26 | |
| AGAP003790 | ANXB9 | annexin B9 | 7555.362031 | 2.513237969 | 0.223011904 | 11.26952386 | 1.85571E-29 | 2.11143E-28 | |

| | | | | | | | | | |
|---|---|---|---|---|---|---|---|---|---|
| AGAP012384 | AGAP012384 | nan | 14.70484317 | 2.510804297 | 0.40542433 | 6.193028178 | 5.90192E-10 | 2.26669E-09 | |
| AGAP000651 | Actin5C | Actin-5C [Source:UniProtKB/Swiss-Prot;Acc:P84185] | 194466.2946 | 2.509401475 | 0.070345517 | 35.6725146 | 1.055E-278 | 1.66E-275 | Granulocytes and Megacytes |
| AGAP002270 | CLIPB7 | CLIP-domain serine protease | 659.568654 | 2.507618185 | 0.252318085 | 9.938321234 | 2.83566E-23 | 2.41543E-22 | |
| AGAP007821 | AGAP007821 | nan | 2115.812727 | 2.502605815 | 0.347010852 | 7.211894963 | 5.51785E-13 | 2.65765E-12 | |
| AGAP009745 | AGAP009745 | Sugar transporter ERD6-like 4 | 1788.51531 | 2.501737477 | 0.183735738 | 13.61595466 | 3.21912E-42 | 5.83217E-41 | |
| AGAP006103 | AGAP006103 | nan | 1120.036332 | 2.497956821 | 0.229281758 | 10.89470372 | 1.22164E-27 | 1.27312E-26 | |
| AGAP012677 | AGAP012677 | nan | 683.9754154 | 2.49692911 | 0.239766617 | 10.41399817 | 2.14039E-25 | 2.02456E-24 | |
| AGAP006347 | AGAP006347 | potassium voltage-gated channel KQT-like subfamily, invertebrate | 1199.507309 | 2.494910391 | 0.18347811 | 13.59786399 | 4.123E-42 | 7.42691E-41 | |
| AGAP009581 | AGAP009581 | collagen type I/II/III/V/XI/XXIV/XXVII alpha | 15.296804 | 2.487902201 | 0.442494923 | 5.62244236 | 1.88276E-08 | 6.23463E-08 | |
| AGAP007346 | LYSC5 | C-type lysozyme | 26.59299175 | 2.48381085 | 0.358168297 | 6.934759072 | 4.06914E-12 | 1.84216E-11 | |
| AGAP002262 | AGAP002262 | Adenylate cyclase 8 [Source:UniProtKB/TrEMBL;Acc:A0A1S4GF11] | 517.9406016 | 2.482252938 | 0.239223687 | 10.37628409 | 3.17909E-25 | 2.98309E-24 | |
| AGAP001021 | AGAP001021 | dihydropyrimidine dehydrogenase (NADP+) | 6635.508297 | 2.481232948 | 0.185252959 | 13.3937561 | 6.57687E-41 | 1.13689E-39 | |
| AGAP005818 | AGAP005818 | nurim homolog | 195.8631516 | 2.480852224 | 0.165012198 | 15.03435656 | 4.37277E-51 | 1.04558E-49 | |
| AGAP008133 | AGAP008133 | Retinaldehyde binding protein 1 [Source:UniProtKB/TrEMBL;Acc:A0/ | 11.84481105 | 2.480216817 | 0.483138189 | 5.13355573 | 2.84319E-07 | 8.40467E-07 | |
| AGAP004636 | AGAP004636 | sodium-independent sulfate anion transporter | 404.5886291 | 2.479725353 | 0.134483713 | 18.43885254 | 6.40869E-76 | 3.0493E-74 | |
| AGAP000998 | AGAP000998 | insulin-like growth factor 2 receptor | 366.6429897 | 2.475052802 | 0.142281523 | 17.39546185 | 8.9304E-68 | 3.35192E-66 | |
| AGAP010392 | AGAP010392 | calumenin | 17468.3635 | 2.466793358 | 0.161961667 | 15.23072349 | 2.21131E-52 | 5.58519E-51 | |
| AGAP010175 | AGAP010175 | adenylyl cyclase-associated protein 1 | 10435.9944 | 2.460454195 | 0.092042101 | 26.73183437 | 2.0084E-157 | 3.71E-155 | Megacytes |
| AGAP006732 | nan | nan | 7.476875059 | 2.45410562 | 0.695502323 | 3.528536914 | 0.000417864 | 0.000846237 | |
| AGAP011276 | AGAP011276 | nan | 103.0668338 | 2.445281652 | 0.229280843 | 10.66500637 | 1.48389E-26 | 1.4731E-25 | |
| AGAP007035 | APL1B | Anopheles Plasmodium-responsive Leucine-Rich Repeat 1B | 1019.493622 | 2.443761697 | 0.276147234 | 8.849488238 | 8.79215E-19 | 5.98057E-18 | |
| AGAP008219 | CYP6Z1 | cytochrome P450 | 96.79059141 | 2.441510767 | 0.318381888 | 7.668497666 | 1.74023E-14 | 9.25207E-14 | |
| AGAP000889 | AGAP000889 | Coactosin-like protein | 14892.6759 | 2.434451398 | 0.081819665 | 29.75386672 | 1.5455E-194 | 4.6968E-192 | |
| AGAP013184 | CLIPB36 | CLIP-domain serine protease | 1066.00722 | 2.426717555 | 0.273967982 | 8.857668471 | 8.17053E-19 | 5.57787E-18 | |
| AGAP009991 | AGAP009991 | Cyclophilin B precursor | 8.154525466 | 2.424712391 | 0.642102818 | 3.776205809 | 0.000159235 | 0.000342893 | |
| AGAP005170 | AGAP005170 | nan | 542.5839387 | 2.42262086 | 0.178555649 | 13.56787576 | 6.20957E-42 | 1.1017E-40 | |
| AGAP010427 | AGAP010427 | serine palmitoyltransferase | 1434.380115 | 2.421769689 | 0.140631482 | 17.22067963 | 1.85794E-66 | 6.70637E-65 | |
| AGAP027986 | AGAP027986 | nan | 5.970193469 | 2.42116634 | 0.663101131 | 3.651277649 | 0.000260939 | 0.000545321 | |
| AGAP010763 | AGAP010763 | nan | 171.5979988 | 2.413536367 | 0.287441286 | 8.396623872 | 4.59499E-17 | 2.8555E-16 | |
| AGAP006455 | AGAP006455 | nan | 16.77499917 | 2.409577183 | 0.400094243 | 6.022524 | 1.71718E-09 | 6.29967E-09 | |
| AGAP007033 | APL1C | Anopheles Plasmodium-responsive Leucine-Rich Repeat 1C | 11735.16759 | 2.407308159 | 0.261412809 | 9.208837799 | 3.29675E-20 | 2.4398E-19 | |
| AGAP010641 | AGAP010641 | nan | 1925.659844 | 2.401682245 | 0.138090315 | 17.39211214 | 9.46799E-68 | 3.5396E-66 | |
| AGAP006891 | AGAP006891 | NADH dehydrogenase (ubiquinone) Fe-S protein 2 | 11.60807987 | 2.399193682 | 0.477958742 | 5.019666909 | 5.17612E-07 | 1.48853E-06 | |
| AGAP010295 | AGAP010295 | nan | 87.26083349 | 2.396442608 | 0.352047721 | 6.807152739 | 9.95492E-12 | 4.3784E-11 | |
| AGAP013492 | AGAP013492 | glucose dehydrogenase (acceptor) | 8.615409134 | 2.394040751 | 0.573053872 | 4.177688813 | 2.94486E-05 | 6.93415E-05 | |
| AGAP009194 | GSTE2 | glutathione S-transferase epsilon class 2 | 1135.33667 | 2.388029203 | 0.115530754 | 20.67007369 | 6.44178E-95 | 4.7045E-93 | Granulocytes |
| AGAP010636 | AGAP010636 | nan | 212.8105952 | 2.38787568 | 0.221887261 | 10.7616619 | 5.22198E-27 | 5.31278E-26 | |
| AGAP004593 | AGAP004593 | EH domain-containing protein 1 | 5308.523884 | 2.387364869 | 0.135960009 | 17.55931679 | 5.04791E-69 | 1.95705E-67 | |
| AGAP002644 | AGAP002644 | phospholipid-translocating ATPase | 1290.995957 | 2.387018549 | 0.156822506 | 15.22114786 | 2.56E-52 | 6.43139E-51 | |
| AGAP006826 | AGAP006826 | integrin alpha-ps | 8461.448649 | 2.384955984 | 0.157677386 | 15.12554234 | 1.09892E-51 | 2.68906E-50 | |
| AGAP008313 | AGAP008313 | nan | 146.6222392 | 2.384647207 | 0.197837156 | 12.05358617 | 1.85692E-33 | 2.48848E-32 | |
| AGAP007286 | Obp48 | odorant-binding protein 48 | 95.08677734 | 2.376598998 | 0.271125081 | 8.765692166 | 1.85633E-18 | 1.2465E-17 | |
| AGAP011666 | AGAP011666 | nan | 49.21896753 | 2.374004061 | 0.312428533 | 7.598550739 | 2.99465E-14 | 1.5613E-13 | |
| AGAP003878 | AGAP003878 | nan | 37837.14551 | 2.367149181 | 0.086603722 | 27.33311137 | 1.7147E-164 | 3.6714E-162 | Oenocytes |
| AGAP002536 | AGAP002536 | nan | 1726.082374 | 2.365798027 | 0.548850053 | 4.31046333 | 1.62913E-05 | 3.94754E-05 | |
| AGAP003796 | AGAP003796 | cyclin-dependent kinases regulatory subunit 1 | 77.15966343 | 2.363815336 | 0.229968024 | 10.278887 | 8.77349E-25 | 7.95525E-24 | |
| AGAP002165 | nan | nan | 9.072117212 | 2.350037526 | 0.530424579 | 4.430483838 | 9.40219E-06 | 2.35392E-05 | |
| AGAP012429 | NMDAR2 | ionotropic receptor NMDAR2 | 24.05147051 | 2.348667404 | 0.379972206 | 6.181155802 | 6.3634E-10 | 2.43499E-09 | |
| AGAP001678 | nan | nan | 1197.097733 | 2.343244202 | 0.21876871 | 10.71105738 | 9.03242E-27 | 9.07719E-26 | |
| AGAP002877 | AGAP002877 | Tetratricopeptide repeat protein 30 homolog [Source:UniProtKB/Sw | 18.09844722 | 2.340862278 | 0.361282961 | 6.479304403 | 9.21464E-11 | 3.77604E-10 | |
| AGAP008835 | CLIPC1 | CLIP-domain serine protease | 35.05362748 | 2.338817255 | 0.339891883 | 6.881062391 | 5.94078E-12 | 2.65882E-11 | |
| AGAP006745 | AGAP006745 | nan | 4198.803551 | 2.338674916 | 0.15043346 | 15.54624159 | 1.68721E-54 | 4.54148E-53 | Megacytes |
| AGAP009098 | AGAP009098 | Protein-glutamine gamma-glutamyltransferase E | 6078.06949 | 2.330846338 | 0.139367798 | 16.72442542 | 8.70113E-63 | 2.8662E-61 | Megacytes |
| AGAP011064 | AGAP011064 | nan | 326.4754085 | 2.30196808 | 0.148222163 | 15.5305255 | 2.15606E-54 | 5.72175E-53 | |
| AGAP011530 | AGAP011530 | nan | 28.33839906 | 2.296811726 | 0.326839474 | 7.027338835 | 2.1051E-12 | 9.7264E-12 | |
| AGAP010160 | AGAP010160 | myosin I | 7761.456702 | 2.295088071 | 0.087350529 | 26.27446115 | 3.7563E-152 | 6.4343E-150 | |
| AGAP010260 | AGAP010260 | nan | 14.24210831 | 2.294109402 | 0.39661208 | 5.784265079 | 7.283E-09 | 2.50687E-08 | |
| AGAP005520 | AGAP005520 | cytochrome b-561 | 172.0419043 | 2.281209432 | 0.203018804 | 11.23644404 | 2.70042E-29 | 3.0541E-28 | |
| AGAP010774 | AGAP010774 | nan | 101.6516555 | 2.276634432 | 0.294695567 | 7.725377251 | 1.11523E-14 | 6.03824E-14 | |
| AGAP007651 | AGAP007651 | growth arrest and DNA-damage-inducible protein | 473.1325499 | 2.264883499 | 0.227178005 | 9.969642523 | 2.06972E-23 | 1.76941E-22 | |
| AGAP002811 | CLIPD4 | CLIP-domain serine protease [Source:UniProtKB/TrEMBL;Acc:A0A1S | 16.51541341 | 2.263256704 | 0.377785221 | 5.990855598 | 2.0874E-09 | 7.61634E-09 | |
| AGAP001761 | AGAP001761 | ceramide synthetase | 1899.92071 | 2.259442539 | 0.172869761 | 13.07020107 | 4.87365E-39 | 7.87559E-38 | |
| AGAP012802 | AGAP012802 | nan | 117.2076668 | 2.250775294 | 0.312326253 | 7.206487686 | 5.74135E-13 | 2.75544E-12 | |
| AGAP000812 | AGAP000812 | calcium binding protein | 885.5706431 | 2.24680094 | 0.098873996 | 22.72388125 | 2.6017E-114 | 2.8836E-112 | |
| AGAP010056 | AGAP010056 | hexosaminidase | 1842.521798 | 2.246490786 | 0.236944089 | 9.481100788 | 2.51615E-21 | 1.98033E-20 | |
| AGAP009154 | ARC-P34 | actin-related protein 2/3 complex subunit 2 | 7240.757708 | 2.243759331 | 0.116223402 | 19.30557272 | 4.82178E-83 | 2.87506E-81 | |
| AGAP028157 | AGAP028157 | nan | 4114.355096 | 2.241755919 | 0.143207911 | 15.65385536 | 3.12689E-55 | 8.56349E-54 | Granulocytes, Dividing granulocytes and Megacytes |
| AGAP006270 | AGAP006270 | fyn-related kinase | 1025.707773 | 2.234811234 | 0.154332825 | 14.48046604 | 1.61002E-47 | 3.48689E-46 | |
| AGAP007887 | AGAP007887 | nan | 42.77086196 | 2.231869475 | 0.254236509 | 8.778713509 | 1.65355E-18 | 1.11431E-17 | |
| AGAP000204 | AGAP000204 | nan | 23.88038773 | 2.230134227 | 0.459672195 | 4.851575216 | 1.22485E-06 | 3.38197E-06 | |
| AGAP004372 | AGAP004372 | elongation of very long chain fatty acids protein 1 | 1966.839537 | 2.227772939 | 0.125738478 | 17.71751155 | 3.07204E-70 | 1.26937E-68 | |
| AGAP004737 | AGAP004737 | Rhomboid-4, isoform B | 1049.272881 | 2.221631193 | 0.120849602 | 18.38343815 | 1.78301E-75 | 8.39888E-74 | |
| AGAP003716 | twf | Twinfilin [Source:UniProtKB/Swiss-Prot;Acc:Q7QG28] | 167.7877541 | 2.21446621 | 0.154420246 | 14.34051731 | 1.22142E-46 | 2.54579E-45 | |
| AGAP011790 | CLIPA2 | CLIP-domain serine protease | 3495.717731 | 2.214259384 | 0.266506231 | 8.308471353 | 9.69428E-17 | 5.90749E-16 | |
| AGAP001424 | AGAP001424 | heat shock protein 90kDa beta | 12691.79985 | 2.210098739 | 0.133106634 | 16.60397142 | 6.52287E-62 | 2.09734E-60 | |
| AGAP008052 | SAP2 | sensory appendage protein 2 | 73.9829377 | 2.205111504 | 0.504938289 | 4.367091094 | 1.25912E-05 | 3.09475E-05 | |
| AGAP004853 | AGAP004853 | Ral GEF with PH domain and SH3-binding motif 1 | 3043.468304 | 2.202453717 | 0.078606414 | 28.01875307 | 9.6025E-173 | 2.3807E-170 | |
| AGAP010862 | nan | nan | 1779.552611 | 2.200882081 | 0.25857126 | 8.511704201 | 1.71395E-17 | 1.09695E-16 | |
| AGAP028566 | AGAP028566 | nan | 59.29647005 | 2.197113336 | 0.376133612 | 5.841310824 | 5.17917E-09 | 1.81184E-08 | |
| AGAP004163 | GSTD7 | glutathione S-transferase delta class 7 | 1846.751146 | 2.193339829 | 0.104345344 | 21.02000675 | 4.30372E-98 | 3.35086E-96 | |
| AGAP008176 | AGAP008176 | dipeptidyl-peptidase 4 | 7.106160326 | 2.19163709 | 0.577323251 | 3.79620444 | 0.000146928 | 0.000317553 | |
| AGAP008016 | AGAP008016 | acyl-CoA oxidase | 578.9547419 | 2.18355879 | 0.163172805 | 13.38187932 | 7.71711E-41 | 1.33156E-39 | |
| AGAP007347 | LYSC1 | C-type lysozyme | 5361.98254 | 2.181613896 | 0.242153177 | 9.009230936 | 2.07507E-19 | 1.46986E-18 | AM granulocytes |
| AGAP011282 | AGAP011282 | Fatty acid 2-hydroxylase [Source:UniProtKB/TrEMBL;Acc:A0A1S4H68 | 106.8653016 | 2.176193005 | 0.22757132 | 9.562685688 | 1.1474E-21 | 9.16848E-21 | |
| AGAP001022 | AGAP001022 | nan | 17.66041077 | 2.169505972 | 0.355300312 | 6.106118967 | 1.02083E-09 | 3.83462E-09 | |
| AGAP004717 | AGAP004717 | nan | 13.75067012 | 2.16622655 | 0.481022941 | 4.50337471 | 6.68828E-06 | 1.70436E-05 | |
| AGAP010530 | CLIPE4 | CLIP-domain serine protease [Source:UniProtKB/TrEMBL;Acc:A0A1S | 5775.541263 | 2.162472443 | 0.182785362 | 11.83066532 | 2.70985E-32 | 3.4734E-31 | |
| AGAP005749 | GSTO1 | glutathione S-transferase omega class 1 | 4862.243234 | 2.161826025 | 0.063184882 | 34.21429221 | 1.4823E-256 | 1.1637E-253 | Dividing granulocytes |
| AGAP028448 | AGAP028448 | nan | 16.04905571 | 2.160657777 | 0.370407885 | 5.833185157 | 5.43792E-09 | 1.89954E-08 | |
| AGAP013358 | AGAP013358 | nan | 7.400413525 | 2.157689653 | 0.540810451 | 3.989733644 | 6.61475E-05 | 0.000149946 | |
| AGAP008407 | TEP13 | thioester-containing protein 13 | 83.36535265 | 2.151160929 | 0.20890573 | 10.29728065 | 7.24813E-25 | 6.63602E-24 | |
| AGAP010057 | AGAP010057 | nicotinic acetylcholine receptor, beta-2 subunit | 8.909378628 | 2.148743804 | 0.533606387 | 4.026832989 | 5.65332E-05 | 0.000128932 | |
| AGAP007876 | AGAP007876 | nan | 35.40621495 | 2.142008732 | 0.269750402 | 7.940706365 | 2.01033E-15 | 1.13818E-14 | |
| AGAP013732 | AGAP013732 | nan | 114.3835523 | 2.14196842 | 0.248318866 | 8.625878716 | 6.36024E-18 | 4.15821E-17 | |
| AGAP010772 | AGAP010772 | nan | 74.42844502 | 2.131014491 | 0.251900239 | 8.459755717 | 2.67938E-17 | 1.69185E-16 | |
| AGAP001175 | GPRCAL2 | putative calcitonin receptor 2 | 44.37738034 | 2.130429032 | 0.275808013 | 7.724318851 | 1.12453E-14 | 6.07814E-14 | |
| AGAP008599 | AGAP008599 | nan | 22.04010987 | 2.129767992 | 0.346934853 | 6.13881244 | 8.31407E-10 | 3.15325E-09 | |
| AGAP009091 | AGAP009091 | aromatic-L-amino-acid decarboxylase | 617.4251181 | 2.124272653 | 0.355365929 | 5.977704888 | 2.26303E-09 | 8.21581E-09 | |
| AGAP010281 | GPRGBB1 | Putative GABA-B receptor 1 [Source:UniProtKB/TrEMBL;Acc:A0A1S4 | 36.83674083 | 2.121555947 | 0.258337054 | 8.212356367 | 2.16889E-16 | 1.29734E-15 | |
| AGAP010457 | AGAP010457 | thymidylate synthase | 944.988603 | 2.121360635 | 0.159232595 | 13.32240195 | 1.715E-40 | 2.9217E-39 | |
| AGAP009595 | AGAP009595 | sodium/potassium-transporting ATPase subunit beta | 3805.376478 | 2.117317925 | 0.104448969 | 20.27131467 | 2.30419E-91 | 1.50748E-89 | |
| AGAP000562 | AGAP000562 | nan | 9218.890892 | 2.116685642 | 0.162448238 | 13.02990829 | 8.2707E-39 | 1.3274E-37 | |
| AGAP008149 | AGAP008149 | Protein phosphatase 1B (formerly 2C) [Source:UniProtKB/TrEMBL;A | 801.9505509 | 2.115960491 | 0.160443742 | 13.18817718 | 1.02637E-39 | 1.69639E-38 | |
| AGAP028513 | AGAP028513 | nan | 59.0244424 | 2.114564177 | 0.407679322 | 5.186832056 | 2.13901E-07 | 6.40549E-07 | |
| AGAP004164 | GSTD1-4 | glutathione S-transferase delta class 1 | 9459.91605 | 2.110874418 | 0.146147076 | 14.44349405 | 2.75513E-47 | 5.88574E-46 | Granulocytes |
| AGAP001534 | AGAP001534 | nan | 11.40408659 | 2.110162853 | 0.508809793 | 4.147252827 | 3.36488E-05 | 7.87201E-05 | |
| AGAP005814 | AGAP005814 | ecdysoneless homolog | 734.854626 | 2.099993164 | 0.080076655 | 26.22478643 | 1.3864E-151 | 2.3323E-149 | |
| AGAP004836 | nan | nan | 650.0710152 | 2.098146961 | 0.175847135 | 11.93165279 | 8.09501E-33 | 1.05481E-31 | |
| AGAP005448 | AGAP005448 | B9 domain-containing protein 2 | 99.60549667 | 2.095463832 | 0.290572621 | 7.211497852 | 5.53397E-13 | 2.66405E-12 | |
| AGAP012056 | AGAP012056 | cofilin | 21864.66673 | 2.09271144 | 0.074059359 | 28.25721795 | 1.1605E-175 | 3.0369E-173 | Granulocytes and Megacytes |
| AGAP012317 | AGAP012317 | nan | 27.02636328 | 2.090756579 | 0.294743093 | 7.093487945 | 1.30773E-12 | 6.10513E-12 | |
| AGAP009528 | AGAP009528 | nan | 517.7739376 | 2.08733478 | 0.118753804 | 17.57699296 | 3.69669E-69 | 1.44508E-67 | |
| AGAP009211 | AGAP009211 | nan | 10.53782511 | 2.086759309 | 0.487911215 | 4.276924252 | 1.89493E-05 | 4.56554E-05 | |
| AGAP008245 | AGAP008245 | nan | 44.13685542 | 2.084210574 | 0.251736754 | 8.279325676 | 1.23875E-16 | 7.49053E-16 | |
| AGAP002461 | AGAP002461 | nan | 707.2169162 | 2.083713242 | 0.147056478 | 14.16947606 | 1.41553E-45 | 2.8495E-44 | |
| AGAP013079 | AGAP013079 | nan | 1464.396434 | 2.081746303 | 0.133219553 | 15.62643213 | 4.80992E-55 | 1.30589E-53 | |
| AGAP003838 | AGAP003838 | nan | 8.749472898 | 2.080685019 | 0.533941455 | 3.896841125 | 9.74555E-05 | 0.000216132 | |
| AGAP003248 | AGAP003248 | nan | 1059.751232 | 2.079895508 | 0.269283056 | 7.723826152 | 1.12889E-14 | 6.0982E-14 | |
| AGAP008551 | AGAP008551 | mothers against decapentaplegic homolog 1 | 487.1543997 | 2.066465676 | 0.121255416 | 17.04225469 | 3.99046E-65 | 1.38724E-63 | |
| AGAP004397 | AGAP004397 | alpha-1,6-mannosyl-glycoprotein beta-1,2-N-acetylglucosaminyltran | 573.8103307 | 2.064323208 | 0.10145681 | 20.34681765 | 4.95411E-92 | 3.35775E-90 | |
| AGAP028537 | AGAP028537 | nan | 11.17762809 | 2.054944515 | 0.454772595 | 4.51861994 | 6.2244E-06 | 1.59089E-05 | |
| AGAP010026 | AGAP010026 | phospholipid-translocating ATPase | 1565.02341 | 2.0488596 | 0.194429848 | 10.53778326 | 5.78463E-26 | 5.58944E-25 | |
| AGAP009221 | SRPN5 | serine protease inhibitor (serpin) 5 | 2214.441769 | 2.047353148 | 0.174240138 | 11.75018094 | 7.04678E-32 | 8.83991E-31 | |
| AGAP003758 | AGAP003758 | nan | 270.257497 | 2.041893372 | 0.194482168 | 10.49912898 | 8.71808E-26 | 8.34686E-25 | |
| AGAP003689 | CLIPC7 | CLIP-domain serine protease [Source:UniProtKB/TrEMBL;Acc:A0A1S | 1211.441982 | 2.04098826 | 0.356864366 | 5.719226844 | 1.0701E-08 | 3.62771E-08 | |
| AGAP011938 | AGAP011938 | lamin | 3921.036628 | 2.033845313 | 0.120428121 | 16.88845846 | 5.4714E-64 | 1.85417E-62 | |
| AGAP000748 | AGAP000748 | Muscular protein 20 | 9.564759859 | 2.033725044 | 0.465122407 | 4.372451238 | 1.22859E-05 | 3.02208E-05 | |
| AGAP004958 | AGAP004958 | ethanolaminephosphotransferase | 294.4964352 | 2.033487138 | 0.13376219 | 15.20225657 | 3.41646E-52 | 8.47013E-51 | |
| AGAP008670 | AGAP008670 | nan | 1126.615038 | 2.031506182 | 0.150608881 | 13.48862147 | 1.82481E-41 | 3.19545E-40 | |
| AGAP011562 | Oseg4 | outer segment 4 | 87.84653089 | 2.016660354 | 0.214531796 | 9.40028654 | 5.44147E-21 | 4.18482E-20 | |
| AGAP012386 | AGAP012386 | nan | 68937.7028 | 2.006246014 | 0.190819364 | 10.51384918 | 7.45858E-26 | 7.15553E-25 | |
| AGAP009828 | AGAP009828 | nan | 15.00615373 | 2.004345543 | 0.575368838 | 3.483583768 | 0.000494748 | 0.000992552 | |
| AGAP001886 | AGAP001886 | nan | 2720.981628 | 2.004081796 | 0.117050849 | 17.12146308 | 1.02662E-65 | 3.67025E-64 | |
