## Supplemental table S2 for "Hemocyte Differentiation to the Megacyte Lineage Enhances Mosquito Immunity Against *Plasmodium*"

| Downregulated genes by Cactus Knockdown | | | | | | | | | |
|---|---|---|---|---|---|---|---|---|---|
| **Accesion Number** | **Name** | **Description** | **baseMean** | **log2FoldChange** | **lfcSE** | **stat** | **pvalue** | **padj** | **Hemocyte type** |
| AGAP004978 | PPO9 | prophenoloxidase 9 | 23777.8002 | -7.931905616 | 0.1685 | -47.063 | 0 | 0 | Oenocytoids |
| AGAP028406 | AGAP028406 | nan | 7750.25117 | -7.799283944 | 0.225 | -34.657 | 3.5436E-263 | 4.173E-260 | Oenocytoids |
| AGAP002867 | CYP6P4 | cytochrome P450 | 25.5488877 | -7.715952754 | 0.9071 | -8.5057 | 1.80456E-17 | 1.15182E-16 | |
| AGAP000144 | AGAP000144 | progestin and adipoQ receptor family member 3 | 499.855567 | -7.69138119 | 0.3774 | -20.382 | 2.43118E-92 | 1.68413E-90 | |
| AGAP006000 | CPR25 | cuticular protein RR-1 family 25 | 15.7863562 | -7.610463392 | 0.9184 | -8.2864 | 1.16761E-16 | 7.07401E-16 | |
| AGAP003581 | AGAP003581 | D-xylulose reductase A | 1106.31964 | -7.122975868 | 0.2636 | -27.026 | 7.223E-161 | 1.4793E-158 | |
| AGAP007990 | AGAP007990 | glucosyl/glucuronosyl transferases | 52.8642935 | -6.975944305 | 1.2596 | -5.5381 | 3.05766E-08 | 9.89565E-08 | |
| AGAP008880 | AGAP008880 | nan | 35.973525 | -6.738191478 | 0.7037 | -9.5758 | 1.01035E-21 | 8.08712E-21 | |
| AGAP006136 | AGAP006136 | monocyte to macrophage differentiation factor 2 | 330.163699 | -6.442614237 | 0.3604 | -17.877 | 1.76868E-71 | 7.4387E-70 | |
| AGAP003571 | AGAP003571 | threonine dehydratase | 1317.19158 | -6.372713602 | 0.1958 | -32.555 | 1.8004E-232 | 9.9772E-230 | |
| AGAP000210 | AGAP000210 | triacylglycerol lipase | 1077.16794 | -6.324830509 | 0.2276 | -27.788 | 6.0342E-170 | 1.4212E-167 | |
| AGAP005752 | AGAP005752 | glucosyl/glucuronosyl transferases | 2189.21022 | -6.213122667 | 0.2718 | -22.859 | 1.1956E-115 | 1.408E-113 | |
| AGAP006327 | LRIM6 | leucine-rich immune protein (Short) | 682.280858 | -6.197194105 | 0.3656 | -16.952 | 1.87318E-64 | 6.39394E-63 | |
| AGAP005751 | AGAP005751 | glucosyl/glucuronosyl transferases | 42.4611745 | -6.194983424 | 0.5971 | -10.375 | 3.21893E-25 | 3.01746E-24 | |
| AGAP006571 | AGAP006571 | nuclear receptor subfamily 1 group D member 3 | 9.68565768 | -6.143983933 | 0.9094 | -6.7559 | 1.41917E-11 | 6.16696E-11 | |
| AGAP002588 | AGAP002588 | nan | 8419.60502 | -6.108430513 | 0.182 | -33.563 | 5.9001E-247 | 3.7056E-244 | |
| AGAP008387 | AGAP008387 | nan | 150.384514 | -6.072258913 | 0.6293 | -9.6498 | 4.92527E-22 | 3.9932E-21 | |
| AGAP012251 | AGAP012251 | inorganic phosphate cotransporter | 4805.26247 | -6.026570224 | 0.2373 | -25.397 | 2.7104E-142 | 4.3279E-140 | |
| AGAP006137 | nan | nan | 22.8320718 | -5.976841918 | 0.697 | -8.575 | 9.90601E-18 | 6.40965E-17 | |
| AGAP009382 | AGAP009382 | nan | 4.77124769 | -5.785400182 | 1.0578 | -5.4691 | 4.52336E-08 | 1.44456E-07 | |
| AGAP004449 | AGAP004449 | pancreatic triacylglycerol lipase | 83.1501711 | -5.775735305 | 0.5781 | -9.9902 | 1.68237E-23 | 1.4435E-22 | |
| AGAP028980 | SSU_rRNA_eukary | Eukaryotic small subunit ribosomal RNA [Source:RFAM;Acc:RF01960 | 655.932095 | -5.712182634 | 0.9746 | -5.8608 | 4.60615E-09 | 1.62041E-08 | |
| AGAP028554 | AGAP028554 | nan | 1196.48986 | -5.646977962 | 0.2079 | -27.167 | 1.5969E-162 | 3.3433E-160 | |
| AGAP002830 | AGAP002830 | C-1-tetrahydrofolate synthase, mitochondrial precursor | 8450.48974 | -5.597586948 | 0.2356 | -23.76 | 8.5705E-125 | 1.0911E-122 | |
| AGAP003636 | AGAP003636 | inositol oxygenase | 2734.91946 | -5.581700591 | 0.2222 | -25.118 | 3.1793E-139 | 4.9102E-137 | |
| AGAP008713 | AGAP008713 | nan | 38.4404323 | -5.575036836 | 0.5162 | -10.799 | 3.46939E-27 | 3.56435E-26 | |
| AGAP010363 | AGAP010363 | nan | 14.7019961 | -5.566885964 | 0.8219 | -6.7732 | 1.25987E-11 | 5.49756E-11 | |
| AGAP009385 | AGAP009385 | cellular retinaldehyde-binding protein | 18.0446469 | -5.565601097 | 0.6956 | -8.0008 | 1.23646E-15 | 7.09424E-15 | |
| AGAP004572 | AGAP004572 | nan | 26.4052676 | -5.542488691 | 0.6784 | -8.1703 | 3.0753E-16 | 1.82906E-15 | |
| AGAP003714 | HPX3 | heme peroxidase 3 | 3408.31985 | -5.478975624 | 0.2971 | -18.44 | 6.24116E-76 | 2.9999E-74 | |
| AGAP012635 | AGAP012635 | nan | 12.9650626 | -5.472974589 | 0.7687 | -7.1197 | 1.08199E-12 | 5.08909E-12 | |
| AGAP000352 | TWDL1 | cuticular protein TWDL family (TWDL1) | 16.2536187 | -5.466467959 | 0.7143 | -7.6528 | 1.96607E-14 | 1.04175E-13 | |
| AGAP003999 | AGAP003999 | nan | 163.64635 | -5.439859124 | 0.3155 | -17.239 | 1.34382E-66 | 4.8693E-65 | |
| AGAP003066 | CYP304B1 | cytochrome P450 | 116.636168 | -5.438909643 | 0.3502 | -15.529 | 2.208E-54 | 5.84314E-53 | |
| AGAP012354 | AGAP012354 | Syntrophin, alpha 1 | 60.4766239 | -5.431457844 | 0.5406 | -10.047 | 9.49269E-24 | 8.25767E-23 | |
| AGAP011098 | AGAP011098 | nan | 137.616454 | -5.404158307 | 0.3614 | -14.954 | 1.47261E-50 | 3.46837E-49 | Oenocytoids |
| AGAP008714 | AGAP008714 | nan | 60.861685 | -5.374765101 | 0.4547 | -11.82 | 3.07677E-32 | 3.92767E-31 | |
| AGAP003582 | AGAP003582 | D-xylulose reductase A | 17.4712129 | -5.29214823 | 0.7554 | -7.0056 | 2.45926E-12 | 1.13128E-11 | |
| AGAP010596 | AGAP010596 | Alkaline phosphatase [Source:UniProtKB/TrEMBL;Acc:Q7Q8R8] | 4.84641083 | -5.291703832 | 1.0968 | -4.8245 | 1.40358E-06 | 3.84842E-06 | |
| AGAP000218 | AGAP000218 | nan | 102.125953 | -5.289965987 | 0.4205 | -12.581 | 2.69796E-36 | 3.95911E-35 | |
| AGAP003545 | Osk | protein oskar | 27.043257 | -5.253269891 | 0.7628 | -6.887 | 5.69752E-12 | 2.55358E-11 | |
| AGAP009219 | AGAP009219 | nan | 23.1633731 | -5.226698387 | 0.6939 | -7.5324 | 4.97972E-14 | 2.55801E-13 | |
| AGAP006135 | AGAP006135 | nan | 3.35151891 | -5.203138233 | 1.0572 | -4.9216 | 8.58324E-07 | 2.40878E-06 | |
| AGAP002317 | Alpha_amylase | amylase | 7282.67516 | -5.083876823 | 0.2684 | -18.941 | 5.22851E-80 | 2.91466E-78 | Oenocytoids |
| AGAP004563 | ANCE9 | angiotensin-converting enzyme 9 | 148.969171 | -5.063188132 | 0.3802 | -13.319 | 1.8048E-40 | 3.06913E-39 | |
| AGAP012617 | AGAP012617 | nan | 179.468881 | -5.043696632 | 0.3593 | -14.039 | 8.96638E-45 | 1.75984E-43 | |
| AGAP005753 | AGAP005753 | glucosyl/glucuronosyl transferases | 141.092797 | -5.024762823 | 0.3538 | -14.203 | 8.76391E-46 | 1.77941E-44 | |
| AGAP007252 | AGAP007252 | Peptidase S1 domain-containing protein [Source:UniProtKB/TrEMBL; | 46.288377 | -5.020262381 | 0.4157 | -12.077 | 1.3917E-33 | 1.87572E-32 | |
| AGAP028560 | AGAP028560 | nan | 474.462774 | -5.002460802 | 0.2269 | -22.049 | 9.7677E-108 | 1.0002E-105 | |
| AGAP005585 | AGAP005585 | nan | 301.522027 | -4.979277064 | 0.2091 | -23.814 | 2.3956E-125 | 3.0917E-123 | |
| AGAP000844 | AGAP000844 | Progestin and adipoQ receptor family member 4 | 788.790571 | -4.907435023 | 0.2318 | -21.17 | 1.8188E-99 | 1.43987E-97 | |
| AGAP011253 | AGAP011253 | nan | 9.54328575 | -4.893404448 | 0.8168 | -5.9911 | 2.08486E-09 | 7.61001E-09 | |
| AGAP005497 | AGAP005497 | SCP domain-containing protein [Source:UniProtKB/TrEMBL;Acc:A0A1 | 3.608338 | -4.856641566 | 1.1163 | -4.3507 | 1.35698E-05 | 3.31968E-05 | |
| AGAP001477 | AGAP001477 | Innexin inx2 | 70.8757624 | -4.795367874 | 0.344 | -13.938 | 3.70678E-44 | 7.17075E-43 | |
| AGAP006609 | AGAP006609 | nan | 21.0348322 | -4.786633919 | 0.6277 | -7.6257 | 2.4273E-14 | 1.27681E-13 | |
| AGAP002506 | AGAP002506 | runt-related transcription factor | 324.201393 | -4.758197781 | 0.1993 | -23.879 | 5.1199E-126 | 6.7936E-124 | |
| AGAP011230 | AGAP011230 | nan | 543.436757 | -4.734809459 | 0.2599 | -18.219 | 3.6357E-74 | 1.64673E-72 | |
| AGAP004450 | AGAP004450 | nan | 134.456281 | -4.713058251 | 0.3123 | -15.091 | 1.85261E-51 | 4.48673E-50 | |
| AGAP006001 | CPR26 | cuticular protein RR-1 family 26 | 127.926692 | -4.681609602 | 0.3444 | -13.593 | 4.39401E-42 | 7.88495E-41 | |
| AGAP004980 | PPO7 | prophenoloxidase 7 | 13.6523774 | -4.67237043 | 0.7342 | -6.364 | 1.96542E-10 | 7.86918E-10 | |
| AGAP003712 | Ast1 | allatostatin 1 | 7.08833786 | -4.665661923 | 0.8559 | -5.4514 | 4.99774E-08 | 1.58852E-07 | |
| AGAP012616 | PPO5 | prophenoloxidase 5 | 21687.4741 | -4.651552574 | 0.2342 | -19.863 | 8.48134E-88 | 5.39883E-86 | Oenocytoids |
| AGAP028480 | AGAP028480 | nan | 3.47488169 | -4.641297013 | 1.0865 | -4.272 | 1.93769E-05 | 4.66165E-05 | |
| AGAP007285 | AGAP007285 | alpha-L-fucosidase | 311.378893 | -4.615829479 | 0.2811 | -16.42 | 1.38179E-60 | 4.33928E-59 | |
| AGAP009924 | AGAP009924 | nan | 9.66472208 | -4.603626508 | 0.7476 | -6.1582 | 7.35802E-10 | 2.80534E-09 | |
| AGAP004981 | PPO4 | prophenoloxidase 4 | 7206.30439 | -4.599210999 | 0.149 | -30.876 | 2.5449E-209 | 9.5902E-207 | Oenocytoids |
| AGAP010975 | AGAP010975 | nan | 26.7616799 | -4.588901081 | 0.7916 | -5.7969 | 6.75457E-09 | 2.33436E-08 | |
| AGAP010657 | AGAP010657 | nan | 41.8936143 | -4.566657813 | 0.4559 | -10.016 | 1.29679E-23 | 1.1198E-22 | |
| AGAP007075 | AGAP007075 | nan | 19.3364744 | -4.564451244 | 0.6107 | -7.4736 | 7.8054E-14 | 3.94077E-13 | |
| AGAP001112 | AGAP001112 | nan | 3.13585904 | -4.540052611 | 1.1756 | -3.862 | 0.000112478 | 0.000247006 | |
| AGAP028398 | AGAP028398 | nan | 8.05143187 | -4.528075965 | 0.8467 | -5.3478 | 8.90251E-08 | 2.75618E-07 | |
| AGAP012644 | AGAP012644 | nan | 5.27808429 | -4.507236925 | 1.0576 | -4.2616 | 2.03004E-05 | 4.86394E-05 | |
| AGAP004154 | AGAP004154 | jagged | 857.959562 | -4.503198656 | 0.2068 | -21.78 | 3.6169E-105 | 3.5129E-103 | |
| AGAP003767 | AGAP003767 | nan | 5196.80068 | -4.496935919 | 0.2348 | -19.151 | 9.52588E-82 | 5.53971E-80 | Oenocytoids |
| AGAP029126 | AGAP029126 | nan | 18.411012 | -4.443084044 | 0.647 | -6.8671 | 6.55376E-12 | 2.92621E-11 | |
| AGAP004249 | AGAP004249 | nan | 51.395445 | -4.437586128 | 0.3893 | -11.399 | 4.20638E-30 | 4.95974E-29 | |
| AGAP006385 | AGAP006385 | nan | 283.540464 | -4.43075434 | 0.7771 | -5.7014 | 1.18834E-08 | 4.00549E-08 | |
| AGAP000605 | AGAP000605 | nan | 261.956624 | -4.424556557 | 0.8846 | -5.0018 | 5.67923E-07 | 1.62577E-06 | |
| AGAP007918 | AGAP007918 | xanthine dehydrogenase/oxidase | 4030.929 | -4.421348327 | 0.1688 | -26.191 | 3.3598E-151 | 5.5532E-149 | |
| AGAP011603 | AGAP011603 | Long-chain acyl-CoA synthetase [Source:UniProtKB/TrEMBL;Acc:A0A | 8573.55885 | -4.420170699 | 0.2497 | -17.703 | 4.00532E-70 | 1.64778E-68 | |
| AGAP004510 | AGAP004510 | Innexin inx2 | 459.008065 | -4.418131468 | 0.2672 | -16.537 | 1.97167E-61 | 6.29665E-60 | |
| AGAP028913 | LSU_rRNA_eukary | Eukaryotic large subunit ribosomal RNA [Source:RFAM;Acc:RF02543] | 99.0881335 | -4.408960933 | 0.9931 | -4.4394 | 9.02088E-06 | 2.26327E-05 | |
| AGAP008141 | AGAP008141 | argininosuccinate lyase | 6002.19367 | -4.39651821 | 0.1751 | -25.108 | 4.0318E-139 | 6.1264E-137 | |
| AGAP007588 | AGAP007588 | glucosyl/glucuronosyl transferases | 394.075897 | -4.386813885 | 0.1809 | -24.256 | 5.7273E-130 | 8.0533E-128 | |
| AGAP000211 | AGAP000211 | triacylglycerol lipase | 5103.25989 | -4.384911177 | 0.1581 | -27.739 | 2.34E-169 | 5.3769E-167 | |
| AGAP028555 | AGAP028555 | nan | 498.282233 | -4.359203037 | 0.2572 | -16.952 | 1.86432E-64 | 6.38683E-63 | |
| AGAP003176 | AGAP003176 | solute carrier family 23 (nucleobase transporter) | 76.3543813 | -4.350232637 | 0.3338 | -13.033 | 7.92785E-39 | 1.27454E-37 | |
| AGAP013159 | nan | nan | 19.4741764 | -4.327824047 | 0.5111 | -8.4678 | 2.49991E-17 | 1.58065E-16 | |
| AGAP007455 | LRIM10 | leucine-rich immune protein (Short) | 3911.49684 | -4.327593865 | 0.2484 | -17.421 | 5.72668E-68 | 2.16671E-66 | |
| AGAP000153 | AGAP000153 | nan | 40.8878693 | -4.315642723 | 0.4851 | -8.8971 | 5.73422E-19 | 3.95766E-18 | |
| AGAP004977 | PPO6 | prophenoloxidase 6 | 79083.991 | -4.312821861 | 0.1806 | -23.881 | 4.8665E-126 | 6.5496E-124 | Oenocytoids |
| AGAP009198 | AGAP009198 | nan | 128.734058 | -4.289383195 | 0.3046 | -14.08 | 5.04817E-45 | 9.99135E-44 | |

| | | | | | | | | | |
|---|---|---|---|---|---|---|---|---|---|
| AGAP013267 | AGAP013267 | nan | 170.044983 | -4.287799532 | 0.3139 | -13.662 | 1.7167E-42 | 3.12824E-41 | |
| AGAP000356 | AGAP000356 | nan | 8.52122871 | -4.252587471 | 0.7446 | -5.7116 | 1.11932E-08 | 3.78095E-08 | |
| AGAP002518 | AGAP002518 | delta-1-pyrroline-5-carboxylate synthetase | 37793.214 | -4.207224571 | 0.1507 | -27.914 | 1.8217E-171 | 4.4007E-169 | Granulocytes |
| AGAP012960 | AGAP012960 | nan | 257.301812 | -4.20622211 | 0.2151 | -19.555 | 3.73425E-85 | 2.32983E-83 | |
| AGAP009039 | AGAP009039 | glucose-6-phosphate 1-epimerase | 151.953295 | -4.20538689 | 0.3275 | -12.84 | 9.8257E-38 | 1.52752E-36 | |
| AGAP009516 | nan | nan | 8.12600818 | -4.204435416 | 0.8637 | -4.8679 | 1.12799E-06 | 3.1246E-06 | |
| AGAP003087 | AGAP003087 | nan | 5.12561631 | -4.202534821 | 0.9682 | -4.3406 | 1.42077E-05 | 3.47212E-05 | |
| AGAP001251 | AGAP001251 | eupolytin | 11.7228522 | -4.184904405 | 0.6219 | -6.7295 | 1.70238E-11 | 7.34345E-11 | |
| AGAP010004 | AGAP010004 | nan | 5.07615339 | -4.182754498 | 0.9041 | -4.6262 | 3.72375E-06 | 9.78562E-06 | |
| AGAP010725 | nan | nan | 174.004261 | -4.178778315 | 0.2638 | -15.838 | 1.70021E-56 | 4.88343E-55 | |
| AGAP003086 | AGAP003086 | nan | 7.02442114 | -4.160037236 | 0.8685 | -4.7898 | 1.6693E-06 | 4.54259E-06 | |
| AGAP006570 | AGAP006570 | myo-inositol-1(or 4)-monophosphatase | 8859.50488 | -4.157395297 | 0.1789 | -23.241 | 1.7485E-119 | 2.1393E-117 | Oenocytoids |
| AGAP006134 | AGAP006134 | nan | 6.65423033 | -4.146921508 | 0.9373 | -4.4243 | 9.6739E-06 | 2.41499E-05 | |
| AGAP002587 | AGAP002587 | nan | 19.6541663 | -4.144652571 | 0.6407 | -6.4688 | 9.87523E-11 | 4.04147E-10 | |
| AGAP005972 | AGAP005972 | 4-nitrophenylphosphatase | 318.701758 | -4.139715515 | 0.2871 | -14.421 | 3.7963E-47 | 8.01904E-46 | |
| AGAP001880 | AGAP001880 | bZIP factor, other | 16.0847949 | -4.130949948 | 0.7097 | -5.8211 | 5.84686E-09 | 2.03409E-08 | |
| AGAP000679 | AGAP000679 | Aminoacylase [Source:UniProtKB/TrEMBL;Acc:Q7QEF5] | 31526.131 | -4.104723825 | 0.1533 | -26.784 | 4.9967E-158 | 9.4147E-156 | Oenocytoids |
| AGAP001010 | AGAP001010 | chondroitin sulfate synthase | 170.09409 | -4.103161998 | 0.2402 | -17.079 | 2.11983E-65 | 7.50785E-64 | |
| AGAP013241 | CYP4D16 | cytochrome P450 | 5.30215358 | -4.090586709 | 0.8342 | -4.9034 | 9.42055E-07 | 2.63122E-06 | |
| AGAP003321 | AGAP003321 | glycine dehydrogenase | 1498.67239 | -4.087626651 | 0.1948 | -20.986 | 8.7899E-98 | 6.78768E-96 | |
| AGAP006258 | PPO2 | prophenoloxidase 2 | 28442.0768 | -4.051077734 | 0.2318 | -17.475 | 2.23633E-68 | 8.5644E-67 | Oenocytoids |
| AGAP009899 | AGAP009899 | nan | 8.83590922 | -4.038931909 | 0.7201 | -5.6089 | 2.03628E-08 | 6.72407E-08 | |
| AGAP002721 | AGAP002721 | Tryptophan 2,3-dioxygenase [Source:UniProtKB/Swiss-Prot;Acc:O774 | 2065.79253 | -4.024997514 | 0.2225 | -18.089 | 3.87998E-73 | 1.73238E-71 | |
| AGAP010799 | AGAP010799 | nan | 5.39154222 | -4.010443217 | 0.988 | -4.0593 | 4.92201E-05 | 0.000113098 | |
| AGAP008359 | AGAP008359 | sodium-coupled monocarboxylate transporter 1 | 3.85256215 | -4.010249167 | 1.0686 | -3.7527 | 0.000174967 | 0.000374628 | |
| AGAP011228 | AGAP011228 | nan | 8413.98229 | -4.009800464 | 0.4668 | -8.5898 | 8.71082E-18 | 5.66745E-17 | Granulocytes |
| AGAP006906 | AGAP006906 | Cat eye syndrome critical region protein 1 | 1347.74014 | -4.000927267 | 0.1127 | -35.501 | 4.6726E-276 | 6.2886E-273 | |
| AGAP006726 | COEAE5G | carboxylesterase | 2201.17039 | -3.997497366 | 0.1943 | -20.57 | 5.05574E-94 | 3.63588E-92 | Oenocytoids |
| AGAP007076 | AGAP007076 | nan | 67.4688145 | -3.987537468 | 0.2863 | -13.93 | 4.15336E-44 | 8.01819E-43 | |
| AGAP009874 | CPR76 | cuticular protein RR-1 family 76 | 5.80861262 | -3.970946909 | 0.9897 | -4.0121 | 6.01712E-05 | 0.000136926 | |
| AGAP005750 | AGAP005750 | glucosyl/glucuronosyl transferases | 1511.87594 | -3.963930585 | 0.1739 | -22.792 | 5.5602E-115 | 6.3882E-113 | |
| AGAP009940 | AGAP009940 | nan | 284.247524 | -3.961724089 | 0.2689 | -14.734 | 3.8722E-49 | 8.68572E-48 | |
| AGAP006206 | nan | nan | 190.510737 | -3.940862111 | 0.2233 | -17.651 | 1.00808E-69 | 4.04134E-68 | |
| AGAP007456 | LRIM8B | leucine-rich immune protein (Short) | 6118.40636 | -3.9304415 | 0.2273 | -17.294 | 5.19666E-67 | 1.89026E-65 | Granulocytes |
| AGAP013029 | AGAP013029 | nan | 140.185467 | -3.929349144 | 0.3094 | -12.698 | 6.04565E-37 | 9.11298E-36 | |
| AGAP001252 | AGAP001252 | eupolytin | 11.917833 | -3.928480813 | 0.5576 | -7.0447 | 1.85817E-12 | 8.61505E-12 | |
| AGAP001962 | GPRVPR1 | GPCR vasopressin family receptor 1 | 171.2186 | -3.907937459 | 0.1723 | -22.679 | 7.2564E-114 | 7.9491E-112 | |
| AGAP006549 | AGAP006549 | nan | 19.9821355 | -3.891416706 | 0.5028 | -7.7395 | 9.98344E-15 | 5.44294E-14 | |
| AGAP011617 | AGAP011617 | nan | 4.90321474 | -3.886640253 | 0.9136 | -4.2541 | 2.09856E-05 | 5.01791E-05 | |
| AGAP006168 | AGAP006168 | BMP binding endothelial regulator | 694.700242 | -3.877349092 | 0.2095 | -18.511 | 1.69422E-76 | 8.40065E-75 | |
| AGAP003067 | CYP304C1 | cytochrome P450 | 233.071707 | -3.875724352 | 0.2526 | -15.343 | 3.92708E-53 | 1.00535E-51 | |
| AGAP009283 | AGAP009283 | nan | 6.56033597 | -3.818322579 | 0.9537 | -4.0035 | 6.24064E-05 | 0.00014167 | |
| AGAP028468 | AGAP028468 | nan | 707.936092 | -3.817097591 | 0.2165 | -17.634 | 1.35679E-69 | 5.37072E-68 | |
| AGAP003289 | AGAP003289 | carbonic anhydrase | 6.26895793 | -3.813869106 | 0.9956 | -3.8308 | 0.000127742 | 0.000278191 | |
| AGAP013755 | AGAP013755 | nan | 7789.42614 | -3.804581874 | 0.246 | -15.468 | 5.74805E-54 | 1.51264E-52 | |
| AGAP000466 | ACE2 | acetylcholinesterase | 130.014914 | -3.801941195 | 0.2838 | -13.395 | 6.48728E-41 | 1.12554E-39 | |
| AGAP007674 | AGAP007674 | GIPC PDZ domain containing family, member 2 | 411.415616 | -3.799060491 | 0.2166 | -17.543 | 6.74851E-69 | 2.60564E-67 | |
| AGAP007454 | LRIM8A | leucine-rich immune protein (Short) | 9438.76355 | -3.794200054 | 0.2357 | -16.095 | 2.75087E-58 | 8.33309E-57 | |
| AGAP010604 | ILP6 | Insulin-like peptide 6 | 8.52382176 | -3.792948395 | 0.8673 | -4.3732 | 1.22421E-05 | 3.01288E-05 | |
| AGAP001415 | AGAP001415 | nan | 3.27470474 | -3.790409077 | 1.0779 | -3.5166 | 0.000437106 | 0.000883306 | |
| AGAP009521 | AGAP009521 | nan | 97.5543461 | -3.789954676 | 0.3334 | -11.368 | 6.06949E-30 | 7.08559E-29 | |
| AGAP011972 | AGAP011972 | Epoxide hydrolase | 198.134239 | -3.788799702 | 0.1764 | -21.48 | 2.3734E-102 | 2.0897E-100 | |
| AGAP000275 | AGAP000275 | epidermal retinal dehydrogenase | 612.865245 | -3.774113828 | 0.1771 | -21.313 | 8.688E-101 | 7.4409E-99 | |
| AGAP001476 | AGAP001476 | Innexin inx2 | 1293.10692 | -3.773854109 | 0.2375 | -15.89 | 7.46388E-57 | 2.17029E-55 | |
| AGAP001281 | AGAP001281 | potassium inwardly-rectifying channel subfamily J | 3123.47654 | -3.759899628 | 0.1662 | -22.619 | 2.7942E-113 | 3.0257E-111 | |
| AGAP000162 | AGAP000162 | Cystathionine beta-synthase [Source:UniProtKB/TrEMBL;Acc:Q7QEV( | 12899.2081 | -3.755407958 | 0.2193 | -17.121 | 1.0285E-65 | 3.67025E-64 | Oenocytoids |
| AGAP013329 | AGAP013329 | nan | 58.5082405 | -3.749058651 | 0.3876 | -9.6719 | 3.97079E-22 | 3.2249E-21 | |
| AGAP013758 | AGAP013758 | xanthine dehydrogenase | 292.99927 | -3.747835889 | 0.2091 | -17.922 | 7.94971E-72 | 3.40428E-70 | |
| AGAP006226 | Aldehyde_oxidase | nan | 176.000978 | -3.746539451 | 0.2621 | -14.293 | 2.4176E-46 | 4.98384E-45 | |
| AGAP002593 | AGAP002593 | outer membrane lipoprotein Blc | 2444.71449 | -3.732204929 | 0.2577 | -14.483 | 1.55164E-47 | 3.3682E-46 | |
| AGAP004367 | AGAP004367 | nan | 6.27641776 | -3.71151124 | 0.7131 | -5.2048 | 1.94256E-07 | 5.8376E-07 | |
| AGAP013028 | AGAP013028 | nan | 1117.68323 | -3.710018318 | 0.2477 | -14.978 | 1.01911E-50 | 2.41232E-49 | |
| AGAP028641 | AGAP028641 | nan | 56.8742437 | -3.703061814 | 0.2931 | -12.633 | 1.38711E-36 | 2.05795E-35 | |
| AGAP007453 | LRIM9 | leucine-rich immune protein (Short) | 4537.20231 | -3.693936772 | 0.2772 | -13.326 | 1.64439E-40 | 2.80649E-39 | Granulocytes |
| AGAP003205 | AGAP003205 | nan | 376.618638 | -3.68920624 | 0.2596 | -14.213 | 7.60358E-46 | 1.55387E-44 | |
| AGAP005717 | LYSC6 | C-type lysozyme (multi-lysozyme domain protein) | 22.5284384 | -3.686764021 | 0.4394 | -8.3901 | 4.85853E-17 | 3.00935E-16 | |
| AGAP003730 | AGAP003730 | neutral ceramidase | 896.204879 | -3.68553131 | 0.2252 | -16.367 | 3.29211E-60 | 1.02023E-58 | |
| AGAP007764 | AGAP007764 | nan | 8.99746337 | -3.67369365 | 0.65 | -5.6516 | 1.58993E-08 | 5.29658E-08 | |
| AGAP004940 | AGAP004940 | nan | 382.503901 | -3.672119957 | 0.2498 | -14.698 | 6.62693E-49 | 1.47246E-47 | |
| AGAP008193 | AGAP008193 | Nidogen (entactin) [Source:UniProtKB/TrEMBL;Acc:A0A1S4GXK2] | 462.638259 | -3.671374459 | 0.2146 | -17.105 | 1.35951E-65 | 4.8332E-64 | |
| AGAP013403 | AGAP013403 | sodium-coupled monocarboxylate transporter 2 | 17.912668 | -3.667844459 | 0.5948 | -6.1668 | 6.9691E-10 | 2.65921E-09 | |
| AGAP006227 | AGAP006227 | alpha esterase | 125.043869 | -3.663923834 | 0.2663 | -13.756 | 4.66562E-43 | 8.72119E-42 | |
| AGAP011373 | AGAP011373 | nan | 3.74438369 | -3.659337389 | 0.9467 | -3.8654 | 0.000110923 | 0.000243818 | |
| AGAP005645 | AGAP005645 | dehydrogenase/reductase SDR family member 11 precursor | 302.388965 | -3.652840205 | 0.4207 | -8.6835 | 3.83873E-18 | 2.54144E-17 | |
| AGAP011507 | AGAP011507 | nan | 904.647956 | -3.646701855 | 0.2315 | -15.755 | 6.31545E-56 | 1.77077E-54 | |
| AGAP006376 | PDP1 | PAR-domain protein 1 | 433.909902 | -3.645096136 | 0.3953 | -9.2217 | 2.92424E-20 | 2.17266E-19 | |
| AGAP011349 | AGAP011349 | GABA-gated chloride channel | 4.25918733 | -3.634551722 | 0.8909 | -4.0795 | 4.51267E-05 | 0.000104098 | |
| AGAP013039 | AGAP013039 | nan | 14.0975022 | -3.614229167 | 0.5779 | -6.2542 | 3.9963E-10 | 1.56313E-09 | |
| AGAP003000 | AGAP003000 | MFS transporter, PCFT/HCP family, solute carrier family 46 (folate tra | 2287.37878 | -3.613250741 | 0.1782 | -20.277 | 2.04251E-91 | 1.35511E-89 | |
| AGAP000351 | GPRNPY1 | neuropeptide Y receptor 1 | 643.541009 | -3.602029942 | 0.1974 | -18.245 | 2.254E-74 | 1.02584E-72 | |
| AGAP006183 | AGAP006183 | slit protein | 37.8090721 | -3.590155032 | 0.4034 | -8.9004 | 5.56703E-19 | 3.84791E-18 | |
| AGAP000728 | AGAP000728 | wengen | 326.556965 | -3.581993277 | 0.2455 | -14.588 | 3.33079E-48 | 7.36605E-47 | |
| AGAP002890 | AGAP002890 | Lipid storage droplets surface-binding protein 1 | 5661.5455 | -3.579754393 | 0.2174 | -16.466 | 6.39551E-61 | 2.02188E-59 | Granulocytes |
| AGAP012957 | CYP4D17 | cytochrome P450 | 153.399151 | -3.57179938 | 0.2777 | -12.86 | 7.5119E-38 | 1.17168E-36 | |
| AGAP009498 | AGAP009498 | solute carrier family 17 (anion/sugar transporter), member 5 | 4273.27844 | -3.570169735 | 0.1863 | -19.159 | 8.07499E-82 | 4.72513E-80 | |
| AGAP012761 | AGAP012761 | Synaptic vesicle protein | 182.275027 | -3.56445447 | 0.1931 | -18.458 | 4.5353E-76 | 2.20243E-74 | |
| AGAP007599 | AGAP007599 | monolysocardiolipin acyltransferase | 3128.34491 | -3.556441965 | 0.1645 | -21.613 | 1.3503E-103 | 1.2472E-101 | |
| AGAP010794 | AGAP010794 | nan | 304.73046 | -3.539026458 | 0.2496 | -14.177 | 1.27206E-45 | 2.56619E-44 | |
| AGAP010726 | nan | nan | 35.5277234 | -3.532529829 | 0.3617 | -9.7655 | 1.58251E-22 | 1.30436E-21 | |
| AGAP005459 | CPR16 | cuticular protein RR-1 family 16 | 438.759887 | -3.526620378 | 0.2208 | -15.975 | 1.92174E-57 | 5.6933E-56 | |
| AGAP003713 | AGAP003713 | nan | 63.6270203 | -3.525856486 | 0.3897 | -9.0467 | 1.47364E-19 | 1.05255E-18 | |
| AGAP012570 | AGAP012570 | nan | 53.1128837 | -3.502552663 | 0.412 | -8.5009 | 1.88068E-17 | 1.19958E-16 | |
| AGAP001791 | AGAP001791 | nan | 331.958159 | -3.502290928 | 0.3097 | -11.308 | 1.20247E-29 | 1.37816E-28 | |
| AGAP005866 | AGAP005866 | glutamate decarboxylase | 15.1689666 | -3.49578152 | 0.5082 | -6.8789 | 6.03302E-12 | 2.69882E-11 | |

| | | | | | | | | | |
|---|---|---|---|---|---|---|---|---|---|
| AGAP001601 | AGAP001601 | Ser/Thr protein phosphatase/nucleotidase | 774.104306 | -3.468240411 | 0.1658 | -20.924 | 3.21599E-97 | 2.42382E-95 | |
| AGAP013226 | AGAP013226 | nan | 1644.12043 | -3.457137788 | 0.1297 | -26.656 | 1.527E-156 | 2.7664E-154 | |
| AGAP012395 | AGAP012395 | peptide-methionine (S)-S-oxide reductase | 3850.14615 | -3.449976302 | 0.2614 | -13.2 | 8.71672E-40 | 1.44578E-38 | |
| AGAP002629 | AGAP002629 | nan | 1275.01367 | -3.44926394 | 0.8126 | -4.2449 | 2.18681E-05 | 5.22096E-05 | |
| AGAP005563 | Tret1 | facilitated trehalose transporter Tret1 | 40579.9769 | -3.441113178 | 0.204 | -16.867 | 7.90767E-64 | 2.66065E-62 | Granulocytes |
| AGAP008558 | AGAP008558 | nan | 4.4651699 | -3.436978214 | 0.8318 | -4.1321 | 3.59406E-05 | 8.37074E-05 | |
| AGAP000604 | AGAP000604 | nan | 4.84913532 | -3.433371312 | 0.9045 | -3.7957 | 0.000147222 | 0.000317969 | |
| AGAP001015 | AGAP001015 | notch gene homolog 1 | 3328.58 | -3.431146181 | 0.2258 | -15.196 | 3.76173E-52 | 9.30165E-51 | |
| AGAP007755 | AGAP007755 | nan | 6.01877295 | -3.428871874 | 0.8028 | -4.2713 | 1.94304E-05 | 4.67331E-05 | |
| AGAP006729 | AGAP006729 | Ester hydrolase C11orf54 | 5111.62935 | -3.413082857 | 0.1268 | -26.924 | 1.1576E-159 | 2.3204E-157 | |
| AGAP002625 | CTL9 | C-type lectin (CTL) | 53.999118 | -3.410111836 | 0.2862 | -11.915 | 9.87787E-33 | 1.28181E-31 | |
| AGAP003501 | AGAP003501 | Lipase [Source:UniProtKB/TrEMBL;Acc:Q5TVS6] | 3.37367287 | -3.398272737 | 0.9266 | -3.6674 | 0.000244995 | 0.000513823 | |
| AGAP010162 | AGAP010162 | nan | 4.9842719 | -3.383001571 | 0.8178 | -4.1369 | 3.52016E-05 | 8.21282E-05 | |
| AGAP004156 | AGAP004156 | synaptic vesicle protein | 160.254657 | -3.380372078 | 0.1899 | -17.797 | 7.42866E-71 | 3.11046E-69 | |
| AGAP003209 | AGAP003209 | C-4 methylsterol oxidase | 81.4206687 | -3.376664049 | 0.2654 | -12.724 | 4.33271E-37 | 6.57303E-36 | |
| AGAP003106 | Nep4 | neprilysin, neutral endopeptidase 4 | 217.721818 | -3.374218543 | 0.207 | -16.304 | 9.2798E-60 | 2.85703E-58 | |
| AGAP001106 | AGAP001106 | Disconnected protein | 138.711785 | -3.372689615 | 0.3356 | -10.05 | 9.14746E-24 | 7.98686E-23 | |
| AGAP006396 | AGAP006396 | phospholipase B, plb1 | 9.53579431 | -3.368509407 | 0.7363 | -4.575 | 4.76107E-06 | 1.23551E-05 | |
| AGAP009751 | ANCE2 | angiotensin-converting enzyme 2 | 1390.42849 | -3.364251639 | 0.1747 | -19.261 | 1.14528E-82 | 6.74354E-81 | |
| AGAP004203 | Vg | vitellogenin | 80.8811125 | -3.3527186 | 0.2805 | -11.954 | 6.1595E-33 | 8.04836E-32 | |
| AGAP012851 | AGAP012851 | Aldo-keto reductase family 1, member C3 | 2689.50054 | -3.350926014 | 0.1899 | -17.65 | 1.02133E-69 | 4.0771E-68 | Oenocytoids |
| AGAP002091 | AGAP002091 | phosphoribosylformylglycinamidine synthase | 5133.69277 | -3.339265688 | 0.2779 | -12.015 | 2.95805E-33 | 3.91402E-32 | |
| AGAP011460 | AGAP011460 | salivary cysteine-rich protein | 213.407015 | -3.339086963 | 0.2451 | -13.625 | 2.84728E-42 | 5.17842E-41 | |
| AGAP012351 | nan | nan | 408.858941 | -3.338581709 | 0.2222 | -15.026 | 4.99819E-51 | 1.18909E-49 | |
| AGAP012341 | AGAP012341 | nan | 12.32807 | -3.337645667 | 0.5395 | -6.1861 | 6.16677E-10 | 2.36263E-09 | |
| AGAP009551 | AGAP009551 | sulfotransferase (Sult) | 13.6902857 | -3.332394949 | 0.5482 | -6.079 | 1.20905E-09 | 4.51642E-09 | |
| AGAP011223 | AGAP011223 | nan | 5276.54106 | -3.326940033 | 0.1604 | -20.743 | 1.42583E-95 | 1.04943E-93 | |
| AGAP002868 | CYP6P1 | cytochrome P450 | 143.40991 | -3.323546926 | 0.2199 | -15.112 | 1.34727E-51 | 3.28824E-50 | |
| AGAP004936 | AGAP004936 | nan | 58521.4869 | -3.319925676 | 0.169 | -19.649 | 5.84364E-86 | 3.69483E-84 | Prohemocytes |
| AGAP007028 | AGAP007028 | glucosyl/glucuronosyl transferases | 366.712893 | -3.317281126 | 0.1607 | -20.644 | 1.11306E-94 | 8.06629E-93 | |
| AGAP011106 | AGAP011106 | nan | 136.410365 | -3.303921891 | 0.3272 | -10.099 | 5.60304E-24 | 4.94717E-23 | |
| AGAP002347 | AGAP002347 | N-acetylglucosamine-6-phosphate deacetylase [Source:UniProtKB/Tr | 2255.65884 | -3.301828055 | 0.2075 | -15.909 | 5.44952E-57 | 1.59441E-55 | |
| AGAP007598 | AGAP007598 | nan | 25.3472269 | -3.275270777 | 0.4455 | -7.3524 | 1.94707E-13 | 9.61391E-13 | |
| AGAP028204 | AGAP028204 | Alpha-mannosidase [Source:UniProtKB/TrEMBL;Acc:A0A1S4HD42] | 17.4900863 | -3.269261159 | 0.5608 | -5.8298 | 5.54826E-09 | 1.93522E-08 | |
| AGAP003580 | AGAP003580 | nan | 14020.0337 | -3.260376232 | 0.1855 | -17.58 | 3.48911E-69 | 1.36962E-67 | |
| AGAP001124 | AGAP001124 | Aminomethyltransferase [Source:UniProtKB/TrEMBL;Acc:Q7PWZ1] | 357.536681 | -3.244512534 | 0.2845 | -11.404 | 3.99122E-30 | 4.71193E-29 | |
| AGAP003542 | AGAP003542 | Epoxide hydrolase 2, cytoplasmic | 310.598056 | -3.234909749 | 0.2424 | -13.347 | 1.22987E-40 | 2.10283E-39 | |
| AGAP028901 | SSU_rRNA_eukary | Eukaryotic small subunit ribosomal RNA [Source:RFAM;Acc:RF01960 | 15.2663413 | -3.225831598 | 0.712 | -4.5309 | 5.87297E-06 | 1.50843E-05 | |
| AGAP005869 | AGAP005869 | nan | 38.8862209 | -3.221163782 | 0.4054 | -7.9458 | 1.9295E-15 | 1.09439E-14 | |
| AGAP004107 | AGAP004107 | nan | 5.57585964 | -3.218920578 | 0.7526 | -4.2769 | 1.89541E-05 | 4.56554E-05 | |
| AGAP006224 | AGAP006224 | aldehyde oxidase | 290.449108 | -3.21716727 | 0.2071 | -15.537 | 1.94911E-54 | 5.21663E-53 | |
| AGAP004731 | AGAP004731 | secretory phospholipase A2 | 1220.10474 | -3.216543734 | 0.2474 | -13.001 | 1.20116E-38 | 1.91799E-37 | |
| AGAP028056 | AGAP028056 | nan | 73.6524111 | -3.200180676 | 0.2355 | -13.588 | 4.69321E-42 | 8.3899E-41 | |
| AGAP001856 | PER | period circadian protein | 1765.52282 | -3.184496564 | 0.1698 | -18.755 | 1.76032E-78 | 9.26479E-77 | |
| AGAP028649 | AGAP028649 | nan | 161.09642 | -3.178349482 | 0.2017 | -15.76 | 5.86927E-56 | 1.65058E-54 | |
| AGAP012394 | AGAP012394 | peptide-methionine (S)-S-oxide reductase | 4493.91225 | -3.172243861 | 0.2496 | -12.711 | 5.12128E-37 | 7.7444E-36 | Granulocytes |
| AGAP004160 | AGAP004160 | nan | 172.689128 | -3.163101787 | 0.3344 | -9.4595 | 3.09405E-21 | 2.417E-20 | |
| AGAP004531 | AGAP004531 | cathepsin B precursor | 6.10946575 | -3.158939631 | 0.7386 | -4.2768 | 1.8958E-05 | 4.56554E-05 | |
| AGAP011051 | AGAP011051 | aldehyde reductase | 295.973218 | -3.158326014 | 0.2415 | -13.078 | 4.37039E-39 | 7.08665E-38 | |
| AGAP000033 | AGAP000033 | Protein retinal degeneration B | 370.801971 | -3.15519594 | 0.2017 | -15.645 | 3.58205E-55 | 9.78158E-54 | |
| AGAP008304 | AGAP008304 | 3',5'-cyclic-nucleotide phosphodiesterase | 1039.83173 | -3.153481463 | 0.2119 | -14.884 | 4.16668E-50 | 9.57421E-49 | |
| AGAP007633 | AGAP007633 | solute carrier family 36 (proton-coupled amino acid transporter) | 14609.7054 | -3.145660971 | 0.1572 | -20.017 | 3.92245E-89 | 2.54851E-87 | |
| AGAP012280 | AGAP012280 | nan | 266.11271 | -3.145504714 | 0.192 | -16.387 | 2.37033E-60 | 7.41891E-59 | |
| AGAP001554 | AGAP001554 | nan | 202.090158 | -3.126662248 | 0.1992 | -15.695 | 1.63043E-55 | 4.53105E-54 | |
| AGAP028956 | SSU_rRNA_eukary | Eukaryotic small subunit ribosomal RNA [Source:RFAM;Acc:RF01960 | 1718.10723 | -3.125931303 | 0.7941 | -3.9364 | 8.27201E-05 | 0.000185284 | |
| AGAP005009 | AGAP005009 | Pyrroline-5-carboxylate reductase [Source:UniProtKB/TrEMBL;Acc:Q | 2617.04663 | -3.123263198 | 0.2089 | -14.953 | 1.49041E-50 | 3.50154E-49 | |
| AGAP011476 | AGAP011476 | nan | 20.2689356 | -3.116272616 | 0.489 | -6.3723 | 1.86267E-10 | 7.47049E-10 | |
| AGAP002720 | AGAP002720 | cathepsin O | 881.741456 | -3.106508609 | 0.2002 | -15.519 | 2.58646E-54 | 6.82551E-53 | |
| AGAP010885 | AGAP010885 | nan | 421.508745 | -3.105294624 | 0.2845 | -10.915 | 9.8121E-28 | 1.02825E-26 | |
| AGAP003587 | AGAP003587 | sodium-independent sulfate anion transporter | 1718.75882 | -3.097151065 | 0.1639 | -18.901 | 1.12606E-79 | 6.16781E-78 | |
| AGAP011462 | AGAP011462 | nan | 4.86235738 | -3.096174854 | 0.7421 | -4.1723 | 3.015E-05 | 7.09398E-05 | |
| AGAP006725 | COEAE3H | carboxylesterase alpha esterase | 407.087548 | -3.091662668 | 0.2244 | -13.779 | 3.41677E-43 | 6.43788E-42 | |
| AGAP004095 | AGAP004095 | nan | 174.525564 | -3.075839582 | 0.2239 | -13.736 | 6.14185E-43 | 1.14127E-41 | |
| AGAP001635 | AGAP001635 | sodium-coupled monocarboxylate transporter 2 | 204.45657 | -3.07551775 | 0.1827 | -16.835 | 1.35364E-63 | 4.53832E-62 | |
| AGAP005223 | AGAP005223 | Lipocln_cytosolic_FA-bd_dom domain-containing protein [Source:Ur | 165.007244 | -3.065980036 | 0.2315 | -13.242 | 5.00596E-40 | 8.39167E-39 | |
| AGAP007589 | AGAP007589 | glucosyl/glucuronosyl transferases | 621.720554 | -3.057865242 | 0.1706 | -17.921 | 8.02864E-72 | 3.42253E-70 | |
| AGAP003500 | AGAP003500 | Lipase [Source:UniProtKB/TrEMBL;Acc:Q7QH37] | 9.20282967 | -3.051704285 | 0.5388 | -5.6644 | 1.47533E-08 | 4.92876E-08 | |
| AGAP005456 | CPR15 | cuticular protein RR-1 family 15 | 34.5987538 | -3.035623721 | 0.321 | -9.4572 | 3.16299E-21 | 2.46677E-20 | |
| AGAP000313 | AGAP000313 | alanine-glyoxylate aminotransferase 2-like | 4145.07087 | -3.034879671 | 0.1817 | -16.704 | 1.22334E-62 | 4.00178E-61 | |
| AGAP013365 | AGAP013365 | nan | 6527.8282 | -3.031818519 | 0.1594 | -19.022 | 1.11373E-80 | 6.32073E-79 | Granulocytes |
| AGAP013453 | AGAP013453 | nan | 12.7575822 | -3.029051293 | 0.5309 | -5.705 | 1.16354E-08 | 3.92613E-08 | |
| AGAP008289 | AGAP008289 | nan | 10.3311855 | -3.024651458 | 0.6155 | -4.9139 | 8.92716E-07 | 2.49934E-06 | |
| AGAP010633 | nan | nan | 1204.37305 | -3.022523715 | 0.2197 | -13.756 | 4.65989E-43 | 8.72119E-42 | |
| AGAP008225 | AGAP008225 | trehalose 6-phosphate phosphatase | 5167.64302 | -3.021205186 | 0.2138 | -14.128 | 2.54971E-45 | 5.11082E-44 | |
| AGAP013231 | AGAP013231 | Nitrilase homolog 2 | 559.870424 | -3.018276405 | 0.2323 | -12.995 | 1.31212E-38 | 2.08809E-37 | |
| AGAP011104 | AGAP011104 | nan | 427.715002 | -3.01468997 | 0.2914 | -10.346 | 4.36944E-25 | 4.06764E-24 | |
| AGAP008214 | CYP6M4 | cytochrome P450 | 15.9113498 | -3.013627712 | 0.5612 | -5.3704 | 7.85622E-08 | 2.44349E-07 | |
| AGAP001597 | AGAP001597 | nan | 4.84873071 | -3.010538762 | 0.771 | -3.905 | 9.42364E-05 | 0.000209535 | |
| AGAP008013 | AGAP008013 | nan | 7263.01289 | -3.010012835 | 0.2617 | -11.5 | 1.31937E-30 | 1.58543E-29 | |
| AGAP003490 | AGAP003490 | alanine-glyoxylate aminotransferase | 795.275596 | -3.007138647 | 0.1399 | -21.496 | 1.6813E-102 | 1.4943E-100 | Oenocytoids |
| AGAP001410 | AGAP001410 | nan | 15.4557166 | -2.998600332 | 0.4252 | -7.0526 | 1.75558E-12 | 8.14747E-12 | |
| AGAP000690 | AGAP000690 | nan | 279.405826 | -2.984546954 | 0.242 | -12.332 | 6.07351E-35 | 8.5785E-34 | |
| AGAP012350 | nan | nan | 1796.16641 | -2.983440327 | 0.1652 | -18.064 | 6.09658E-73 | 2.70924E-71 | |
| AGAP009029 | AGAP009029 | Protein real-time [Source:UniProtKB/TrEMBL;Acc:A0A1S4GZW6] | 79.806766 | -2.981978623 | 0.279 | -10.689 | 1.14734E-26 | 1.14746E-25 | |
| AGAP008712 | AGAP008712 | Solute carrier organic anion transporter family member [Source:UniF | 112.559908 | -2.978096348 | 0.2888 | -10.313 | 6.15574E-25 | 5.6634E-24 | |
| AGAP011648 | AGAP011648 | nan | 11.7237491 | -2.97673365 | 0.5862 | -5.0782 | 3.81005E-07 | 1.11163E-06 | |
| AGAP010968 | CLIPA9 | CLIP-domain serine protease | 10945.0038 | -2.971260035 | 0.2474 | -12.012 | 3.06453E-33 | 4.04922E-32 | Dividing Granulocytes |
| AGAP012352 | AGAP012352 | Niemann-Pick C2 protein | 5006.94598 | -2.970282005 | 0.213 | -13.947 | 3.27275E-44 | 6.34416E-43 | Dividing Granulocytes |
| AGAP005175 | AGAP005175 | acetyl-CoA carboxylase / biotin carboxylase | 23704.2072 | -2.959842964 | 0.2158 | -13.714 | 8.40361E-43 | 1.54932E-41 | |
| AGAP028439 | AGAP028439 | nan | 5256.99719 | -2.953650419 | 0.1848 | -15.982 | 1.71359E-57 | 5.09267E-56 | Granulocytes |
| AGAP001600 | AGAP001600 | Ser/Thr protein phosphatase/nucleotidase | 104.140397 | -2.952627152 | 0.2597 | -11.37 | 5.91826E-30 | 6.9262E-29 | |
| AGAP012953 | AGAP012953 | epidermal retinal dehydrogenase | 1187.39227 | -2.94249174 | 0.1982 | -14.85 | 6.99271E-50 | 1.59899E-48 | |
| AGAP013431 | nan | nan | 136.457476 | -2.940686527 | 0.2337 | -12.582 | 2.65475E-36 | 3.90178E-35 | |
| AGAP002323 | AGAP002323 | nan | 554.594019 | -2.925967201 | 0.1587 | -18.44 | 6.30938E-76 | 3.01729E-74 | |

| | | | | | | | | | |
|---|---|---|---|---|---|---|---|---|---|
| AGAP003962 | AGAP003962 | WSCD family member AGAP003962 | 21.0022062 | -2.917402552 | 0.3655 | -7.9829 | 1.42915E-15 | 8.17985E-15 | |
| AGAP009922 | AGAP009922 | salivary purine nucleosidase | 430.34796 | -2.915677883 | 0.1489 | -19.586 | 2.05101E-85 | 1.28817E-83 | |
| AGAP003993 | AGAP003993 | nan | 6.7600248 | -2.911199701 | 0.7979 | -3.6488 | 0.00026348 | 0.00055002 | |
| AGAP006369 | CPR144 | cuticular protein RR-2 family 144 | 19.9078638 | -2.909985591 | 0.5092 | -5.7152 | 1.09544E-08 | 3.70561E-08 | |
| AGAP003453 | AGAP003453 | nan | 5931.74076 | -2.909971112 | 0.1682 | -17.302 | 4.57113E-67 | 1.66917E-65 | |
| AGAP009128 | AGAP009128 | mitochondrial carnitine/acylcarnitine carrier protein | 2204.91283 | -2.909484598 | 0.1843 | -15.786 | 3.85496E-56 | 1.10053E-54 | |
| AGAP008596 | AGAP008596 | long-chain-fatty-acid--CoA ligase ACSBG | 10721.8263 | -2.903218293 | 0.241 | -12.048 | 1.98212E-33 | 2.64124E-32 | |
| AGAP005674 | AGAP005674 | nan | 2631.86204 | -2.888483883 | 0.2026 | -14.258 | 3.98509E-46 | 8.19729E-45 | |
| AGAP003015 | AGAP003015 | Argininosuccinate synthase [Source:UniProtKB/Swiss-Prot;Acc:Q7PR3 | 905.992913 | -2.883968137 | 0.2339 | -12.329 | 6.3048E-35 | 8.87855E-34 | |
| AGAP006740 | AGAP006740 | dolichyl-phosphate mannosyltransferase polypeptide 2, regulatory su | 1307.9002 | -2.880732197 | 0.2493 | -11.553 | 7.08769E-31 | 8.57165E-30 | |
| AGAP007752 | AGAP007752 | nan | 40.4594783 | -2.87946621 | 0.3539 | -8.1358 | 4.09318E-16 | 2.41919E-15 | |
| AGAP002610 | AGAP002610 | nan | 5.03221191 | -2.863640478 | 0.72 | -3.9771 | 6.97499E-05 | 0.000157619 | |
| AGAP005301 | AGAP005301 | nan | 45.0072735 | -2.856312215 | 0.3721 | -7.6761 | 1.63972E-14 | 8.73252E-14 | |
| AGAP008213 | CYP6M3 | cytochrome P450 | 104.606549 | -2.852590248 | 0.3069 | -9.2951 | 1.47038E-20 | 1.10819E-19 | |
| AGAP008207 | CYP6Y2 | cytochrome P450 | 86.035535 | -2.847535037 | 0.2449 | -11.628 | 2.96365E-31 | 3.65452E-30 | |
| AGAP008783 | AGAP008783 | Arginase [Source:UniProtKB/TrEMBL;Acc:A0A1S4H002] | 1167.31795 | -2.845130597 | 0.1847 | -15.404 | 1.54982E-53 | 4.01123E-52 | |
| AGAP008892 | AGAP008892 | nan | 417.908217 | -2.843210636 | 0.2159 | -13.169 | 1.3275E-39 | 2.18643E-38 | |
| AGAP010142 | Dat | dopamine N-acetyltransferase | 2530.63992 | -2.838286283 | 0.1599 | -17.747 | 1.82768E-70 | 7.58525E-69 | |
| AGAP002789 | AGAP002789 | nan | 5.70084052 | -2.837506957 | 0.6762 | -4.196 | 2.71662E-05 | 6.41919E-05 | |
| AGAP003303 | AGAP003303 | nan | 109.004999 | -2.836879562 | 0.2309 | -12.288 | 1.04903E-34 | 1.46197E-33 | |
| AGAP002672 | AGAP002672 | nan | 668.892011 | -2.835433553 | 0.1838 | -15.429 | 1.04764E-53 | 2.72647E-52 | |
| AGAP006699 | AGAP006699 | hairy and enhancer of split, invertebrate | 858.698334 | -2.832903216 | 0.1086 | -26.077 | 6.587E-150 | 1.0699E-147 | |
| AGAP012387 | TOLL6 | TOLL-like receptor 6 | 77.3245835 | -2.828099213 | 0.3668 | -7.7112 | 1.24652E-14 | 6.70671E-14 | |
| AGAP001590 | AGAP001590 | nan | 302.039014 | -2.823836965 | 0.2181 | -12.949 | 2.37916E-38 | 3.76076E-37 | |
| AGAP013189 | AGAP013189 | nan | 3067.58971 | -2.817727508 | 0.1822 | -15.466 | 5.85753E-54 | 1.53715E-52 | |
| AGAP008632 | AGAP008632 | alpha-aminoadipic semialdehyde synthase | 3981.68771 | -2.812682021 | 0.1597 | -17.613 | 1.96282E-69 | 7.73712E-68 | Dividing Granulocytes |
| AGAP011459 | AGAP011459 | nan | 33.1253239 | -2.812048835 | 0.3174 | -8.8599 | 8.00967E-19 | 5.47598E-18 | |
| AGAP008288 | TIM | timeless | 7798.30818 | -2.805276006 | 0.1311 | -21.393 | 1.5406E-101 | 1.3315E-99 | |
| AGAP004270 | AGAP004270 | hexosaminidase | 24.5336753 | -2.795173226 | 0.4168 | -6.7071 | 1.98595E-11 | 8.53933E-11 | |
| AGAP028177 | AGAP028177 | nan | 7.9695215 | -2.79499991 | 0.609 | -4.5894 | 4.446E-06 | 1.15774E-05 | |
| AGAP009330 | AGAP009330 | Troponin C isoform 4'' | 63.4230335 | -2.788674109 | 0.3713 | -7.511 | 5.86569E-14 | 2.99192E-13 | |
| AGAP008208 | CYP6Y1 | cytochrome P450 | 263.444328 | -2.785155 | 0.2155 | -12.922 | 3.36308E-38 | 5.27181E-37 | |
| AGAP000262 | AGAP000262 | nan | 12.6655084 | -2.784010858 | 0.5806 | -4.7949 | 1.62755E-06 | 4.43795E-06 | |
| AGAP007601 | AGAP007601 | nan | 537.939718 | -2.781081293 | 0.2205 | -12.614 | 1.76954E-36 | 2.61298E-35 | |
| AGAP013770 | AGAP013770 | nan | 73.4854943 | -2.780319016 | 0.364 | -7.6391 | 2.18759E-14 | 1.15523E-13 | |
| AGAP001520 | AGAP001520 | protein phosphatase 1, regulatory (inhibitor) subunit 3 | 3229.13407 | -2.778218252 | 0.153 | -18.163 | 1.01454E-73 | 4.5514E-72 | |
| AGAP003849 | AGAP003849 | nan | 13.1587734 | -2.777089907 | 0.5736 | -4.8415 | 1.28833E-06 | 3.54997E-06 | |
| AGAP029107 | AGAP029107 | nan | 1012.84628 | -2.772678791 | 0.237 | -11.698 | 1.29917E-31 | 1.6147E-30 | |
| AGAP005496 | LRIM12 | leucine-rich immune protein (Short) | 203.283658 | -2.766593335 | 0.3225 | -8.5793 | 9.54397E-18 | 6.18389E-17 | |
| AGAP028029 | AGAP028029 | nan | 5.02626872 | -2.762000872 | 0.7591 | -3.6384 | 0.000274325 | 0.000570386 | |
| AGAP011714 | nan | nan | 439.767191 | -2.761655121 | 0.1904 | -14.505 | 1.12542E-47 | 2.46572E-46 | |
| AGAP006158 | AGAP006158 | nan | 36.2486886 | -2.760644472 | 0.3219 | -8.5766 | 9.76845E-18 | 6.32498E-17 | |
| AGAP008009 | AGAP008009 | RYK receptor-like tyrosine kinase [Source:UniProtKB/TrEMBL;Acc:A0/ | 4.41505268 | -2.760315422 | 0.7329 | -3.7665 | 0.000165534 | 0.000355562 | |
| AGAP005103 | AGAP005103 | nan | 477.425162 | -2.759866256 | 0.2061 | -13.394 | 6.57512E-41 | 1.13689E-39 | |
| AGAP004976 | PPO8 | prophenoloxidase 8 | 43.90878 | -2.747668438 | 0.2434 | -11.287 | 1.5251E-29 | 1.74369E-28 | |
| AGAP007801 | AGAP007801 | vrille | 2468.89931 | -2.747094542 | 0.0853 | -32.212 | 1.2181E-227 | 5.738E-225 | |
| AGAP028400 | AGAP028400 | nan | 5.69646929 | -2.744756716 | 0.7615 | -3.6042 | 0.000313127 | 0.000645648 | |
| AGAP008142 | nan | nan | 22.9964762 | -2.744114338 | 0.4755 | -5.7714 | 7.8605E-09 | 2.69679E-08 | |
| AGAP001480 | AGAP001480 | nan | 1445.71215 | -2.743518159 | 0.097 | -28.277 | 6.6421E-176 | 1.7879E-173 | |
| AGAP012295 | CYP9L1 | cytochrome P450 | 635.65781 | -2.738935645 | 0.1749 | -15.664 | 2.67983E-55 | 7.36056E-54 | |
| AGAP005776 | PDH | pigment dispersing hormone | 28.7151958 | -2.738428116 | 0.5075 | -5.3955 | 6.83162E-08 | 2.13823E-07 | |
| AGAP002641 | AGAP002641 | nan | 9.02264866 | -2.729059706 | 0.6329 | -4.312 | 1.61757E-05 | 3.92154E-05 | |
| AGAP004534 | AGAP004534 | cathepsin B precursor | 86.8145119 | -2.728691805 | 0.2043 | -13.355 | 1.10832E-40 | 1.9019E-39 | |
| AGAP007094 | AGAP007094 | nan | 7.26679985 | -2.727472848 | 0.6001 | -4.5451 | 5.49007E-06 | 1.41587E-05 | |
| AGAP000968 | AGAP000968 | nan | 94.3562328 | -2.725815367 | 0.2311 | -11.795 | 4.12248E-32 | 5.22015E-31 | |
| AGAP002341 | AGAP002341 | nan | 1003.57572 | -2.723399398 | 0.1967 | -13.849 | 1.29596E-43 | 2.48155E-42 | |
| AGAP007339 | AGAP007339 | troponin C, isoform 3 | 44.4004595 | -2.720632296 | 0.3134 | -8.6813 | 3.91329E-18 | 2.58898E-17 | |
| AGAP009176 | AGAP009176 | fatty acid synthase, animal type | 33667.3064 | -2.719926417 | 0.1833 | -14.837 | 8.43178E-50 | 1.92338E-48 | |
| AGAP000167 | AGAP000167 | Lipid storage droplets surface-binding protein 2 | 9070.62516 | -2.714439432 | 0.0793 | -34.228 | 9.2256E-257 | 7.9013E-254 | Granulocytes |
| AGAP010269 | AGAP010269 | ANK_REP_REGION domain-containing protein [Source:UniProtKB/TrE | 68.6841714 | -2.713220433 | 0.2536 | -10.698 | 1.0417E-26 | 1.04292E-25 | |
| AGAP013400 | AGAP013400 | nan | 5086.33564 | -2.703161119 | 0.1731 | -15.612 | 6.00712E-55 | 1.62624E-53 | |
| AGAP005739 | nan | nan | 138.026837 | -2.700610582 | 0.4011 | -6.7332 | 1.66021E-11 | 7.17472E-11 | |
| AGAP007647 | AGAP007647 | nan | 3027.80365 | -2.699881035 | 0.1583 | -17.054 | 3.27409E-65 | 1.14241E-63 | |
| AGAP004157 | AGAP004157 | synaptic vesicle protein | 31.6579558 | -2.699466121 | 0.3022 | -8.9333 | 4.13507E-19 | 2.87927E-18 | |
| AGAP005834 | COEJHE2E | carboxylesterase | 258.838753 | -2.687648621 | 0.2812 | -9.5594 | 1.18386E-21 | 9.45181E-21 | |
| AGAP009218 | AGAP009218 | nan | 526.027122 | -2.686723095 | 0.5865 | -4.5807 | 4.63322E-06 | 1.20446E-05 | |
| AGAP002503 | AGAP002503 | 4-coumarate:CoA ligase | 3661.11725 | -2.686283431 | 0.1765 | -15.218 | 2.68018E-52 | 6.67989E-51 | |
| AGAP010969 | AGAP010969 | Roundabout 1 | 62.0145588 | -2.682168675 | 0.2699 | -9.9381 | 2.84093E-23 | 2.41774E-22 | |
| AGAP012216 | AGAP012216 | sterol O-acyltransferase | 169.461239 | -2.681926554 | 0.2115 | -12.682 | 7.48376E-37 | 1.11735E-35 | |
| AGAP007738 | AGAP007738 | nan | 221.051401 | -2.681722644 | 0.2178 | -12.311 | 7.92426E-35 | 1.11093E-33 | |
| AGAP005740 | nan | nan | 41.0247269 | -2.673648343 | 0.4357 | -6.1371 | 8.40181E-10 | 3.18525E-09 | |
| AGAP002268 | AGAP002268 | nan | 4372.36981 | -2.664891922 | 0.2027 | -13.149 | 1.72456E-39 | 2.83049E-38 | |
| AGAP003934 | AGAP003934 | Battenin [Source:UniProtKB/TrEMBL;Acc:Q7Q2B1] | 810.701736 | -2.661543747 | 0.0961 | -27.689 | 9.5586E-169 | 2.1441E-166 | |
| AGAP001127 | AGAP001127 | P37NB protein | 576.584241 | -2.660769671 | 0.1794 | -14.834 | 8.88369E-50 | 2.0167E-48 | |
| AGAP005992 | CYP302A1 | cytochrome P450 | 80.4973131 | -2.651850026 | 0.2698 | -9.8275 | 8.56846E-23 | 7.15E-22 | |
| AGAP000180 | AGAP000180 | phosphoribosylaminoimidazole carboxylase | 5773.79183 | -2.649490911 | 0.2247 | -11.792 | 4.28269E-32 | 5.41574E-31 | |
| AGAP004954 | AGAP004954 | rhythmically expressed gene 2 protein | 313.42621 | -2.649035701 | 0.2208 | -11.995 | 3.77261E-33 | 4.96393E-32 | |
| AGAP002835 | AGAP002835 | alpha-tocopherol transfer protein-like protein | 53.4834947 | -2.646952914 | 0.2813 | -9.4099 | 4.96717E-21 | 3.82631E-20 | |
| AGAP007754 | AGAP007754 | nan | 1841.38919 | -2.645616117 | 0.122 | -21.68 | 3.2013E-104 | 3.0159E-102 | |
| AGAP007165 | AGAP007165 | Late trypsin | 9.33984706 | -2.641139965 | 0.5333 | -4.9521 | 7.34118E-07 | 2.07692E-06 | |
| AGAP006467 | AGAP006467 | nan | 20.2503579 | -2.640444595 | 0.3706 | -7.1241 | 1.04729E-12 | 4.93078E-12 | |
| AGAP003097 | AGAP003097 | yellow protein | 84.2787943 | -2.636689025 | 0.2939 | -8.9702 | 2.95951E-19 | 2.08071E-18 | |
| AGAP011924 | AGAP011924 | Potassium channel subfamily K, invertebrate [Source:UniProtKB/TrEI | 79.1578349 | -2.634037214 | 0.2057 | -12.805 | 1.52893E-37 | 2.3652E-36 | |
| AGAP007711 | AGAP007711 | nan | 6245.8815 | -2.63161604 | 0.1939 | -13.571 | 5.97309E-42 | 1.06175E-40 | Dividing Granulocytes |
| AGAP009770 | GPRCAL1 | putative calcitonin receptor 1 | 370.072892 | -2.631610182 | 0.2542 | -10.351 | 4.14188E-25 | 3.85961E-24 | |
| AGAP028580 | AGAP028580 | nan | 107.348715 | -2.630831885 | 0.2637 | -9.9782 | 1.89803E-23 | 1.62557E-22 | |
| AGAP009414 | AGAP009414 | 1-acyl-sn-glycerol-3-phosphate acyltransferase alpha | 792.162407 | -2.620327636 | 0.1958 | -13.381 | 7.79382E-41 | 1.34233E-39 | |
| AGAP002718 | AGAP002718 | 4-coumarate:CoA ligase isoform 2 [Source:UniProtKB/TrEMBL;Acc:Q | 1450.95555 | -2.616001711 | 0.1703 | -15.357 | 3.18945E-53 | 8.18741E-52 | |
| AGAP007198 | AGAP007198 | nan | 756.12112 | -2.613410195 | 0.1623 | -16.105 | 2.3612E-58 | 7.17575E-57 | |
| AGAP003483 | AGAP003483 | nan | 394.558751 | -2.608053865 | 0.218 | -11.965 | 5.40774E-33 | 7.08572E-32 | |
| AGAP002069 | AGAP002069 | decaprenyl-diphosphate synthase subunit 1 | 36.9606939 | -2.606488166 | 0.3439 | -7.5801 | 3.45261E-14 | 1.78917E-13 | |
| AGAP001779 | AGAP001779 | nan | 200.711143 | -2.606480434 | 0.1899 | -13.728 | 6.92004E-43 | 1.28082E-41 | |
| AGAP028646 | ANCE3 | angiotensin-converting enzyme 3 | 31.9498117 | -2.60620726 | 0.2999 | -8.6916 | 3.57438E-18 | 2.37143E-17 | |
| AGAP000290 | CLIPA27 | CLIP-domain serine protease | 3535.03588 | -2.602632476 | 0.176 | -14.784 | 1.86988E-49 | 4.22449E-48 | Granulocytes |

| | | | | | | | | | |
|---|---|---|---|---|---|---|---|---|---|
| AGAP001524 | AGAP001524 | nan | 71.7487177 | -2.599672448 | 0.2849 | -9.1264 | 7.07987E-20 | 5.16249E-19 | |
| AGAP008982 | AGAP008982 | CzcD (Cation-efflux system membrane protein) [Source:UniProtKB/Tr | 356.806884 | -2.599164944 | 0.229 | -11.352 | 7.28096E-30 | 8.46838E-29 | |
| AGAP007029 | AGAP007029 | glucosyl/glucuronosyl transferases | 527.444528 | -2.597261687 | 0.1792 | -14.493 | 1.35051E-47 | 2.93838E-46 | |
| AGAP010860 | AGAP010860 | sodium-dependent nutrient amino acid transporter 2 | 700.333353 | -2.595959467 | 0.219 | -11.855 | 2.03445E-32 | 2.61838E-31 | |
| AGAP008143 | AGAP008143 | nan | 23.8458732 | -2.595511659 | 0.4224 | -6.145 | 7.99724E-10 | 3.03676E-09 | |
| AGAP007163 | AGAP007163 | 3',5'-cyclic-nucleotide phosphodiesterase | 313.285862 | -2.594609145 | 0.1854 | -13.993 | 1.73114E-44 | 3.38362E-43 | |
| AGAP012477 | AGAP012477 | gamma-butyrobetaine dioxygenase | 43.7310946 | -2.58874021 | 0.3542 | -7.3092 | 2.68647E-13 | 1.31204E-12 | |
| AGAP011322 | AGAP011322 | fibulin 2 | 10.4308474 | -2.588621121 | 0.6504 | -3.9799 | 6.89343E-05 | 0.000155926 | |
| AGAP013537 | AGAP013537 | nan | 15.564936 | -2.586296847 | 0.5132 | -5.0394 | 4.66996E-07 | 1.34915E-06 | |
| AGAP006579 | AGAP006579 | nan | 18.5285262 | -2.586225151 | 0.5121 | -5.0505 | 4.40577E-07 | 1.27595E-06 | |
| AGAP006548 | AGAP006548 | glycine cleavage system H protein | 3029.33721 | -2.585621626 | 0.2382 | -10.854 | 1.91878E-27 | 1.99084E-26 | |
| AGAP002084 | AGAP002084 | glycerol-3-phosphate O-acyltransferase 3/4 | 1244.10441 | -2.578332454 | 0.21 | -12.279 | 1.1747E-34 | 1.62988E-33 | |
| AGAP012307 | AGAP012307 | Spondin-1 | 15732.9862 | -2.571529683 | 0.1781 | -14.441 | 2.84551E-47 | 6.06505E-46 | |
| AGAP005581 | AGAP005581 | 3-hydroxyisobutyrate dehydrogenase [Source:UniProtKB/TrEMBL;Ac | 4012.06571 | -2.565208818 | 0.1729 | -14.836 | 8.61132E-50 | 1.95959E-48 | |
| AGAP001561 | GPRNNA5 | putative GPCR class a orphan receptor 5 | 39.8573453 | -2.56234367 | 0.3002 | -8.5357 | 1.3925E-17 | 8.95477E-17 | |
| AGAP009364 | AGAP009364 | nan | 32.8702336 | -2.56014532 | 0.4466 | -5.7327 | 9.88478E-09 | 3.36798E-08 | |
| AGAP005783 | AGAP005783 | nan | 129.288309 | -2.559195545 | 0.2169 | -11.798 | 4.00764E-32 | 5.08156E-31 | |
| AGAP011427 | nan | nan | 197.796839 | -2.55540795 | 0.2501 | -10.217 | 1.67081E-24 | 1.50054E-23 | |
| AGAP010632 | nan | nan | 559.977344 | -2.549597648 | 0.2491 | -10.236 | 1.37148E-24 | 1.23643E-23 | |
| AGAP004600 | nan | nan | 197.437918 | -2.544823074 | 0.1531 | -16.62 | 4.99063E-62 | 1.61016E-60 | |
| AGAP007920 | AGAP007920 | glucuronosyltransferase | 1090.97589 | -2.540702277 | 0.2074 | -12.249 | 1.69933E-34 | 2.35087E-33 | |
| AGAP009763 | AGAP009763 | nan | 108.051319 | -2.535435227 | 0.2509 | -10.105 | 5.26959E-24 | 4.65711E-23 | |
| AGAP002449 | AGAP002449 | 3-OH androgenic UDPGT | 10.2672766 | -2.535213461 | 0.4894 | -5.1807 | 2.21103E-07 | 6.60646E-07 | |
| AGAP002156 | GPRGNR1 | putative gonadotrophin releasing hormone receptor 1 | 1190.95674 | -2.535035992 | 0.2038 | -12.439 | 1.60396E-35 | 2.31053E-34 | |
| AGAP003759 | AGAP003759 | juvenile hormone-inducible protein | 900.466307 | -2.532817671 | 0.2116 | -11.969 | 5.16558E-33 | 6.7873E-32 | Granulocytes |
| AGAP006698 | AGAP006698 | nan | 20.9079053 | -2.523103693 | 0.371 | -6.8001 | 1.04529E-11 | 4.59101E-11 | |
| AGAP009472 | ABCG20 | ATP-binding cassette transporter (ABC transporter) family G member | 63.036108 | -2.517693555 | 0.275 | -9.1558 | 5.39325E-20 | 3.95407E-19 | |
| AGAP010887 | CPR113 | cuticular protein RR-2 family 113 | 16.3552987 | -2.515612185 | 0.5228 | -4.8114 | 1.49864E-06 | 4.10428E-06 | |
| AGAP009757 | ANCE7 | angiotensin-converting enzyme 7 | 10.4647292 | -2.508544242 | 0.5018 | -4.9994 | 5.75158E-07 | 1.64598E-06 | |
| AGAP003422 | AGAP003422 | endoplasmic reticulum metallopeptidase 1 | 125.955835 | -2.507949863 | 0.2339 | -10.725 | 7.80256E-27 | 7.87866E-26 | |
| AGAP006741 | AGAP006741 | nan | 578.501361 | -2.504956326 | 0.2433 | -10.296 | 7.37979E-25 | 6.75E-24 | |
| AGAP002250 | AGAP002250 | calcium/calmodulin-dependent protein kinase kinase | 527.504932 | -2.503700527 | 0.2193 | -11.418 | 3.41461E-30 | 4.04642E-29 | |
| AGAP012399 | AGAP012399 | maltase | 3023.90855 | -2.503289364 | 0.1176 | -21.292 | 1.356E-100 | 1.1509E-98 | |
| AGAP006569 | AGAP006569 | acetyl-CoA synthetase | 716.52121 | -2.503169264 | 0.1061 | -23.592 | 4.6247E-123 | 5.8092E-121 | |
| AGAP007761 | AGAP007761 | Complement control protein | 14.786335 | -2.502881112 | 0.5086 | -4.9208 | 8.61892E-07 | 2.41807E-06 | |
| AGAP009312 | AGAP009312 | nan | 9.586563 | -2.493802194 | 0.551 | -4.5257 | 6.02074E-06 | 1.54368E-05 | |
| AGAP011750 | AGAP011750 | alpha-N-acetylglucosaminidase | 722.334084 | -2.493652027 | 0.2349 | -10.618 | 2.45678E-26 | 2.42106E-25 | |
| AGAP009347 | AGAP009347 | FK506-binding protein 4/5 | 3624.89028 | -2.484843102 | 0.1689 | -14.711 | 5.48884E-49 | 1.22247E-47 | |
| AGAP009556 | AGAP009556 | nan | 559.315853 | -2.47441434 | 0.1965 | -12.595 | 2.26295E-36 | 3.33114E-35 | |
| AGAP006819 | AGAP006819 | abhydrolase domain-containing protein 1/3 | 631.818128 | -2.470024441 | 0.2032 | -12.156 | 5.29825E-34 | 7.24453E-33 | |
| AGAP002826 | AGAP002826 | nan | 33.9130791 | -2.469666613 | 0.6566 | -3.7611 | 0.000169195 | 0.00036293 | |
| AGAP009896 | AGAP009896 | proton-coupled amino acid transporter | 69.8293157 | -2.466530637 | 0.2596 | -9.5024 | 2.05032E-21 | 1.6232E-20 | |
| AGAP011509 | COEunkn | carboxylesterase | 420.334576 | -2.458427804 | 0.3092 | -7.9515 | 1.84278E-15 | 1.04836E-14 | |
| AGAP000095 | AGAP000095 | anoctamin 5 | 4124.76665 | -2.45804903 | 0.0772 | -31.844 | 1.5845E-222 | 6.7855E-220 | |
| AGAP008557 | AGAP008557 | AMP-binding domain protein | 292.756205 | -2.451986748 | 0.1995 | -12.293 | 9.84826E-35 | 1.37452E-33 | |
| AGAP000698 | AGAP000698 | nan | 981.786796 | -2.446687047 | 0.0813 | -30.098 | 5.1128E-199 | 1.661E-196 | Granulocytes |
| AGAP002109 | AGAP002109 | solute carrier family 31 (copper transporter), member 1 | 5710.06577 | -2.444179463 | 0.2399 | -10.19 | 2.18898E-24 | 1.95844E-23 | |
| AGAP012813 | AGAP012813 | nan | 152.580053 | -2.439718865 | 0.1883 | -12.958 | 2.1264E-38 | 3.36687E-37 | |
| AGAP001884 | AGAP001884 | fumarate hydratase, class II | 11280.3377 | -2.438626332 | 0.1692 | -14.409 | 4.55178E-47 | 9.57194E-46 | Oenocytoids |
| AGAP007662 | AGAP007662 | All-trans/9-cis | 11.9938402 | -2.436478461 | 0.4515 | -5.3959 | 6.81775E-08 | 2.1346E-07 | |
| AGAP005617 | AGAP005617 | nan | 3311.35222 | -2.43499106 | 0.1134 | -21.465 | 3.2861E-102 | 2.8665E-100 | |
| AGAP009609 | AGAP009609 | homogentisate 1,2-dioxygenase | 22.0763605 | -2.427578942 | 0.3998 | -6.0727 | 1.25793E-09 | 4.69501E-09 | |
| AGAP001989 | AGAP001989 | nan | 4244.17765 | -2.419207125 | 0.2014 | -12.009 | 3.18551E-33 | 4.20318E-32 | |
| AGAP003927 | ILP5 | Insulin-like peptide 5 | 32.0637047 | -2.41812528 | 0.5408 | -4.4711 | 7.78321E-06 | 1.969E-05 | |
| AGAP011197 | AGAP011197 | nan | 15631.9393 | -2.417966052 | 0.3117 | -7.7563 | 8.74367E-15 | 4.78641E-14 | Granulocytes |
| AGAP004802 | AGAP004802 | 4-hydroxyphenylpyruvate dioxygenase [Source:UniProtKB/TrEMBL;A | 135.189293 | -2.413744061 | 0.2708 | -8.914 | 4.92069E-19 | 3.41117E-18 | |
| AGAP012391 | AGAP012391 | nan | 24.8363814 | -2.412875336 | 0.3979 | -6.0641 | 1.32681E-09 | 4.93677E-09 | |
| AGAP003579 | AGAP003579 | cadherin-87A | 139.194367 | -2.412126322 | 0.1743 | -13.838 | 1.49847E-43 | 2.84045E-42 | |
| AGAP004620 | AGAP004620 | Envelysin | 15.5825335 | -2.411984736 | 0.4254 | -5.6696 | 1.43097E-08 | 4.79074E-08 | |
| AGAP028196 | AGAP028196 | nan | 266.574124 | -2.409816791 | 0.2173 | -11.09 | 1.404E-28 | 1.53982E-27 | |
| AGAP002891 | GPRMGL4 | putative metabotropic glutamate receptor 4 | 373.811128 | -2.407173492 | 0.2192 | -10.979 | 4.80847E-28 | 5.11871E-27 | |
| AGAP002738 | SCRB5 | Class B Scavenger Receptor (CD36 domain). | 38.217221 | -2.404354095 | 0.3112 | -7.7259 | 1.11032E-14 | 6.01859E-14 | |
| AGAP002761 | AGAP002761 | 3-methylcrotonyl-CoA carboxylase beta subunit | 1711.80514 | -2.404176528 | 0.2023 | -11.881 | 1.4801E-32 | 1.91276E-31 | |
| AGAP000131 | AGAP000131 | nan | 29.2004817 | -2.403920305 | 0.3135 | -7.6688 | 1.73551E-14 | 9.23222E-14 | |
| AGAP011715 | nan | nan | 2052.03365 | -2.403424709 | 0.1053 | -22.815 | 3.2285E-115 | 3.755E-113 | |
| AGAP008688 | AGAP008688 | nan | 3288.38596 | -2.402442913 | 0.1915 | -12.544 | 4.30178E-36 | 6.29302E-35 | |
| AGAP002925 | AGAP002925 | poly(U)-specific endoribonuclease | 6803.74621 | -2.401733656 | 0.2459 | -9.7675 | 1.55225E-22 | 1.28504E-21 | |
| AGAP007095 | AGAP007095 | nan | 8.07568557 | -2.400270434 | 0.5705 | -4.2069 | 2.58844E-05 | 6.13015E-05 | |
| AGAP011231 | AGAP011231 | fibrinogen | 77.3634722 | -2.397781425 | 0.3923 | -6.1117 | 9.85592E-10 | 3.71262E-09 | |
| AGAP000327 | AGAP000327 | tyrosine aminotransferase | 1656.41281 | -2.397077006 | 0.2245 | -10.679 | 1.2738E-26 | 1.27258E-25 | |
| AGAP003358 | AGAP003358 | Sugar transporter SWEET [Source:UniProtKB/TrEMBL;Acc:A0NDW1] | 110.058707 | -2.396761041 | 0.3004 | -7.9789 | 1.47585E-15 | 8.44202E-15 | |
| AGAP011052 | nan | nan | 857.99609 | -2.396236463 | 0.1623 | -14.767 | 2.39414E-49 | 5.38309E-48 | |
| AGAP006637 | AGAP006637 | Oatp58Dc | 12.5468924 | -2.394554131 | 0.6858 | -3.4915 | 0.000480403 | 0.000965212 | |
| AGAP011830 | AGAP011830 | myo-inositol-1-phosphate synthase | 34.555182 | -2.389023549 | 0.3819 | -6.255 | 3.97412E-10 | 1.55547E-09 | |
| AGAP003267 | AGAP003267 | nan | 45.6408423 | -2.385621788 | 0.3007 | -7.9344 | 2.11486E-15 | 1.19592E-14 | |
| AGAP011158 | AGAP011158 | Kynurenine--oxoglutarate transaminase 3 isoform 1 | 2474.65055 | -2.384308505 | 0.1095 | -21.769 | 4.5306E-105 | 4.3554E-103 | Oenocytoids |
| AGAP007664 | AGAP007664 | coiled-coil domain-containing protein 34 | 27.849818 | -2.384306765 | 0.3838 | -6.213 | 5.19672E-10 | 2.00896E-09 | |
| AGAP007675 | AGAP007675 | dynein, axonemal heavy chain | 6.36055081 | -2.380253171 | 0.6441 | -3.6957 | 0.000219288 | 0.000463625 | |
| AGAP000238 | AGAP000238 | nan | 703.056376 | -2.380158991 | 0.2402 | -9.9074 | 3.86679E-23 | 3.27305E-22 | |
| AGAP007781 | AGAP007781 | eukaryotic translation initiation factor 4E-binding protein 1 | 14699.8994 | -2.378865811 | 0.0693 | -34.312 | 5.16E-258 | 5.4014E-255 | |
| AGAP012000 | AGAP012000 | fibrinogen and fibronectin | 4234.79495 | -2.37771306 | 0.1924 | -12.361 | 4.23122E-35 | 6.01242E-34 | Oenocytoids |
| AGAP008012 | AGAP008012 | nan | 596.141968 | -2.377128217 | 0.2183 | -10.888 | 1.3098E-27 | 1.36349E-26 | |
| AGAP000015 | AGAP000015 | proto-oncogene tyrosine-protein kinase ROS | 56.839972 | -2.376981853 | 0.2965 | -8.0166 | 1.08705E-15 | 6.27132E-15 | |
| AGAP003005 | AGAP003005 | serine/threonine-protein kinase SIK | 567.698708 | -2.376750193 | 0.1422 | -16.71 | 1.10866E-62 | 3.63925E-61 | |
| AGAP012742 | AGAP012742 | nan | 8.2364549 | -2.373141712 | 0.5079 | -4.6725 | 2.97616E-06 | 7.89368E-06 | |
| AGAP010645 | AGAP010645 | nan | 625.912603 | -2.372574179 | 0.2409 | -9.8491 | 6.91789E-23 | 5.77778E-22 | |
| AGAP006743 | AGAP006743 | nan | 1458.16669 | -2.372151238 | 0.1901 | -12.479 | 9.73705E-36 | 1.41782E-34 | Oenocytoids |
| AGAP005302 | AGAP005302 | DIRAS family, GTP-binding Ras-like 2 | 68.5889576 | -2.368684796 | 0.2906 | -8.15 | 3.63803E-16 | 2.15288E-15 | |
| AGAP001293 | AGAP001293 | nan | 24.530601 | -2.36509304 | 0.3108 | -7.6086 | 2.77013E-14 | 1.44825E-13 | |
| AGAP009405 | CPAP3-E | cuticular protein | 75.4477584 | -2.363863534 | 0.2379 | -9.9369 | 2.87681E-23 | 2.44607E-22 | |
| AGAP000278 | OBP9 | odorant-binding protein 9 | 136.522372 | -2.361210706 | 0.2298 | -10.276 | 9.01486E-25 | 8.16625E-24 | |
| AGAP009381 | AGAP009381 | Cellular retinaldehyde-binding protein [Source:UniProtKB/TrEMBL;A | 25.722645 | -2.359419209 | 0.5046 | -4.6756 | 2.93042E-06 | 7.77675E-06 | |
| AGAP007692 | SRPN14 | serine protease inhibitor (serpin) 14 | 234.911196 | -2.359105557 | 0.2157 | -10.937 | 7.67766E-28 | 8.08171E-27 | |
| AGAP006018 | AGAP006018 | nan | 82.4562232 | -2.358385416 | 0.2127 | -11.087 | 1.45341E-28 | 1.59031E-27 | |

| | | | | | | | | | |
|---|---|---|---|---|---|---|---|---|---|
| AGAP004106 | AGAP004106 | nan | 155.190438 | -2.355642662 | 0.2036 | -11.568 | 5.97053E-31 | 7.2485E-30 | |
| AGAP002457 | AGAP002457 | nan | 22.7808396 | -2.355336707 | 0.4475 | -5.2637 | 1.41202E-07 | 4.28979E-07 | |
| AGAP008582 | AGAP008582 | Alpha-mannosidase [Source:UniProtKB/TrEMBL;Acc:A0A1S4GYP5] | 44.6842384 | -2.352787645 | 0.251 | -9.3718 | 7.13126E-21 | 5.47098E-20 | |
| AGAP006728 | AGAP006728 | nan | 27.5540987 | -2.352718202 | 0.42 | -5.6017 | 2.12274E-08 | 6.99488E-08 | |
| AGAP000553 | white | protein white | 69.5525027 | -2.350714155 | 0.3416 | -6.882 | 5.90298E-12 | 2.64316E-11 | |
| AGAP009635 | AGAP009635 | nan | 15.4947577 | -2.344111236 | 0.536 | -4.3735 | 1.22252E-05 | 3.01028E-05 | |
| AGAP006724 | COEAE3G | carboxylesterase | 47.2679065 | -2.337800547 | 0.3693 | -6.33 | 2.45154E-10 | 9.74511E-10 | |
| AGAP001717 | AGAP001717 | nan | 21.0105193 | -2.337232131 | 0.4685 | -4.9889 | 6.07266E-07 | 1.73313E-06 | |
| AGAP028648 | AGAP028648 | nan | 510.247443 | -2.333370386 | 0.2169 | -10.758 | 5.43989E-27 | 5.52255E-26 | |
| AGAP010766 | AGAP010766 | nan | 17.7400371 | -2.332750466 | 0.4366 | -5.3425 | 9.16865E-08 | 2.83299E-07 | |
| AGAP003238 | AGAP003238 | N-myc downstream regulated | 1764.02464 | -2.331890497 | 0.21 | -11.103 | 1.21359E-28 | 1.33566E-27 | |
| AGAP007525 | AGAP007525 | coronin homolog | 483.016816 | -2.330480232 | 0.1292 | -18.034 | 1.06008E-72 | 4.66682E-71 | |
| AGAP009109 | AGAP009109 | nan | 656.568432 | -2.32919262 | 0.1171 | -19.894 | 4.56758E-88 | 2.92729E-86 | |
| AGAP005095 | AGAP005095 | actin beta/gamma 1 | 571.590763 | -2.32903806 | 0.2336 | -9.9721 | 2.01957E-23 | 1.7281E-22 | |
| AGAP009415 | AGAP009415 | lysophosphatidate acyltransferase | 191.787636 | -2.327522704 | 0.218 | -10.675 | 1.33206E-26 | 1.32797E-25 | |
| AGAP012052 | nan | nan | 10.0520965 | -2.326512702 | 0.5746 | -4.0491 | 5.14087E-05 | 0.000117754 | |
| AGAP009688 | AGAP009688 | Leucine-rich repeats and immunoglobulin-like domains protein 3 | 74.0276439 | -2.325611898 | 0.2811 | -8.2738 | 1.29807E-16 | 7.83919E-16 | |
| AGAP011948 | AGAP011948 | threonine 3-dehydrogenase | 947.615608 | -2.319896367 | 0.1697 | -13.673 | 1.4782E-42 | 2.7041E-41 | |
| AGAP009848 | nan | nan | 6.61977499 | -2.318913746 | 0.6367 | -3.6421 | 0.000270373 | 0.000562913 | |
| AGAP003030 | AGAP003030 | Pyruvate dehydrogenase E1 component subunit alpha [Source:UniPr | 1193.86372 | -2.313211468 | 0.1672 | -13.835 | 1.57042E-43 | 2.97087E-42 | |
| AGAP011523 | AGAP011523 | nan | 58.4287778 | -2.312625646 | 0.3869 | -5.9779 | 2.26011E-09 | 8.20836E-09 | |
| AGAP013439 | AGAP013439 | nan | 32.3701908 | -2.311477393 | 0.3233 | -7.1502 | 8.66827E-13 | 4.11197E-12 | |
| AGAP004119 | AGAP004119 | cGMP-specific 3', 5'-cyclic phosphodiesterase | 135.308647 | -2.311172394 | 0.1737 | -13.305 | 2.16381E-40 | 3.66642E-39 | |
| AGAP010508 | AGAP010508 | proton-coupled amino acid transporter | 82.2617448 | -2.306618937 | 0.2546 | -9.0608 | 1.29533E-19 | 9.29737E-19 | |
| AGAP012019 | AGAP012019 | nan | 257.54224 | -2.306162464 | 0.1967 | -11.723 | 9.77371E-32 | 1.21958E-30 | |
| AGAP012662 | AGAP012662 | omega-amidase | 919.738534 | -2.305791851 | 0.2436 | -9.467 | 2.87909E-21 | 2.25657E-20 | |
| AGAP002663 | AGAP002663 | nan | 76.0958659 | -2.301545809 | 0.2791 | -8.2454 | 1.64551E-16 | 9.89302E-16 | |
| AGAP011940 | AGAP011940 | nan | 23493.8308 | -2.300702471 | 0.1589 | -14.475 | 1.73554E-47 | 3.733E-46 | Granulocytes |
| AGAP028892 | LSU_rRNA_eukary | Eukaryotic large subunit ribosomal RNA [Source:RFAM;Acc:RF02543] | 383.839363 | -2.300185518 | 0.386 | -5.9591 | 2.53662E-09 | 9.13512E-09 | |
| AGAP005564 | AGAP005564 | CP2 domain-containing protein [Source:UniProtKB/TrEMBL;Acc:A7U | 11.7002051 | -2.299782375 | 0.4128 | -5.5709 | 2.53403E-08 | 8.27777E-08 | |
| AGAP001659 | AGAP001659 | hexamerin | 13.2905745 | -2.298331936 | 0.4735 | -4.8539 | 1.21073E-06 | 3.34497E-06 | |
| AGAP012954 | AGAP012954 | nan | 72.053172 | -2.294932598 | 0.296 | -7.7532 | 8.95762E-15 | 4.89784E-14 | |
| AGAP005370 | COEBE4C | carboxylesterase beta esterase | 16.5310695 | -2.291844587 | 0.4735 | -4.8406 | 1.2946E-06 | 3.56517E-06 | |
| AGAP008704 | AGAP008704 | Juvenile hormone-inducible protein [Source:UniProtKB/TrEMBL;Acc:. | 421.928565 | -2.28767662 | 0.2088 | -10.957 | 6.12395E-28 | 6.48244E-27 | |
| AGAP004318 | CLIPC3 | CLIP-domain serine protease | 1204.74386 | -2.287218806 | 0.2052 | -11.144 | 7.6366E-29 | 8.47402E-28 | |
| AGAP004262 | TO3 | takeout 3 | 29.1144885 | -2.286912991 | 0.3274 | -6.9845 | 2.85881E-12 | 1.30805E-11 | |
| AGAP006047 | CYP4J9 | cytochrome P450 | 102.844225 | -2.284419971 | 0.1975 | -11.565 | 6.18441E-31 | 7.4985E-30 | |
| AGAP011503 | AGAP011503 | nan | 642.360089 | -2.282940118 | 0.2205 | -10.351 | 4.13535E-25 | 3.85734E-24 | |
| AGAP012577 | AGAP012577 | nan | 3224.89105 | -2.281795555 | 0.2424 | -9.4125 | 4.84298E-21 | 3.73369E-20 | |
| AGAP003578 | AGAP003578 | aldehyde dehydrogenase (NAD+) | 2418.49943 | -2.281313782 | 0.0916 | -24.907 | 6.3135E-137 | 9.4412E-135 | Granulocytes |
| AGAP004762 | AGAP004762 | alpha-tocopherol transfer protein-like protein | 85.5034718 | -2.281133253 | 0.2467 | -9.2472 | 2.30543E-20 | 1.71967E-19 | |
| AGAP001470 | AGAP001470 | nan | 17971.8688 | -2.281092055 | 0.1994 | -11.439 | 2.68319E-30 | 3.20385E-29 | Granulocytes |
| AGAP003168 | AGAP003168 | Isocitrate dehydrogenase [NADP] [Source:UniProtKB/TrEMBL;Acc:Q7 | 6440.11392 | -2.275208578 | 0.1805 | -12.604 | 1.99719E-36 | 2.94453E-35 | |
| AGAP007334 | AGAP007334 | Tektin [Source:UniProtKB/TrEMBL;Acc:Q7QJ75] | 29.2657475 | -2.27501447 | 0.3021 | -7.5315 | 5.01499E-14 | 2.57472E-13 | |
| AGAP008889 | ABCG15 | ATP-binding cassette transporter (ABC transporter) family G member | 182.714194 | -2.265786564 | 0.2511 | -9.0235 | 1.82163E-19 | 1.29619E-18 | |
| AGAP003773 | AGAP003773 | nan | 703.878338 | -2.265195948 | 0.2323 | -9.7522 | 1.80474E-22 | 1.47847E-21 | |
| AGAP003301 | AGAP003301 | M-phase inducer phosphatase | 714.854198 | -2.260530545 | 0.2052 | -11.019 | 3.10743E-28 | 3.34191E-27 | |
| AGAP001116 | AGAP001116 | D-amino-acid oxidase | 916.292957 | -2.260425193 | 0.2041 | -11.077 | 1.63119E-28 | 1.7807E-27 | |
| AGAP012165 | nan | nan | 7.09811052 | -2.25837811 | 0.5328 | -4.2386 | 2.24916E-05 | 5.36302E-05 | |
| AGAP028764 | aga-mir-34 | nan | 9.22155097 | -2.25816415 | 0.4525 | -4.99 | 6.03724E-07 | 1.72406E-06 | |
| AGAP001657 | AGAP001657 | hexamerin | 23.7940774 | -2.25530474 | 0.3638 | -6.1989 | 5.685E-10 | 2.18606E-09 | |
| AGAP013540 | AGAP013540 | nan | 99.607894 | -2.255033994 | 0.2446 | -9.2187 | 3.0078E-20 | 2.23298E-19 | |
| AGAP007279 | AGAP007279 | nan | 293.691065 | -2.250852654 | 0.2131 | -10.561 | 4.51074E-26 | 4.381E-25 | |
| AGAP011984 | AGAP011984 | UDP-N-acetyl-alpha-D-galactosamine:polypeptide N-acetylgalactosar | 4104.57233 | -2.248194125 | 0.1976 | -11.378 | 5.38698E-30 | 6.33592E-29 | Granulocytes |
| AGAP008614 | AGAP008614 | nan | 116.770244 | -2.242769973 | 0.2087 | -10.747 | 6.12674E-27 | 6.21313E-26 | |
| AGAP001987 | AGAP001987 | nan | 91.8940082 | -2.241147168 | 0.2601 | -8.615 | 6.99526E-18 | 4.56387E-17 | |
| AGAP009115 | AGAP009115 | phosphatidylinositol phospholipase C, beta | 121.799069 | -2.240868083 | 0.1531 | -14.638 | 1.61471E-48 | 3.57934E-47 | |
| AGAP029185 | AGAP029185 | nan | 6259.02738 | -2.240571367 | 0.2081 | -10.768 | 4.85248E-27 | 4.94219E-26 | |
| AGAP007881 | AGAP007881 | Steroid dehydrogenase | 75.5187113 | -2.240117035 | 0.2082 | -10.76 | 5.31949E-27 | 5.40614E-26 | |
| AGAP000128 | AGAP000128 | MFS transporter, VNT family, synaptic vesicle glycoprotein 2 | 190.229327 | -2.239006767 | 0.2558 | -8.7539 | 2.0619E-18 | 1.37767E-17 | |
| AGAP003449 | AGAP003449 | rootletin | 132.804659 | -2.238462492 | 0.2285 | -9.7942 | 1.1924E-22 | 9.8887E-22 | |
| AGAP010640 | AGAP010640 | nan | 3865.77393 | -2.238179986 | 0.183 | -12.229 | 2.16425E-34 | 2.98527E-33 | |
| AGAP013036 | AGAP013036 | nan | 47.3192495 | -2.236496656 | 0.28 | -7.9881 | 1.37024E-15 | 7.85219E-15 | |
| AGAP009131 | AGAP009131 | nan | 27.7632898 | -2.234811024 | 0.3467 | -6.4465 | 1.14452E-10 | 4.65967E-10 | |
| AGAP000156 | AGAP000156 | nan | 28.4121895 | -2.231484108 | 0.315 | -7.0843 | 1.39724E-12 | 6.5133E-12 | |
| AGAP003328 | AGAP003328 | NADH dehydrogenase (ubiquinone) 1 alpha subcomplex 6 | 105.32323 | -2.230101589 | 0.3969 | -5.6186 | 1.92548E-08 | 6.36714E-08 | |
| AGAP007014 | AGAP007014 | Amino_oxidase domain-containing protein [Source:UniProtKB/TrEMI | 31.9757341 | -2.225944393 | 0.4408 | -5.05 | 4.41757E-07 | 1.27898E-06 | |
| AGAP013290 | LRIM27 | leucine-rich immune protein (Coil-less) | 401.682148 | -2.225568481 | 0.2364 | -9.4164 | 4.66818E-21 | 3.60188E-20 | |
| AGAP028564 | AGAP028564 | nan | 1831.46812 | -2.224442565 | 0.1833 | -12.138 | 6.6206E-34 | 8.97448E-33 | |
| AGAP012008 | AGAP012008 | nan | 207.211918 | -2.219825262 | 0.2262 | -9.8122 | 9.97317E-23 | 8.30011E-22 | |
| AGAP012053 | AGAP012053 | alpha,alpha-trehalase | 4513.24466 | -2.219554737 | 0.1444 | -15.372 | 2.53072E-53 | 6.53202E-52 | |
| AGAP009144 | AGAP009144 | nan | 360.441606 | -2.218211654 | 0.1419 | -15.635 | 4.18545E-55 | 1.13963E-53 | |
| AGAP004376 | AGAP004376 | aldose 1-epimerase | 454.806495 | -2.217002701 | 0.2047 | -10.829 | 2.51583E-27 | 2.59602E-26 | Oenocytoids |
| AGAP004306 | AGAP004306 | nan | 6.12836441 | -2.215525833 | 0.593 | -3.7362 | 0.000186819 | 0.000398737 | |
| AGAP006907 | AGAP006907 | Cat eye syndrome chromosome region, candidate 1a | 1136.7432 | -2.212906822 | 0.117 | -18.921 | 7.64914E-80 | 4.21418E-78 | |
| AGAP010473 | AGAP010473 | Pleckstrin-like proteiny domain-containing family G member 5 | 18.2736161 | -2.212106507 | 0.4326 | -5.1137 | 3.1588E-07 | 9.2997E-07 | |
| AGAP002324 | AGAP002324 | mitochondrial sodium/hydrogen exchanger NHA2 | 498.561816 | -2.210831099 | 0.2296 | -9.6301 | 5.96462E-22 | 4.8234E-21 | |
| AGAP000299 | AGAP000299 | nan | 255.253733 | -2.205293292 | 0.1263 | -17.455 | 3.17072E-68 | 1.20449E-66 | |
| AGAP012290 | AGAP012290 | nan | 131.239237 | -2.203350671 | 0.188 | -11.721 | 1.00038E-31 | 1.24664E-30 | |
| AGAP004118 | SCRAL1 | Class A Scavenger Receptor (SRCR domain) with Lysyl Oxidase domai | 14.0255715 | -2.202704126 | 0.442 | -4.9831 | 6.25627E-07 | 1.78391E-06 | |
| AGAP007790 | AGAP007790 | Sodium/potassium-transporting ATPase subunit beta [Source:UniPro | 5685.58683 | -2.202668102 | 0.106 | -20.78 | 6.54492E-96 | 4.85509E-94 | |
| AGAP001713 | AGAP001713 | stearoyl-CoA desaturase (delta-9 desaturase) | 24038.5132 | -2.200516903 | 0.2207 | -9.9689 | 2.08606E-23 | 1.78175E-22 | |
| AGAP003751 | CPAP1-C | cuticular protein | 107.406582 | -2.19811333 | 0.1952 | -11.258 | 2.10985E-29 | 2.39481E-28 | |
| AGAP012679 | AGAP012679 | nan | 23.4968382 | -2.197140641 | 0.4351 | -5.0497 | 4.42501E-07 | 1.28074E-06 | |
| AGAP012275 | AGAP012275 | nan | 113.518995 | -2.194052528 | 0.2382 | -9.2126 | 3.18275E-20 | 2.361E-19 | |
| AGAP007049 | AGAP007049 | nan | 3830.83849 | -2.193460503 | 0.2469 | -8.8836 | 6.47324E-19 | 4.44817E-18 | Megacytes |
| AGAP001283 | AGAP001283 | potassium inwardly-rectifying channel subfamily J | 22.1943654 | -2.193404679 | 0.3257 | -6.735 | 1.63985E-11 | 7.08994E-11 | |
| AGAP003441 | AGAP003441 | nan | 14.2669086 | -2.192601287 | 0.3697 | -5.9306 | 3.01748E-09 | 1.08131E-08 | |
| AGAP000097 | AGAP000097 | solute carrier family 25 | 2876.95234 | -2.189979922 | 0.2078 | -10.541 | 5.60827E-26 | 5.42459E-25 | |
| AGAP006756 | AGAP006756 | nan | 103.630912 | -2.189459005 | 0.1996 | -10.968 | 5.436E-28 | 5.76069E-27 | |
| AGAP027997 | LRIM5 | leucine-rich immune protein (Short) | 1455.95722 | -2.188307329 | 0.196 | -11.166 | 6.00569E-29 | 6.68789E-28 | |
| AGAP011643 | AGAP011643 | nan | 10.6236155 | -2.188271772 | 0.6175 | -3.544 | 0.00039417 | 0.000800836 | |
| AGAP028636 | AGAP028636 | nan | 1573.60888 | -2.185494737 | 0.2674 | -8.1742 | 2.97744E-16 | 1.7731E-15 | |
| AGAP010365 | AGAP010365 | nan | 8.4200103 | -2.181281288 | 0.4973 | -4.3866 | 1.15145E-05 | 2.84645E-05 | |

| | | | | | | | | | |
|---|---|---|---|---|---|---|---|---|---|
| AGAP000023 | AGAP000023 | nan | 18.0493725 | -2.178246081 | 0.5402 | -4.0324 | 5.52155E-05 | 0.000126075 | |
| AGAP002178 | AGAP002178 | homeobox protein homothorax | 44.5424978 | -2.174738692 | 0.2413 | -9.0134 | 1.99778E-19 | 1.41832E-18 | |
| AGAP013476 | AGAP013476 | nan | 14.4147151 | -2.173877245 | 0.5294 | -4.1062 | 4.02251E-05 | 9.31794E-05 | |
| AGAP006638 | AGAP006638 | Oatp58Dc | 12.6633594 | -2.170027724 | 0.5634 | -3.8517 | 0.000117306 | 0.000257068 | |
| AGAP000571 | CLIPC5 | CLIP-domain serine protease | 10.8570501 | -2.165583685 | 0.5998 | -3.6106 | 0.000305454 | 0.000630656 | |
| AGAP002377 | AGAP002377 | fatty-acid amide hydrolase 2 | 68.9084911 | -2.163506379 | 0.3091 | -6.9998 | 2.5633E-12 | 1.17684E-11 | |
| AGAP008311 | AGAP008311 | acylphosphatase | 7988.93375 | -2.161992961 | 0.2504 | -8.6338 | 5.93622E-18 | 3.89722E-17 | |
| AGAP002419 | CYP4D22 | cytochrome P450 | 1923.83179 | -2.160518349 | 0.0876 | -24.671 | 2.1876E-134 | 3.1707E-132 | |
| AGAP004730 | AGAP004730 | phospholipase A2, venom | 19.4809999 | -2.159578666 | 0.3953 | -5.463 | 4.68228E-08 | 1.49228E-07 | |
| AGAP001111 | AGAP001111 | nan | 471.415494 | -2.158151069 | 0.205 | -10.528 | 6.44399E-26 | 6.20745E-25 | |
| AGAP007400 | AGAP007400 | nan | 755.641709 | -2.15537651 | 0.1853 | -11.634 | 2.75536E-31 | 3.40213E-30 | |
| AGAP012320 | OBP25 | odorant-binding protein 25 | 77.2698096 | -2.154520754 | 0.2584 | -8.3378 | 7.56694E-17 | 4.64115E-16 | |
| AGAP000419 | nan | nan | 29.8116697 | -2.1542553 | 0.3554 | -6.0611 | 1.35205E-09 | 5.02273E-09 | |
| AGAP007843 | nan | nan | 2508.82842 | -2.146932903 | 0.1011 | -21.226 | 5.4709E-100 | 4.5212E-98 | |
| AGAP001264 | AGAP001264 | Plasma glutamate carboxypeptidase | 9682.13789 | -2.139077005 | 0.1423 | -15.037 | 4.20384E-51 | 1.00775E-49 | Granulocytes |
| AGAP028393 | ssu rRNA | nan | 17124.0786 | -2.137588739 | 0.2965 | -7.2087 | 5.65039E-13 | 2.71455E-12 | |
| AGAP000300 | AGAP000300 | Bifunctional pyrimidine biosynthesis protein (PyrABCN) | 3883.62761 | -2.13416748 | 0.0795 | -26.828 | 1.5227E-158 | 2.9887E-156 | |
| AGAP000881 | AGAP000881 | aldehyde dehydrogenase family 7 member A1 | 4456.34368 | -2.133098404 | 0.0716 | -29.786 | 5.9931E-195 | 1.882E-192 | Granulocytes |
| AGAP002620 | MISO | mating-induced stimulator of oogenesis MISO | 13.3059206 | -2.127162419 | 0.4494 | -4.733 | 2.21264E-06 | 5.95579E-06 | |
| AGAP001650 | AGAP001650 | cation transport regulator-like protein 2 | 1404.09983 | -2.124739553 | 0.2002 | -10.611 | 2.65818E-26 | 2.61406E-25 | |
| AGAP009285 | AGAP009285 | nan | 575.172146 | -2.12378252 | 0.1703 | -12.471 | 1.07776E-35 | 1.56209E-34 | |
| AGAP028491 | AQP2 | aquaporin | 2622.49654 | -2.123261447 | 0.2287 | -9.2834 | 1.64145E-20 | 1.23318E-19 | |
| AGAP003931 | AGAP003931 | nan | 89.0536391 | -2.12050679 | 0.2109 | -10.056 | 8.66738E-24 | 7.57471E-23 | |
| AGAP006009 | CPR30 | cuticular protein RR-1 family 30 | 17.0728277 | -2.119690384 | 0.3787 | -5.5974 | 2.176E-08 | 7.16284E-08 | |
| AGAP008113 | AGAP008113 | nan | 1100.17195 | -2.119589278 | 0.2139 | -9.9107 | 3.73952E-23 | 3.16817E-22 | |
| AGAP000834 | AGAP000834 | nan | 15.6386385 | -2.118096973 | 0.4258 | -4.9747 | 6.53363E-07 | 1.85962E-06 | |
| AGAP012900 | AGAP012900 | nan | 109.39396 | -2.11366951 | 0.2964 | -7.1317 | 9.91026E-13 | 4.6729E-12 | |
| AGAP028653 | AGAP028653 | nan | 16.7094924 | -2.113573078 | 0.4028 | -5.2473 | 1.54356E-07 | 4.67583E-07 | |
| AGAP012223 | nan | nan | 428.887679 | -2.112026931 | 0.1719 | -12.285 | 1.09582E-34 | 1.52267E-33 | |
| AGAP012034 | AGAP012034 | nan | 298.621047 | -2.110518824 | 0.2595 | -8.1338 | 4.15994E-16 | 2.45556E-15 | |
| AGAP004742 | AGAP004742 | Pyruvate carboxylase [Source:UniProtKB/TrEMBL;Acc:A7UUW7] | 9558.43452 | -2.106403767 | 0.1597 | -13.187 | 1.03766E-39 | 1.71205E-38 | AM granulocytes |
| AGAP010066 | AGAP010066 | nan | 558.200174 | -2.104753471 | 0.2562 | -8.2155 | 2.11199E-16 | 1.26491E-15 | |
| AGAP028470 | AGAP028470 | nan | 8.91645284 | -2.103492404 | 0.5561 | -3.7827 | 0.000155167 | 0.000334515 | |
| AGAP006821 | AGAP006821 | acetyl-CoA acyltransferase 2 | 7478.7622 | -2.098850418 | 0.1901 | -11.042 | 2.40893E-28 | 2.60557E-27 | |
| AGAP013078 | AGAP013078 | Plasma glutamate carboxypeptidase precursor | 71.6591333 | -2.097192675 | 0.2216 | -9.4634 | 2.97979E-21 | 2.32968E-20 | |
| AGAP001991 | nan | nan | 10.2003167 | -2.094781512 | 0.586 | -3.575 | 0.000350277 | 0.00071707 | |
| AGAP007315 | AGAP007315 | nan | 753.192575 | -2.09397512 | 0.2065 | -10.139 | 3.71041E-24 | 3.30083E-23 | |
| AGAP004700 | AGAP004700 | nan | 416.744629 | -2.093142641 | 0.2482 | -8.4324 | 3.38527E-17 | 2.12476E-16 | |
| AGAP000130 | nan | nan | 5.44505255 | -2.092342209 | 0.5987 | -3.4948 | 0.000474493 | 0.000953743 | |
| AGAP006278 | AGAP006278 | nan | 13581.1932 | -2.086522229 | 0.0655 | -31.854 | 1.1709E-222 | 5.2528E-220 | Granulocytes |
| AGAP008967 | AGAP008967 | calcium/calmodulin-dependent 3',5'-cyclic nucleotide phosphodieste | 107.393113 | -2.086261108 | 0.2924 | -7.1353 | 9.66081E-13 | 4.55756E-12 | |
| AGAP013192 | AGAP013192 | venom allergen | 17.0219767 | -2.085497543 | 0.3864 | -5.3973 | 6.76602E-08 | 2.11911E-07 | |
| AGAP005586 | AGAP005586 | nan | 11.1820093 | -2.085485398 | 0.5688 | -3.6664 | 0.000245949 | 0.00051571 | |
| AGAP012006 | AGAP012006 | Protein phosphatase 1 regulatory subunit 16A [Source:UniProtKB/Tr | 173.707427 | -2.084268535 | 0.1802 | -11.569 | 5.92785E-31 | 7.20598E-30 | |
| AGAP004890 | AGAP004890 | ribose-phosphate pyrophosphokinase | 8852.92536 | -2.083813712 | 0.158 | -13.191 | 9.92841E-40 | 1.64386E-38 | |
| AGAP012435 | AGAP012435 | nan | 15.6197647 | -2.08339812 | 0.3835 | -5.4323 | 5.56293E-08 | 1.7569E-07 | |
| AGAP000629 | AGAP000629 | nan | 23.8618019 | -2.083250723 | 0.3235 | -6.4401 | 1.19419E-10 | 4.85561E-10 | |
| AGAP008717 | AGAP008717 | hydroxymethylglutaryl-CoA lyase | 520.615658 | -2.082417529 | 0.1793 | -11.613 | 3.55499E-31 | 4.36088E-30 | |
| AGAP000090 | AGAP000090 | adenylate cyclase | 1948.60327 | -2.080296444 | 0.207 | -10.049 | 9.31266E-24 | 8.12248E-23 | |
| AGAP003167 | AGAP003167 | NAD(P) transhydrogenase | 1535.80979 | -2.080269361 | 0.1178 | -17.661 | 8.34232E-70 | 3.35868E-68 | |
| AGAP002992 | AGAP002992 | Carbonic anhydrase [Source:UniProtKB/TrEMBL;Acc:Q5TU56] | 1392.54205 | -2.079466438 | 0.1968 | -10.566 | 4.27788E-26 | 4.15913E-25 | |
| AGAP011518 | AGAP011518 | nan | 24.3270571 | -2.077310625 | 0.3884 | -5.349 | 8.84174E-08 | 2.73917E-07 | |
| AGAP008051 | SAP1 | sensory appendage protein 1 | 5638.92957 | -2.076512063 | 0.2105 | -9.8655 | 5.87431E-23 | 4.92804E-22 | |
| AGAP005507 | nSyb | synaptic vesicle-associated integral membrane protein | 79.0111676 | -2.074773617 | 0.181 | -11.465 | 1.98279E-30 | 2.37355E-29 | |
| AGAP006512 | nan | nan | 110.440291 | -2.074064219 | 0.2843 | -7.2943 | 3.00213E-13 | 1.46015E-12 | |
| AGAP009433 | AGAP009433 | arylformamidase | 907.898295 | -2.07304017 | 0.2937 | -7.0572 | 1.6988E-12 | 7.89173E-12 | |
| AGAP001936 | AGAP001936 | phosphatidylinositol phospholipase C, beta | 237.607847 | -2.072051026 | 0.1885 | -10.992 | 4.15409E-28 | 4.43212E-27 | |
| AGAP000129 | AGAP000129 | nan | 69.638292 | -2.071984736 | 0.1994 | -10.391 | 2.71112E-25 | 2.55415E-24 | |
| AGAP013443 | AGAP013443 | Plasma glutamate carboxypeptidase | 108.119815 | -2.071928878 | 0.1693 | -12.236 | 2.00547E-34 | 2.77031E-33 | |
| AGAP006132 | AGAP006132 | ganglioside-induced differentiation-associated-protein 1 | 111.53105 | -2.070212766 | 0.1752 | -11.814 | 3.29319E-32 | 4.19826E-31 | |
| AGAP002632 | AGAP002632 | nan | 10674.1222 | -2.068286741 | 0.2431 | -8.5067 | 1.7888E-17 | 1.1433E-16 | AM granulocytes |
| AGAP006731 | nan | nan | 8.65256326 | -2.066824188 | 0.5161 | -4.0043 | 6.21956E-05 | 0.000141294 | |
| AGAP003283 | AGAP003283 | atrial natriuretic peptide receptor A | 24.3287584 | -2.066317297 | 0.3369 | -6.1331 | 8.61977E-10 | 3.26525E-09 | |
| AGAP005865 | AGAP005865 | fumarylacetoacetase | 3370.03313 | -2.063049793 | 0.1288 | -16.022 | 8.92832E-58 | 2.67028E-56 | Oenocytoids |
| AGAP010367 | nan | nan | 140.777242 | -2.062172048 | 0.2229 | -9.2509 | 2.22549E-20 | 1.66268E-19 | |
| AGAP007636 | AGAP007636 | phosphatidate phosphatase LPIN | 2381.53633 | -2.062107291 | 0.1664 | -12.393 | 2.85874E-35 | 4.09304E-34 | |
| AGAP011317 | AGAP011317 | nan | 7508.68761 | -2.061480433 | 0.2178 | -9.4641 | 2.96155E-21 | 2.31926E-20 | |
| AGAP006908 | AGAP006908 | nan | 276.466364 | -2.060194817 | 0.2114 | -9.7458 | 1.92275E-22 | 1.57378E-21 | |
| AGAP002194 | AGAP002194 | 1D-myo-inositol-triphosphate 3-kinase | 1155.23353 | -2.060069468 | 0.0836 | -24.651 | 3.5995E-134 | 5.138E-132 | |
| AGAP011062 | AGAP011062 | GLOBIN domain-containing protein [Source:UniProtKB/TrEMBL;Acc:( | 35.0196878 | -2.059952927 | 0.2494 | -8.26 | 1.45708E-16 | 8.77695E-16 | |
| AGAP005183 | AGAP005183 | nan | 451.147516 | -2.057479912 | 0.2324 | -8.8523 | 8.57642E-19 | 5.84226E-18 | |
| AGAP000249 | AGAP000249 | mannosyl-glycoprotein endo-beta-N-acetylglucosaminidase | 24190.3946 | -2.05668358 | 0.1607 | -12.796 | 1.73325E-37 | 2.67687E-36 | Oenocytoids |
| AGAP009417 | AGAP009417 | nan | 7.27352313 | -2.052987884 | 0.5719 | -3.5901 | 0.0003306 | 0.000679001 | |
| AGAP004232 | AGAP004232 | pellino | 1462.74894 | -2.050918804 | 0.258 | -7.9499 | 1.86699E-15 | 1.06149E-14 | |
| AGAP009648 | AGAP009648 | 2-oxo-4-hydroxy-4-carboxy-5-ureidoimidazoline decarboxylase | 1149.88975 | -2.046919414 | 0.2289 | -8.9438 | 3.75945E-19 | 2.62743E-18 | |
| AGAP001690 | AGAP001690 | nan | 12.1928332 | -2.046676305 | 0.4324 | -4.7336 | 2.20626E-06 | 5.94031E-06 | |
| AGAP002378 | AGAP002378 | Adenylosuccinate lyase [Source:UniProtKB/TrEMBL;Acc:Q7QBZ6] | 3550.03286 | -2.046063833 | 0.2044 | -10.011 | 1.36952E-23 | 1.18045E-22 | |
| AGAP000768 | AGAP000768 | Septin 4 [Source:UniProtKB/TrEMBL;Acc:A0A1S4G9T6] | 54.721443 | -2.042753052 | 0.2127 | -9.6045 | 7.65258E-22 | 6.1778E-21 | |
| AGAP001257 | AGAP001257 | UTP--glucose-1-phosphate uridylyltransferase | 7912.21303 | -2.041670623 | 0.156 | -13.084 | 4.06898E-39 | 6.63215E-38 | |
| AGAP003239 | AGAP003239 | nan | 593.849156 | -2.041349275 | 0.2561 | -7.9721 | 1.55975E-15 | 8.89495E-15 | |
| AGAP000806 | AGAP000806 | Angiopoietin-like 1 | 4950.15451 | -2.039105955 | 0.0944 | -21.601 | 1.7461E-103 | 1.5971E-101 | |
| AGAP008138 | AGAP008138 | nan | 8.29291267 | -2.038910263 | 0.5414 | -3.7663 | 0.000165684 | 0.000355723 | |
| AGAP007757 | AGAP007757 | Gustatory receptor [Source:UniProtKB/TrEMBL;Acc:Q7PIY0] | 42.4383324 | -2.038875842 | 0.2814 | -7.2444 | 4.34501E-13 | 2.10027E-12 | |
| AGAP002068 | AGAP002068 | nan | 111.485108 | -2.034382171 | 0.2654 | -7.6667 | 1.76426E-14 | 9.3693E-14 | |
| AGAP009786 | AGAP009786 | phosphoribosylamine--glycine ligase / phosphoribosylglycinamide fo | 3517.44325 | -2.034314259 | 0.1794 | -11.339 | 8.4086E-30 | 9.75583E-29 | |
| AGAP000481 | AGAP000481 | cyanogenic beta-glucosidase | 308.242708 | -2.033760652 | 0.1819 | -11.183 | 4.96101E-29 | 5.54421E-28 | |
| AGAP007304 | AGAP007304 | nan | 12.5422352 | -2.030801082 | 0.459 | -4.4243 | 9.67432E-06 | 2.41499E-05 | |
| AGAP001065 | AGAP001065 | glycine hydroxymethyltransferase | 6907.60554 | -2.030513674 | 0.1874 | -10.835 | 2.35607E-27 | 2.43383E-26 | |
| AGAP010733 | AGAP010733 | Vanin-like protein 1 | 976.750068 | -2.023907092 | 0.1783 | -11.35 | 7.40834E-30 | 8.60592E-29 | |
| AGAP012557 | AGAP012557 | nan | 141.642617 | -2.020568992 | 0.1793 | -11.269 | 1.87319E-29 | 2.12875E-28 | |
| AGAP000366 | AGAP000366 | nan | 21.0535763 | -2.018835092 | 0.3807 | -5.3036 | 1.1351E-07 | 3.47765E-07 | |
| AGAP000198 | Cht5-5 | Chitinase [Source:UniProtKB/TrEMBL;Acc:A0A1S4G846] | 8.17446088 | -2.017481331 | 0.5173 | -3.9001 | 9.61601E-05 | 0.00021351 | |
| AGAP003586 | AGAP003586 | Phosphate carrier, mitochondrial | 5065.702 | -2.017271645 | 0.2432 | -8.2937 | 1.09798E-16 | 6.6693E-16 | |
| AGAP003141 | AGAP003141 | Insulin-related peptide binding protein | 349.295186 | -2.01487268 | 0.1762 | -11.436 | 2.77389E-30 | 3.30377E-29 | |

| | | | | | | | | |
|---|---|---|---|---|---|---|---|---|
| AGAP010042 | AGAP010042 | solute carrier organic anion transporter family member | 21.0389471 | -2.013165455 | 0.3254 | -6.1858 | 6.17889E-10 | 2.36631E-09 |
| AGAP001515 | AGAP001515 | nan | 423.014864 | -2.012263574 | 0.2512 | -8.0115 | 1.13334E-15 | 6.52239E-15 |
| AGAP001200 | AGAP001200 | glycogen debranching enzyme | 2019.46101 | -2.008162939 | 0.1894 | -10.605 | 2.82763E-26 | 2.77338E-25 |
| AGAP004793 | AGAP004793 | ornithine--oxo-acid transaminase | 92.9537762 | -2.006348426 | 0.2092 | -9.5893 | 8.86788E-22 | 7.14054E-21 |
| AGAP005662 | AGAP005662 | acyl-CoA dehydrogenase | 10248.125 | -2.004762998 | 0.177 | -11.326 | 9.73147E-30 | 1.12079E-28 |
| AGAP006513 | nan | nan | 210.228167 | -2.004080989 | 0.1764 | -11.364 | 6.3446E-30 | 7.39758E-29 |
| AGAP002429 | CYP314A1 | cytochrome P450 | 38.4907408 | -2.003366386 | 0.2933 | -6.8304 | 8.46568E-12 | 3.74437E-11 |
| AGAP001550 | AGAP001550 | sodium-coupled monocarboxylate transporter 1 | 198.112491 | -2.001364547 | 0.2249 | -8.8995 | 5.61224E-19 | 3.87631E-18 |
