## Supplemental table S3 for "Hemocyte Differentiation to the Megacyte Lineage Enhances Mosquito Immunity Against *Plasmodium*"

| Toll pathway upregulated genes in Cactus-silenced hemocytes | | | | | | | | |
|---|---|---|---|---|---|---|---|---|
| **Accesion Number** | **Name** | **Description** | **baseMean** | **log2FoldChange** | **lfcSE** | **stat** | **pvalue** | **padj** |
| AGAP002842 | AGAP002842 | CLIPD1 protein | 908.0228542 | 9.35826901 | 0.461470828 | 20.27922121 | 1.96213E-91 | 1.31101E-89 |
| AGAP001002 | AGAP001002 | Toll protein | 116127.845 | 6.177628682 | 0.268236101 | 23.03056397 | 2.3037E-117 | 2.7825E-115 |
| AGAP010833 | CLIPB14 | CLIP-domain serine protease | 350.1728028 | 5.609302809 | 0.696698509 | 8.051262835 | 8.19441E-16 | 4.75953E-15 |
| AGAP010816 | TEP3 | thioester-containing protein 3 | 62660.7236 | 4.153502512 | 0.220728572 | 18.81724002 | 5.45532E-79 | 2.90365E-77 |
| AGAP001004 | TOLL1A | TOLL-like receptor 1A | 757.4906654 | 4.105939711 | 0.211676876 | 19.39720481 | 8.1483E-84 | 4.95259E-82 |
| AGAP001376 | SRPN17 | serine protease inhibitor (serpin) 17 | 2134.989079 | 3.685525524 | 0.244477096 | 15.07513619 | 2.36029E-51 | 5.67252E-50 |
| AGAP003139 | SRPN9 | serine protease inhibitor (serpin) 9 | 3098.83565 | 3.528515234 | 0.1217241 | 28.98781114 | 9.3726E-185 | 2.6757E-182 |
| AGAP006343 | PGRPS2 | peptidoglycan recognition protein (short) | 19.36251327 | 3.509206816 | 0.504347815 | 6.957910224 | 3.45357E-12 | 1.57179E-11 |
| AGAP001648 | CLIPB17 | CLIP-domain serine protease | 9264.846222 | 3.29031506 | 0.30899714 | 10.64836735 | 1.7745E-26 | 1.7542E-25 |
| AGAP003249 | CLIPB3 | CLIP-domain serine protease | 699.1854621 | 3.050823047 | 0.238996434 | 12.76514047 | 2.56687E-37 | 3.93211E-36 |
| AGAP008366 | TEP2 | thioester-containing protein 2 | 2622.051057 | 2.943343731 | 0.0858671 | 34.27789849 | 1.6754E-257 | 1.5784E-254 |
| AGAP011294 | DEF1 | defensin anti-microbial peptide | 9881.820375 | 2.922360646 | 0.52534384 | 5.562758 | 2.65544E-08 | 8.64739E-08 |
| AGAP009212 | SRPN6 | serine protease inhibitor (serpin) 6 | 4770.187149 | 2.896154957 | 0.163756954 | 17.68569143 | 5.40539E-70 | 2.20451E-68 |
| AGAP006342 | PGRPS3 | peptidoglycan recognition protein (short) | 17.74977137 | 2.729041906 | 0.702853207 | 3.882804945 | 0.000103258 | 0.00022841 |
| AGAP005246 | SRPN10 | serine protease inhibitor (serpin) 10 | 11070.86802 | 2.628637134 | 0.086237398 | 30.481406 | 4.5966E-204 | 1.6039E-201 |
| AGAP007036 | APL1A | Anopheles Plasmodium-responsive Leucine-Rich Repeat 1A | 1456.477362 | 2.623653516 | 0.264571135 | 9.916627985 | 3.52463E-23 | 2.9888E-22 |
| AGAP013027 | AGAP013027 | Protein toll | 9828.824707 | 2.611676022 | 0.210977531 | 12.3789297 | 3.39845E-35 | 4.84369E-34 |
| AGAP003247 | CLIPB19 | CLIP-domain serine protease | 1759.954841 | 2.570395939 | 0.274299017 | 9.370780717 | 7.1998E-21 | 5.51458E-20 |
| AGAP002270 | CLIPB7 | CLIP-domain serine protease | 659.568654 | 2.507618185 | 0.252318085 | 9.938321234 | 2.83566E-23 | 2.41543E-22 |
| AGAP007035 | APL1B | Anopheles Plasmodium-responsive Leucine-Rich Repeat 1B | 1019.493622 | 2.443761697 | 0.276147234 | 8.849488238 | 8.79215E-19 | 5.98057E-18 |
| AGAP013184 | CLIPB36 | CLIP-domain serine protease | 1066.00722 | 2.426717555 | 0.273967982 | 8.857668471 | 8.17053E-19 | 5.57787E-18 |
| AGAP007033 | APL1C | Anopheles Plasmodium-responsive Leucine-Rich Repeat 1C | 11735.16759 | 2.407308159 | 0.261412809 | 9.208837799 | 3.29675E-20 | 2.4398E-19 |
| AGAP008835 | CLIPC1 | CLIP-domain serine protease | 35.05362748 | 2.338817255 | 0.339891883 | 6.881062391 | 5.94078E-12 | 2.65882E-11 |
| AGAP002811 | CLIPD4 | CLIP-domain serine protease [Source:UniProtKB/TrEMBL;Acc:A0A1S4G | 16.51541341 | 2.263256704 | 0.377785221 | 5.990855598 | 2.0874E-09 | 7.61634E-09 |
| AGAP011790 | CLIPA2 | CLIP-domain serine protease | 3495.717731 | 2.214259384 | 0.266506231 | 8.308471353 | 9.69428E-17 | 5.90749E-16 |
| AGAP009221 | SRPN5 | serine protease inhibitor (serpin) 5 | 2214.441769 | 2.047353148 | 0.174240138 | 11.75018094 | 7.04678E-32 | 8.83991E-31 |
| AGAP003689 | CLIPC7 | CLIP-domain serine protease [Source:UniProtKB/TrEMBL;Acc:A0A1S4G | 1211.441982 | 2.04098826 | 0.356864366 | 5.719226844 | 1.0701E-08 | 3.62771E-08 |
